# Dietaryindex: A User-Friendly and Versatile R Package for Standardizing Dietary Pattern Analysis in Epidemiological and Clinical Studies

**DOI:** 10.1101/2023.08.07.548466

**Authors:** Jiada James Zhan, Rebecca A Hodge, Anne L Dunlop, Matthew M. Lee, Linh Bui, Donghai Liang, Erin P Ferranti

## Abstract

**Background:** Few standardized and open-source tools exist for calculating dietary pattern indexes from dietary intake data in epidemiological and clinical studies. Miscalculations of dietary indexes, with suspected erroneous findings, are occasionally noted in the literature.

**Objective:** The primary aim is to develop and validate dietaryindex, a user-friendly and versatile R package that standardizes the calculation of dietary indexes.

**Methods:** Dietaryindex utilizes a two-step process: an initial calculation of serving size for each food and nutrient category, followed by the calculation of individual dietary indexes. It includes generic functions that accept any preprocessed serving sizes of food groups and nutrients, with the standard serving sizes defined according to the methodologies used in well-known prospective cohort studies. For ease of use, dietaryindex also offers one-step functions that directly reference common datasets and tools, including the National Health and Nutrition Examination Survey (NHANES) and Block Food Frequency Questionnaire, eliminating the need for data preprocessing. At least two independent researchers validated the serving size definitions and scoring algorithms of dietaryindex.

**Results:** Dietaryindex can calculate multiple dietary indexes of high interest in research, including Healthy Eating Index (HEI) - 2020, Alternative Healthy Eating Index 2010, Dietary Approaches to Stop Hypertension Index, Alternate Mediterranean Diet Score, Dietary Inflammatory Index, American Cancer Society 2020 dietary index, and Planetary Health Diet Index from the EAT-Lancet Commission. In our validation process, dietaryindex demonstrated full accuracy (100%) in all generic functions with two-decimal rounding precision in comparison to hand-calculated results. Similarly, using NHANES 2017-2018 data and ASA24 and DHQ3 example data, the HEI2015 outputs from dietaryindex aligned (99.95% - 100%) with results using the SAS codes from the National Cancer Institute.

**Conclusions:** Dietaryindex is a user-friendly, versatile, and validated informatics tool for standardized dietary index calculations. We have open-sourced all the validation files and codes with detailed tutorials on GitHub (https://github.com/jamesjiadazhan/dietaryindex).

## Introduction

The role of dietary patterns in understanding the relationship between lifestyle and human diseases is well-established in the fields of nutrition, medicine, and epidemiology. Unhealthy dietary patterns are associated with a higher risk of various chronic diseases, including type 2 diabetes,^1^ cardiovascular disease,^2^ cancer,^3^ hypertension,^4^ and obesity.^5^ Given that long-term dietary interventions are often either infeasible or ethically problematic in human studies, dietary pattern indexes (dietary indexes) are often used as an alternative strategy to quantify the individuals’ dietary intake and to assess their relationship with chronic diseases in prospective cohort studies and large-scale cross-sectional studies, including the National Health and Nutrition Examination Survey (NHANES).^6^

However, the comparison of dietary intake findings across diverse epidemiological studies poses a challenge without standardized methods for calculating dietary indexes. This is particularly true for studies employing quintile-based dietary indexes or those applying heterogenous serving size definitions for foods and nutrients. The lack of standardized methods may deter researchers from designing clinical studies based on epidemiological findings and comparing clinical results with epidemiological results precisely. Therefore, it is essential to apply dietary indexes in a standardized manner across epidemiological and clinical studies, in such a way that their results are comparable. This standardization will empower researchers to compare results across different research settings and thereby enhance the translation of research findings into practical clinical applications.

Yet obtaining standardized dietary index computation can be complex and prone to inaccuracies. Miscalculations of dietary indexes can lead to erroneous results and have been occasionally noted in the literature, particularly for complex dietary indexes such as the Dietary Inflammatory Index (DII).^7^ Moreover, although there are few tools for calculating Healthy Eating Index-2015,^8,9^ no standardized and open-source tools exist for calculating dietary indexes from dietary data for both epidemiological and clinical studies using various nutritional assessment tools. To fill this knowledge gap, we developed ‘dietaryindex’, an R package that provides user-friendly and streamlined methods to standardize the compilation of dietary intake data into index-based dietary patterns. To validate this package, we compared dietaryindex-generated results with hand-calculated results using simulated datasets. The ‘dietaryindex’ package could play an important role in assessing adherence to dietary patterns in epidemiologic and clinical studies and facilitating precision nutrition.

## Methods

### Computation process

The dietaryindex package uses a structured two-step computation process. The first step aligns units of various food groups and nutrients to the standard units proposed by dietaryindex, by referring to methodologies from well-known prospective cohort studies (Supplementary Material 1).^10–12^ This step ensures the comparability of serving sizes across various dietary assessment tools. The second step inputs the standardized serving sizes into dietary index scoring algorithms, yielding the final dietary index total score and any dietary index component scores (Supplementary Material 1).

The package’s flexible structure allows for the input of preprocessed serving sizes in two-step function computations. For instance, to calculate the Alternate Healthy Eating Index (AHEI) score, researchers only need to convert their data to the relevant serving sizes, as per the ‘dietaryindex’ guidelines. This cleaned data can then be input into a function to generate the standard AHEI index.

For convenience, dietaryindex also offers one-step functions requiring no data preprocessing. It directly processes common datasets and tools such as NHANES, Automated Self-Administered 24-hour Dietary Assessment Tool (ASA24), Diet History Questionnaire III (DHQ3), and Block Food Frequency Questionnaire (Block FFQ). Users can input raw data directly, and dietaryindex will output the dietary index scores, removing the need for manual data cleaning. This user-friendly feature enables nearly instantaneous computation of various dietary indexes by eliminating the need for any manual data cleaning for common nutrition-related datasets and tools, which also simultaneously eliminates any misclassification of serving sizes that might arise in the manual cleaning process.

The package also uses the tibble data structure in R for quick calculations, even with large datasets. For instance, a typical NHANES dataset, containing nearly 2 million rows and 138 columns, can be processed in seconds. This swift computation reduces the time needed for data processing and analysis in large-scale dietary epidemiological and clinical studies.

### Validation

We prioritized accuracy and precision in the development of dietaryindex. To ensure this, we engaged at least two independent researchers in an extensive validation process. The validation process has 3 parts:

1. Zhan, Hodge, and Lee reviewed and confirmed the dietaryindex’s serving size definitions for food groups and nutrients by documenting them clearly in Supplementary Material 1 and comparing them to the methodologies used in large, well-known prospective cohort studies.^10–12^ This stage served as an internal validation.
2. To assess the accuracy and precision of the dietaryindex’s scoring algorithms, Zhan and Hodge developed simulated serving size data (sample size ranges from 10 to 26). They manually generated various serving sizes within the minimum and maximum index range and created CSV files for each dietary index. They then compared hand-calculated results with the results generated by dietaryindex. This comparison included all dietary indexes available in the dietaryindex package. Both hand-calculated and dietaryindex-calculated results were rounded to the nearest two decimal places to assess for an exact match. This stage served as internal validation.
3. For external validation, Zhan and Hodge compared HEI2015 scores generated from dietaryindex and the National Cancer Institute (NCI)-published SAS programs for NHANES and ASA24, using data from NHANES 2017-2018 (n=7122) and our example data for ASA24 (n=21) and DHQ3 (n=23) (Supplementary Material 2).^8^ The rationale is that the HEI2015 SAS codes from NCI are the only dietary index codes from a well-established institution that we can directly validate our calculation with their calculation as the gold standard. DHQ3 has internally calculated HEI2015 scores from NCI and was used to compare with the dietaryindex’s result. The NHANES and ASA24 validations followed the 2-decimal rounding and exact matching process, the same as the first validation section. The DHQ3 validation followed a different procedure in that it subtracts NCI internal-calculated results from the dietaryindex’s results, takes the absolute value of the difference, and checks if all differences are not greater than our defined maximum tolerance, 0.5. This procedure was adopted since the DHQ3 HEI2015 scores used different rounding rules compared to the rounding rules used in NCI SAS codes for NHANES and ASA24.

For transparency and verification of the package’s robustness, all validation codes and documents are openly accessible (Supplementary Material 2 and https://github.com/jamesjiadazhan/dietaryindex/tree/main/Validation%20file%20for%20publication).

## Results

### Dietary index options

Dietaryindex can calculate multiple dietary pattern indexes of high interest in epidemiologic, public health, nutrition, and clinical research studies, including Healthy Eating Index-2020 and Healthy Eating Index-Toddlers-2020 (HEI2020),^13,14^ Healthy Eating Index-2015 (HEI2015),^15^ Alternative Healthy Eating Index 2010 (AHEI),^10^ Alternate Healthy Eating Index, slightly modified for pregnancy (AHEI-P),^16^ Dietary Approaches to Stop Hypertension Index (DASH, quintile-based),^11^ DASH Index in serving sizes adapted from the DASH trial (DASHI),^4,17^ Alternate Mediterranean Diet Score (aMED, median-based),^12^ MED Index in serving sizes from the PREDIMED trial (MEDI),^18^ Dietary Inflammatory Index (DII),^19^ American Cancer Society 2020 dietary index (ACS 2020),^20^ and Planetary Health Diet Index based on the EAT-Lancet Commission (PHDI).^21^ All of these dietary indexes can be calculated by using any dietary assessment tool with preprocessed data that have standardized serving sizes for each food group and nutrient (Table 1). Additionally, many of these dietary indexes can be calculated using common nutrition datasets and tools, such as NHANES, ASA24, DHQ3, and Block FFQ without any data pre-processing (Table 1). To facilitate the NHANES-related functions, NHANES nutrition datasets from 2005 to 2018 have been pre-compiled and included in dietaryindex, and the NHANES functions are also versatile, allowing users to enter either first-day data, second-day data, or combine both, and return results accordingly.

**Table 1.** The list of dietary pattern indexes that the dietaryindex package can calculate.

| Dietary pattern index | Generic functions for any dietary assessments | Specific 1-step functions for common nutrition datasets and tools |  |  |  |
| --- | --- | --- | --- | --- | --- |
|  |  | NHANES | ASA24 | DHQ3 | BLOCK FFQ |
| HEI2020 | HEI2020() | HEI2020_NHANES_FPED() |  |  |  |
| HEI2015 | HEI2015() | HEI2015_NHANES_FPED() | HEI2015_ASA24() | HEI2015_DHQ3() | HEI2015_BLOCK() |
| AHEI | AHEI() | AHEI_NHANES_FPED() | AHEI_F_ASA24()<br>for female only<br>AHEI_M_ASA24()<br>for male only | AHEI_DHQ3() | AHEI_BLOCK() |
| AHEIP | AHEIP() |  |  |  | AHEIP_BLOCK() |
| DASH | DASH() | DASH_NHANES_FPED() | DASH_ASA24() | DASH_DHQ3() | DASH_BLOCK() |
| DASHI | DASHI() | DASHI_NHANES_FPED() |  |  |  |
| aMED | aMED() | MED_NHANES_FPED() | MED_ASA24() | MED_DHQ3() | MED_BLOCK() |
| MEDI | MEDI()<br>0/1 scoring criteria<br>MEDI_V2()<br>5 points scoring<br>criteria | MEDI_NHANES_FPED()<br>0/1 scoring criteria |  |  |  |
| DII | DII() | DII_NHANES_FPED() | DII_ASA24() |  | DII_BLOCK() |
| ACS<br>2020 | ACS2020_V1()<br>ACS2020_V2() |  |  |  |  |
| PHDI | PHDI() |  |  |  |  |

### Validation

All dietaryindex-calculated results exhibited a full 100% accuracy when subjected to 2-decimal rounding in comparison with the hand-calculated results (Figure 1 and Supplementary Material 3). Moreover, when utilizing the NHANES 2017-2018 data and our ASA24 example data, the HEI2015 outputs from the dietaryindex package aligned (99.95% -100%) with the results derived from the SAS codes in NHANES and ASA24, again featuring a 2-decimal rounding precision (Figure 2 and 3). Compared with the internal-calculated HEI2015 results in our DHQ3 example data, the HEI2015 outputs from the dietaryindex package also demonstrated a full 100% accuracy, with all the differences between the two results not greater than 0.5 (Figure 4).

**Figure 1.**
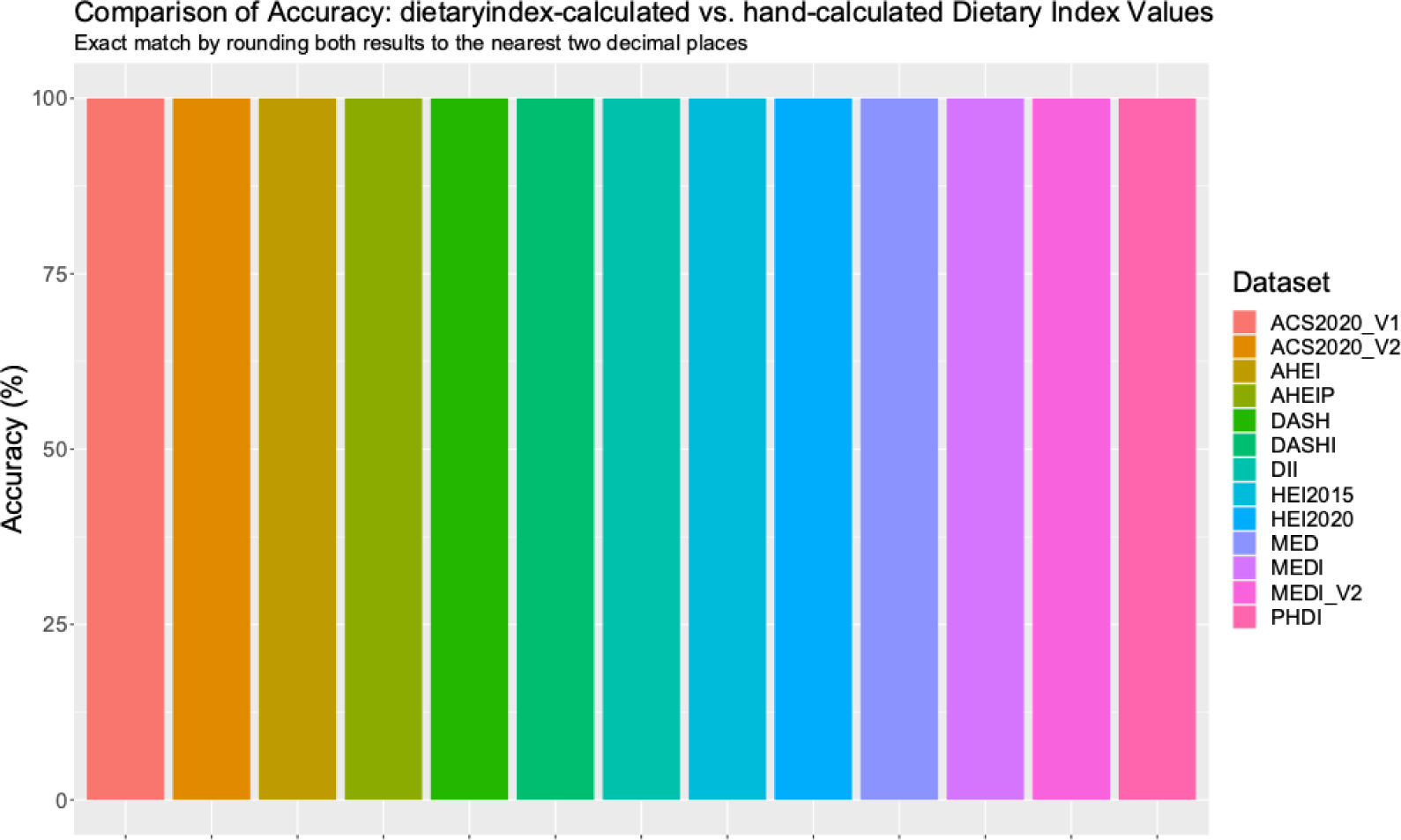
Comparison of Accuracy: dietaryindex-calculated vs. hand-calculated Dietary Index Values using the simulation datasets (sample sizes range from 10 to 26).

**Figure 2.**
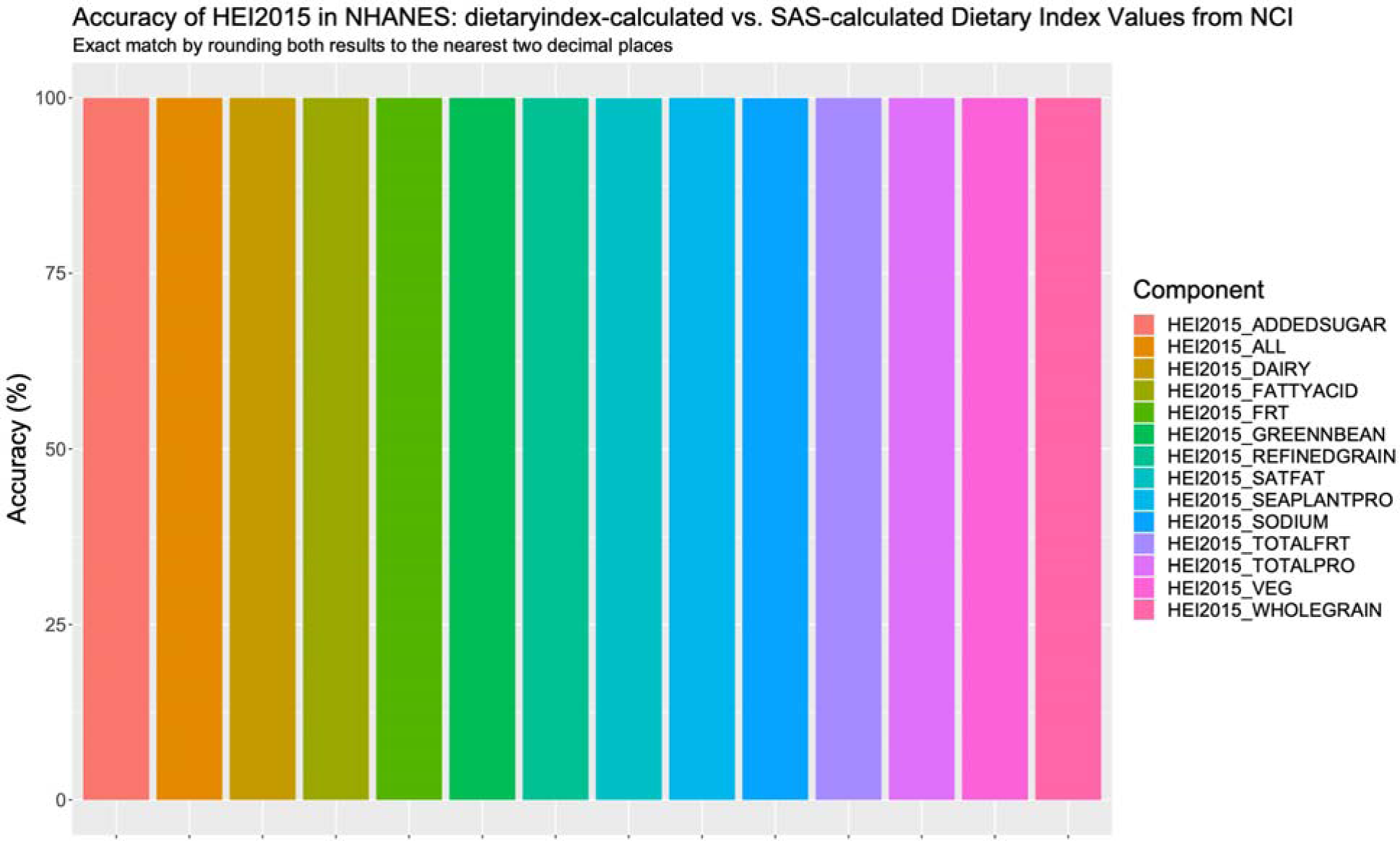
Accuracy of HEI2015 in NHANES: dietaryindex-calculated vs. SAS-calculated results from National Cancer Institute using the NHANES 2017-2018 data (n=7122).

**Figure 3.**
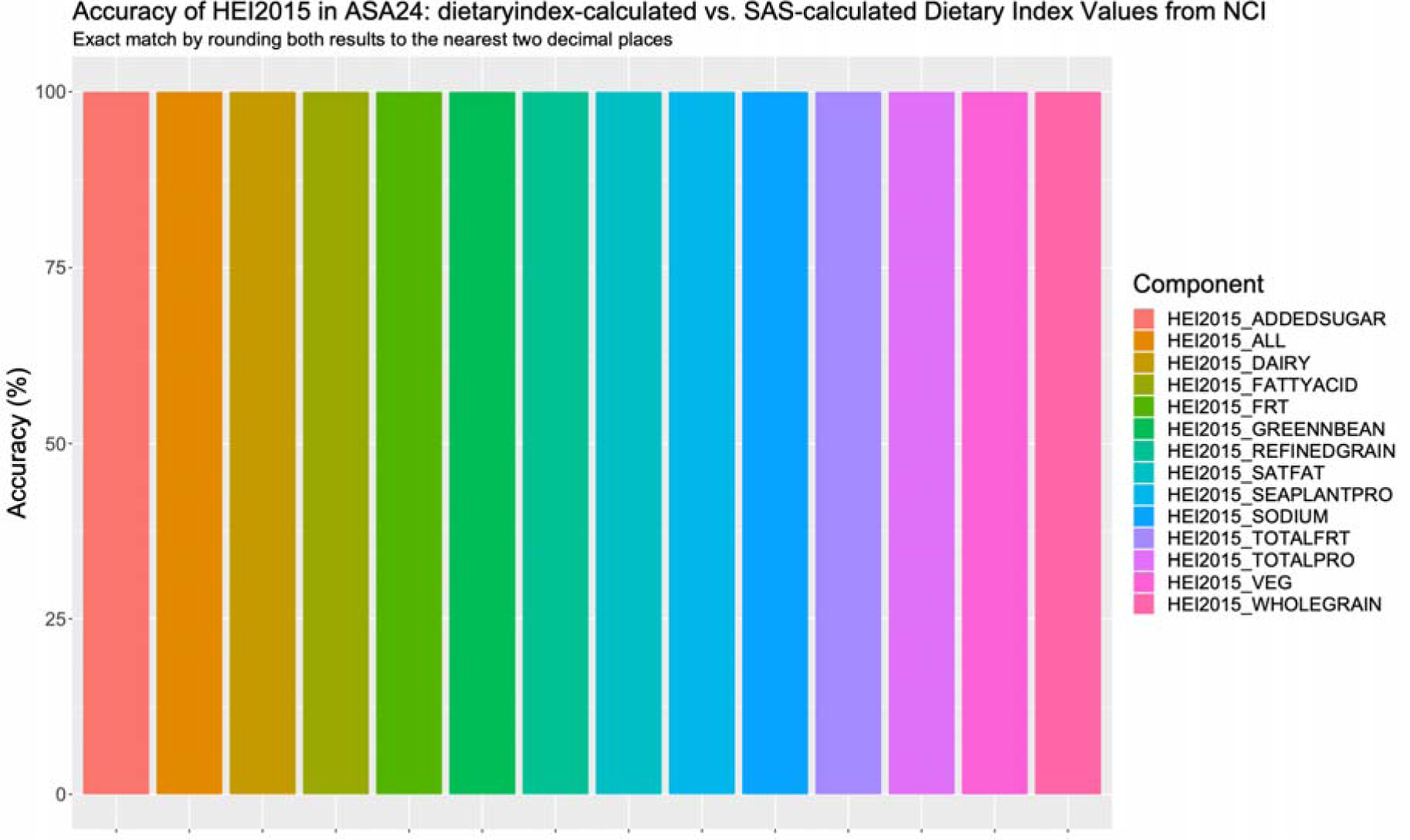
Accuracy of HEI2015 in ASA24: dietaryindex-calculated vs. SAS-calculated results from National Cancer Institute using the ASA24 example data (n=21).

**Figure 4.**
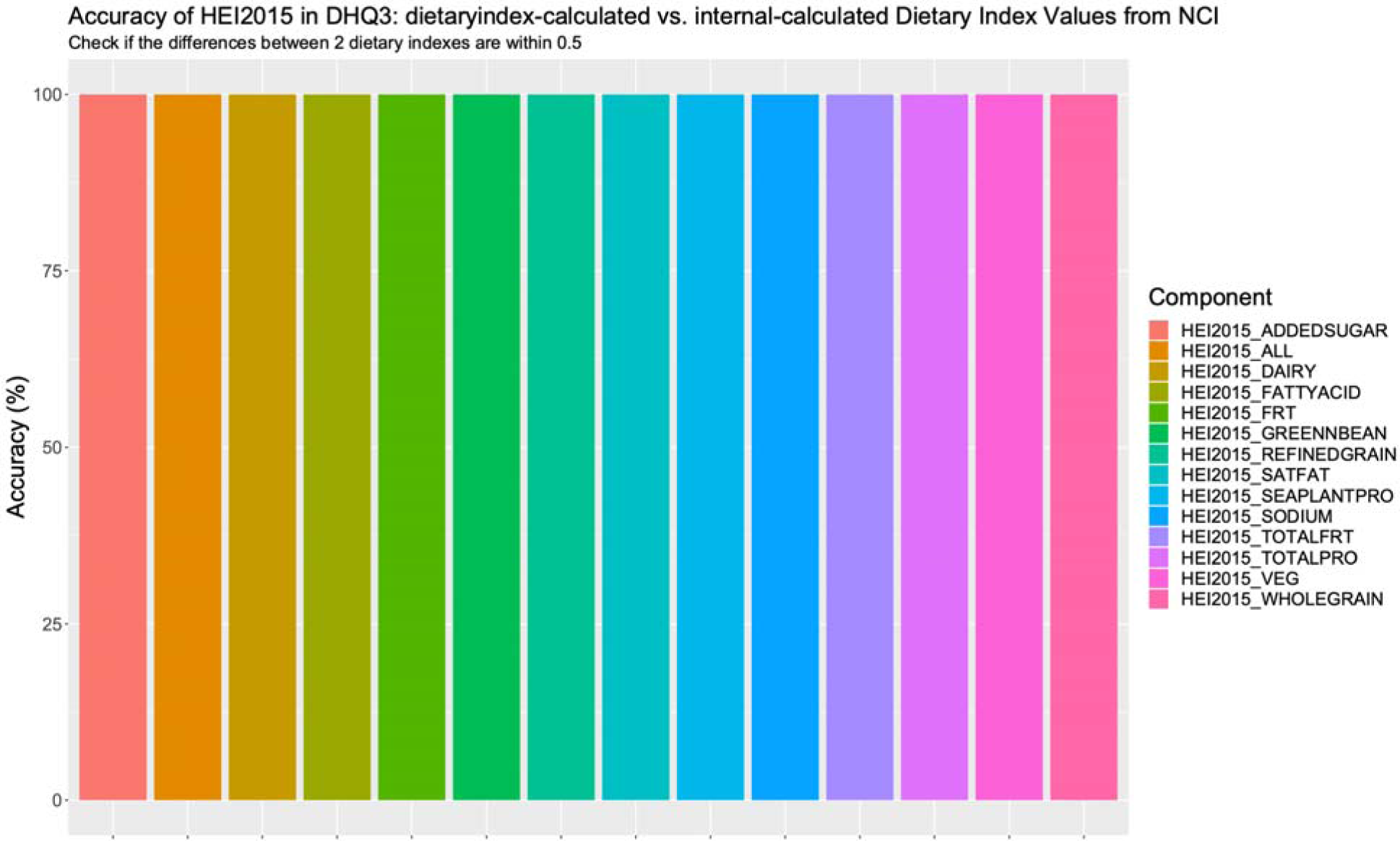
Accuracy of HEI2015 in DHQ3: dietaryindex-calculated vs. internal-calculated results from National Cancer Institute using the DHQ3 example data (n=23).

### Case studies

In this section, to demonstrate that our package can support a wide range of research questions with a concise programming structure, we delineate three distinct methodological exemplifications of using the dietaryindex package in epidemiological and clinical research:

1. Case study 1: A comparative analysis of dietary intake according to the DASHI and MEDI dietary intakes for participants enrolled in clinical trials (i.e., The Dietary Approaches to Stop Hypertension [DASH] trial and Prevención con Dieta Mediterránea [PREDIMED] trial) and a population-based epidemiological study (i.e., NHANES) from 2017-2018.
2. Case study 2: A time series of cross-sectional computation of the HEI2020 in the NHANES dataset spanning 2005 to 2018, stratifying into toddler and non-toddler populations.
3. Case study 3: A comprehensive calculation of multiple dietary indexes—HEI2020, AHEI, DASH, DASHI, MED, MEDI, DII—within a single year (2017-2018), leveraging data from the NHANES study.

#### Case study 1

*A comparative analysis of dietary intake according to the DASHI and MEDI dietary intakes for participants enrolled in clinical trials (i*.*e*., *The Dietary Approaches to Stop Hypertension [DASH] trial and Prevención con Dieta Mediterránea [PREDIMED] trial) and a population-based epidemiological study (i*.*e*., *NHANES) from 2017-2018*.

Here, we present an example of a comparative analysis of DASHI and MEDI dietary indexes among participants enrolled in the DASH and PREDIMED trials and in the NHANES

2017-2018 sample. The rationale for using DASHI and MEDI, instead of DASH and aMED, is that DASHI and MEDI are serving size-based dietary indexes that can be used to compare results in different populations, whereas DASH and aMED are quintile-based and median-based dietary indexes that are incomparable across studies with different populations.

We first extracted the raw data from the DASH trial, PREDIMED trial, and NHANES and stored them in the dietaryindex package. 2-day dietary recall data were used in the NHANES 2017-2018 cycle. Then, we used these data as inputs to calculate the mean total score for DASHI in the DASH trial and NHANES, as well as for MEDI in the PREDIMED trial and NHANES using the following dietaryindex’s functions: DASHI(), MEDI(), DASHI_NHANES_FPED(), and MEDI_NHANES_FPED() (Supplementary Material 4). To make the DASHI and MEDI comparable in different settings, we derived the mean dietary index percentile by dividing all DASHI by 50, the total possible score of DASHI, and all MEDI by 11, the total possible score of MEDI, and then both multiplied by 100.

As shown in Figure 5, the DASHI percentile was low in NHANES (37.42) and in the DASH trial control group that consumed a typical American diet (9.25), but it was very high in the DASH trial treatment groups (97.64 in the low sodium group and 97.37 in the medium sodium group). Similarly, the MEDI percentile was low in NHANES (32.12) and the MEDI low-fat control diet (45.45), while substantially higher in the Mediterranean diet with nuts (63.64) and in the Mediterranean diet with olive oil (72.73).

**Figure 5.**
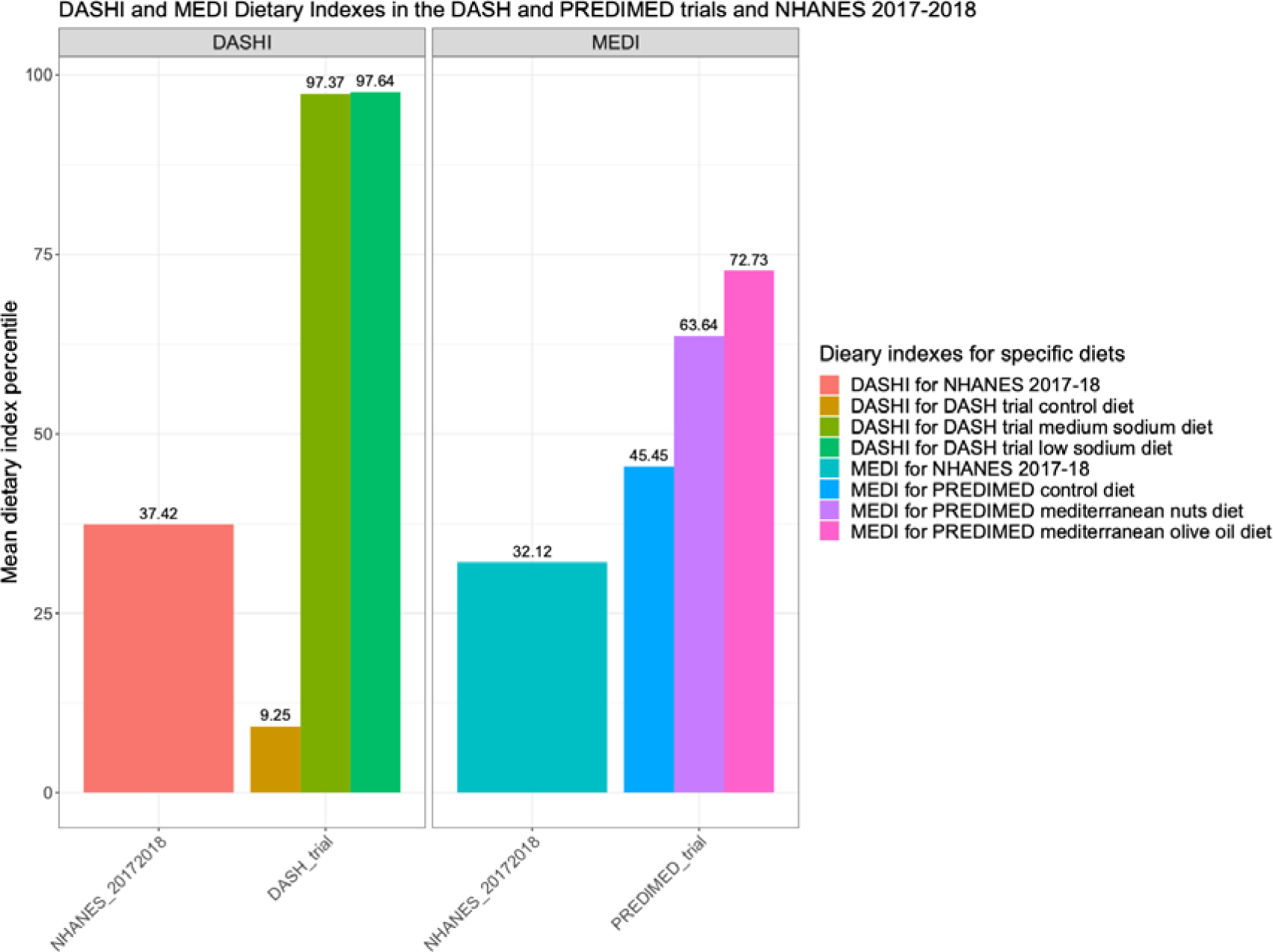
DASHI and MEDI dietary indexes in DASH and PREDIMED trials and NHANES in 2017-18

From the comparisons, there are several notable observations. First, dietary intake according to DASHI and MEDI varied considerably within the same NHANES data set. Specifically, for 2017-2018 NHANES data, DASHI scores were substantially higher than MEDI scores, indicating that the United States (US) population had higher adherence to the DASH compared to the Mediterranean diet in 2017-2018; this may reflect cultural-specific characteristics of the Mediterranean diet. Second, the DASHI and MEDI scores in NHANES were substantially lower than for those in treatment groups of the DASH and PREDIMED trials, indicating that the dietary intake in the US population has substantial room for improvement. Third, the difference in MEDI score between the Mediterranean diet with nuts treatment group and the low-fat control diet group was not as large as the difference in DASHI score between the low sodium DASH diet treatment group and the typical American diet control group; given that both the PREDIMED and DASH trial demonstrate significant cardiometabolic benefits of the Mediterranean and DASH diets, respectively,^4,18,22–24^ this may demonstrate that the free-living MED diet or DASH diet between 60 and 70 percentile MEDI or DASHI score may be a good alternative to the gold standard controlled feeding DASH diet in the DASH trial since the free-living dietary intervention is more sustainable and may still be able to generate long-term beneficial health effects as shown in the PREDIMED trial.

By using the standard frameworks proposed by dietaryindex, results from clinical and epidemiological studies can be compared to generate innovative insights. Additionally, forest plots for dietary indexes in the meta-analysis for clinical and epidemiological studies become possible when using dietaryindex. Thus, empowering the comparison between clinical and epidemiological studies with dietary intake measurements is one of the greatest strengths of dietaryindex.

#### Case study 2 A time series of cross-sectional computation of the HEI2020 from 2005 to 2018

We used dietaryindex to conduct a time series of cross-sectional computation of HEI2020 from 2005 to 2018 using NHANES 1-day dietary recall data for toddlers between 12 and 24 months of age and non-toddlers (children and adults) older than 24 months of age (Supplementary Material 4). As shown in Figure 6, across all years, toddlers and non-toddlers had suboptimal dietary intake as indicated by HEI2020 scores ranging from a low of 48.16 to a high of 56.64 out of 100, with toddlers having somewhat higher HEI2020 scores than non-toddlers in each year. In addition, by using the same standards in dietaryindex, we can visualize the trend of HEI2020 from 2005 to 2018 in toddlers and non-toddlers separately. Figure 6 shows that toddlers experienced twice the improvement of HEI2020 from 2005 to 2018 as did the child and adult sample (+2.87 vs. +0.86). Using dietaryindex, time series of cross-sectional calculations of dietary indexes for multiple years can be accomplished in a streamlined manner that generates high-quality, comparable results.

**Figure 6.**
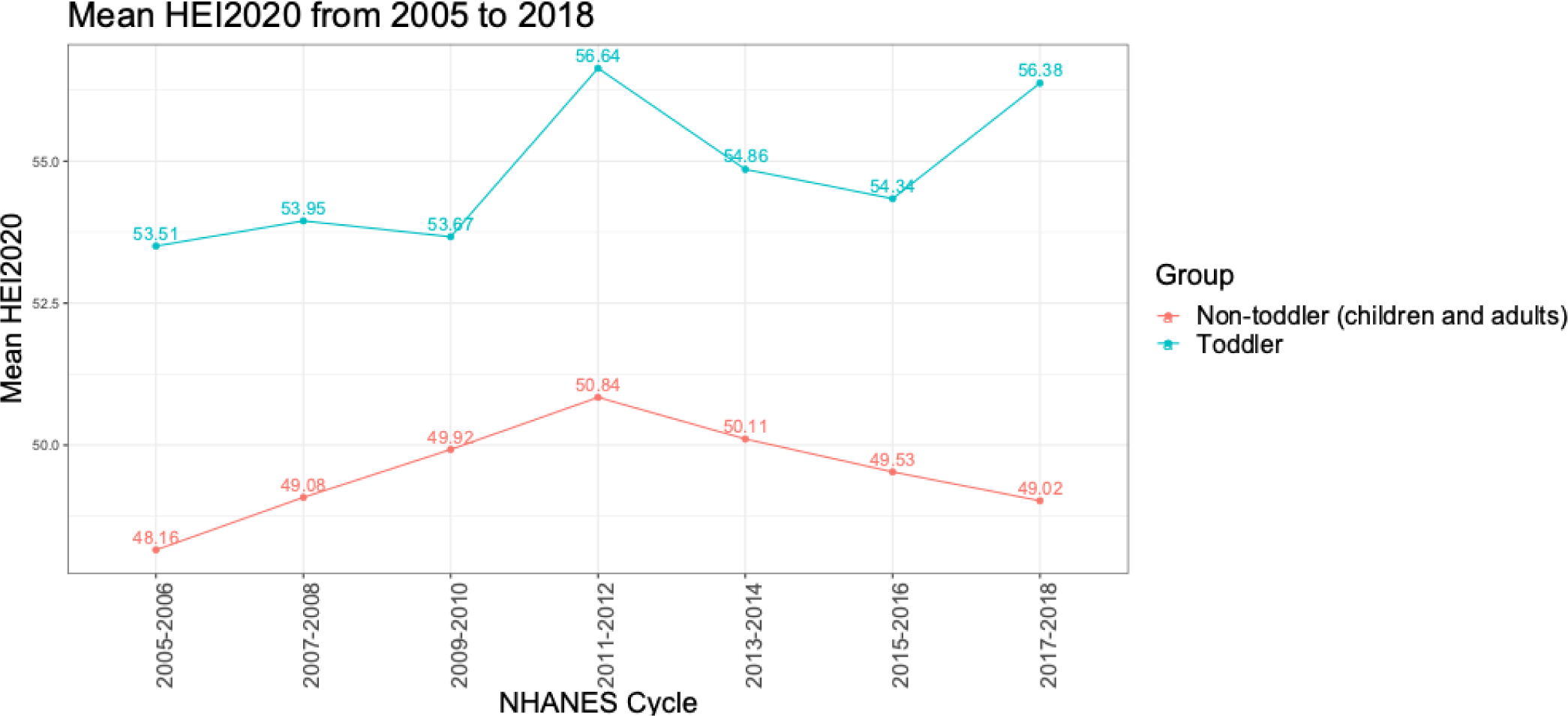
HEI2020 from 2005 to 2018 in toddlers and non-toddlers (children and adults) using NHANES data

#### Case study 3 Comprehensive Calculation of Multiple Dietary indexes, using NHANES data in 2017-2018

We conducted a comprehensive calculation of multiple dietary indexes, including HEI2020, AHEI, DASH, DASHI, MED, MEDI, and DII, with the NHANES data from 2017-2018 using dietaryindex. This method allowed us to visualize and compare intakes across different dietary indexes for any dietary assessment tool using a comparable approach.

Similar to case study 1, we first extracted the NHANES raw datasets (NHANES_20052006, NHANES_20072008, NHANES_20092010, NHANES_20112012, NHANES_20132014, NHANES_20152016, and NHANES_20172018) and stored the data in the dietaryindex package. Then, we used these data as inputs to calculate HEI2020, AHEI, DASH, DASHI, MED, MEDI, and DII using the relevant dietaryindex’s functions, including HEI2020_NHANES_FPED(), AHEI_NHANES_FPED(), DASH_NHANES_FPED(), DASHI_NHANES_FPED(), MED_NHANES_FPED(), MEDI_NHANES_FPED(), and DII_NHANES_FPED()). Two-day dietary recall data were used in the NHANES. Given that the dietary indexes have different total possible scores, to make them comparable, we derived the mean dietary index percentile for each dietary index by dividing their mean total scores by their total possible scores, then multiplying by 100.

As shown in Figure 7, based on NHANES 2017-2018, dietary intake per the dietary indexes showed substantial variation: AHEI – 33.63 percentile, DASH – 66.59 percentile, DASHI – 37.42, percentile, DII – 75 percentile, HEI2020 – 50.09 percentile, MED – 64.41 percentile, and MEDI – 32.12 percentile. Notably, scores based on the DASH, a quintile-based dietary index, and MED, a median-based dietary index, were consistently higher compared to scores based on serving size-based dietary indexes, which may suggest that DASH and MED overestimate the average dietary intake status compared to serving size-based dietary index methods. Unlike other dietary indexes, DII aims to measure dietary inflammatory potential where a higher score indicates a more pro-inflammatory diet, and a lower score indicates a more anti-inflammatory diet. Thus, the 75th percentile in DII demonstrates that in 2017-2018 the US population had a strongly pro-inflammatory diet, which is consistent with the results of other serving size-based dietary indexes.

**Figure 7.**
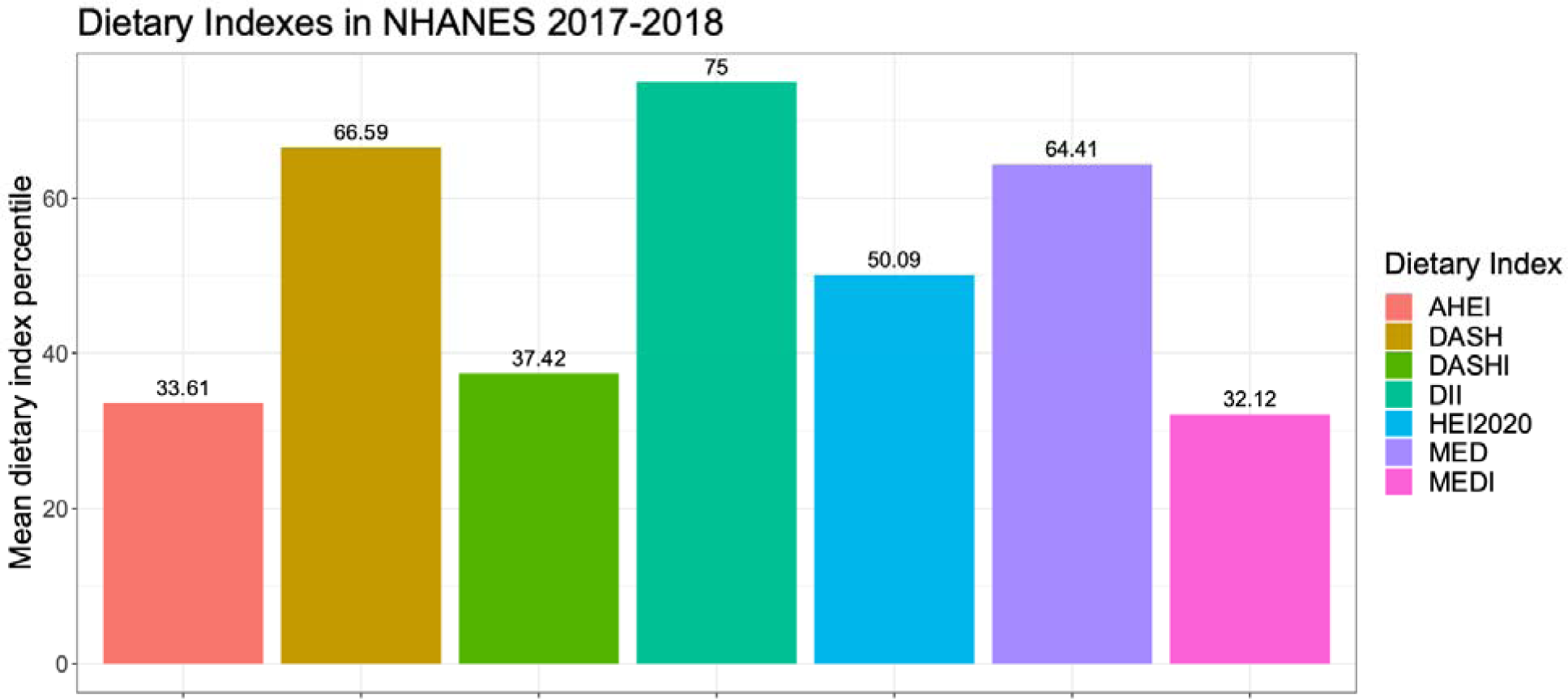
Multiple Dietary indexes, using the NHANES data in 2017-2018

By using dietaryindex, users can easily generate multiple dietary indexes and compare them in common nutritional datasets and tools, which is valuable during the exploratory stage of data analysis. Additionally, it enables researchers to have the ability to consider the selection of dietary indexes for a specific disease if the role of a diet on the disease is not well-studied.

## Discussion

We have developed dietaryindex, a proficient R package that offers versatile and user-friendly tools to standardize the consolidation of dietary intake data into several index-based dietary patterns. This tool was designed to cater to clinical and epidemiological research studies, with its most prominent strength lying in its inherent flexibility and convenience. The package incorporates a broad range of popular dietary index algorithms, including ACS 2020, AHEI, AHEIP, DASH, DASHI, DII, HEI2015, HEI2020, aMED, MEDI, and PHDI, among others. Additionally, dietaryindex has passed the internal and external validation process with remarkable accuracy.

This package also enables direct reference and incorporation of the output datasets from a variety of common dietary assessment tools, including but not limited to NHANES, ASA24, DHQ3, and Block FFQ, thereby catering to diverse research requirements and facilitating a more multifaceted understanding of dietary patterns. Moreover, the dietaryindex package goes beyond being just a computation tool -it provides an inclusive and user-friendly environment for researchers by providing internal access to multiple datasets, including NHANES data (2005-2018), DASH trial data, and PREDIMED trial data. Dietaryindex also supports multiple data entry options in NHANES and ASA24 by allowing users to enter either first-day data, second-day data, or combine both, and return results accordingly for NHANES functions and to enter multiple or single ASA24 recalls or records collected and always return the average dietary indexes for each individual per day. These convenient features can save significant time and resources for users by bypassing the complex process of dietary index calculations and focusing more on analyses and manuscript preparations.

We demonstrated three unique applications of the dietaryindex package in the case studies to shed light on dietaryindex’s capacity to inform various types of analysis, namely comparative studies between clinical and epidemiological research, time-series analyses of dietary indexes, and calculations of multiple dietary indexes within a single year. Specifically, we showed the ability of the dietaryindex package to compare data from clinical and epidemiological research studies using serving size-based dietary indexes, including DASHI and MEDI, thereby equipping researchers with an efficient tool for comparing outcomes from diverse studies; this will be pivotal to enabling a richer and more nuanced understanding of dietary patterns and their impact on health moving forward. Additionally, we demonstrated that the temporal analysis facilitated by the dietaryindex package is capable of delivering critical insights into changing dietary patterns over time, which could help enable the development of targeted dietary interventions to improve public health. Lastly, we illustrated that simultaneous computation of multiple dietary indexes within a single year allows for a thorough comparative analysis of different dietary adherence, thereby facilitating nuanced evaluations of dietary intakes in different settings and guiding personalized dietary recommendations. These varied applications underscore the package’s adaptability, versatility, and application to a variety of research scenarios, thereby enhancing its utility in the realm of nutritional epidemiology and biochemistry, the easy translation of clinical trial outcomes into population health interventions and surveillance – all using robust, replicable, and transparent methods that can greatly enhance the quality and comparability of dietary pattern research while minimizing the potential for any errors and miscalculations.

Although there are few tools for calculating Healthy Eating Index-2015,^8,9^ no standardized and open-source tools exist for calculating different dietary indexes from dietary data for both epidemiological and clinical studies using various nutritional assessment tools. To our knowledge, dietaryindex is the first bioinformatic tool to calculate multiple dietary indexes using different nutrition assessment tools. Therefore, dietaryindex provides a comprehensive framework for applying dietary indexes in different settings with a user-friendly design that enables most researchers to conduct analyses related to dietary indexes efficiently, bypassing most tedious works related to the literature research about dietary indexes and the serving size alignment of foods and nutrients for dietary indexes. This is a unique advantage of dietaryindex compared to other SAS and R packages for dietary index calculation.

There are some limitations of dietaryindex related to the measurement errors of nutritional assessments, arbitrary serving size definitions, and entry barriers to using R programming. The first limitation is that we did not directly validate other dietary index functions for NHANES, ASA24, and DHQ3 except HEI2015 since no standardized and open-source tools exist for calculating other dietary indexes. Nevertheless, dietaryindex follows a structured two-step computation process with an initial calculation of serving sizes followed by the calculation of dietary indexes. Given that all dietary indexes’ calculations were validated using simulated datasets, and all the serving size definitions proposed by dietaryindex were listed in Supplementary Material 1, we have indirectly confirmed the accuracy of other dietary index functions. The second limitation relates to the measurement errors in the nutritional assessment tools used. Nevertheless, the nutritional assessment tools supported by dietaryindex, including ASA24, DHQ3, Block FFQ, and the 24-hour dietary recall used in NHANES, have been extensively validated and commonly used in epidemiological and clinical studies.^25–29^ The third limitation is the arbitrary serving size definition for each dietary index in the dietaryindex package. We derived most serving size definitions in the dietaryindex about food groups, including vegetables, fruits, whole grains, sugar-sweetened beverages, nuts and legumes, and red/processed meat, from Chiuve et al. in order to standardize the serving size definitions in most dietary indexes;^10^ although arbitrary, using consistent serving size definitions across different dietary index enhances the precision and comparability of results. The fourth limitation is related to the food codes used in some dietary index functions to recognize specific foods, including sugar-sweetened beverages and low-fat dairy, in NHANES, ASA24, DHQ3, and Block FFQ. The default food codes were derived from the Food and Nutrient Database for Dietary Studies (FNDDS) in 2017-2018, which may not include all food codes from other years. However, dietaryindex allows users to input their own food codes for those specific foods that need to be manually indicated. The last limitation is the necessity of using R for the dietaryindex package, which may pose a challenge to some introductory R users. However, the user-friendliness of dietaryindex significantly mitigates this barrier. The package provides a range of single-line functions tailored to various nutritional assessment tools, which enable users to compute dietary indexes with minimal coding effort. The package also comes with explicit documentation and hands-on tutorials, supporting users through every step of the process, making it understandable and reproducible. The simplified functionality and robust support offered by the package make it an accessible and powerful tool for dietary research.

As an open-source project, dietaryindex is freely available on GitHub (https://github.com/jamesjiadazhan/dietaryindex), along with all the R codes used in the validation process, validation figure generation, and case studies (Supplementary Material 2, 3, 4).

## Conclusions

The dietaryindex R package is a user-friendly, versatile, and validated informatics tool that allows for the standard compilation and analysis of dietary pattern indexes for researchers in the fields of nutrition, epidemiology, and medicine. Dietaryindex is freely available on GitHub (https://github.com/jamesjiadazhan/dietaryindex), accompanied by detailed tutorials to assist all users in leveraging its capabilities.

## Supporting information

Supplement 1 dietaryindex serving size definitions and scoring algorithms

Supplement 2 Validation file for publication

Supplement 3 Validation Figures

Supplement 4 R scripts for case studies

