## Supplement 2 Validation file for publication for "Dietaryindex: A User-Friendly and Versatile R Package for Standardizing Dietary Pattern Analysis in Epidemiological and Clinical Studies": HEI2015_ASA24_dietaryindex.r

```
library(dplyr)
library(haven)
library(readr)
# Loading dependency packages first can help avoid the namespace conflict if you want to use dplyr by yourself later in addition to the internal use of dplyr within the diataryindex package
# Load the dietaryindex package
library(dietaryindex)

# set working directory
setwd("/Users/james/Desktop/Emory University - Ph.D./dietaryindex_package/dietaryindex/Validation file for publication/HEI2015_ASA24_example_data")

# Read in the ASA24 data
ASA24_exp = read_csv("THR_2022-09-13_86071_Totals.csv")

# Calculate HEI-2015 score using the dietaryindex package
HEI2015_ASA24_exp = HEI2015_ASA24(ASA24_exp)

# Write the HEI-2015 score calculated by dietaryindex to a csv file
write_csv(HEI2015_ASA24_exp, "HEI2015_ASA24_dietaryindex.csv")
```
