## Supplement 2 Validation file for publication for "Dietaryindex: A User-Friendly and Versatile R Package for Standardizing Dietary Pattern Analysis in Epidemiological and Clinical Studies": SAS code ASA24 validation.sas

```
*ASA24 SAS code;
/*1. Create a folder on your computer “home folder”, and save the ASA24 data, and the required HEI-2015 macro in it. Specify the path to the folder. */
/*2. Filename here specifies the input file. */

filename Totals '/home/u59265813/sasuser.v94/AA cohort/ASA24/dietaryindex example data/THR_2022-09-13_86071_Totals.csv'; /*In this example, the ASA24-2016 or ASA24-2018 Daily Total Nutrient and Pyramid Equivalents data “Totals”, are saved in a folder called “Totals”, within the “home” folder. The data are in csv format. */

/*3. Create a folder in the "home" folder, where the output file, containing HEI-2015 component and total scores for each respondent, for the intake day, are to be exported. Specify the name of the folder. */

filename res '/home/u59265813/sasuser.v94/AA cohort/ASA24/dietaryindex example data/HEI2015_ASA24_NCI_SAS.csv'; /*In this Example, the folder is called “RES”, within the “home” folder, and the exported results will be a csv file called “hei2015r”. */

/*4. Read in required HEI-2015 scoring macro. This macro must be saved within the home folder. */

%include '/home/u59265813/sasuser.v94/AA cohort/ASA24/dietaryindex example data/hei2015.score.macro.sas';

/*NOTE: Once you have completed all the steps above, all you need to do is run the SAS program below. Unless you used different names for your dataset and folders, no other action is required from you. */

/*Step 1.
Input daily total data and create five additional required variables.  These variables are:  
FWHOLEFRT, MONOPOLY, VTOTALLEG, VDRKGRLEG, PFALLPROTLEG, and PFSEAPLANTLEG
*/

Proc import datafile=Totals
  Out=Totals
  Dbms=csv
  Replace;
  Getnames=yes;
Run;


DATA Totals;
  SET Totals;

  FWHOLEFRT=F_CITMLB+F_OTHER;

  MONOPOLY=MFAT+PFAT;

  VTOTALLEG=V_TOTAL+V_LEGUMES;
  VDRKGRLEG=V_DRKGR+V_LEGUMES;

  PFALLPROTLEG=PF_MPS_TOTAL+PF_EGGS+PF_NUTSDS+PF_SOY+PF_LEGUMES; 
  PFSEAPLANTLEG=PF_SEAFD_HI+PF_SEAFD_LOW+PF_NUTSDS+PF_SOY+PF_LEGUMES;
  
run; 


/*Step 2. 
 Runs the HEI2015 scoring macro which calculates intake density amounts and HEI scores.
*/

%HEI2015 (indat=Totals,
          kcal= KCAL,
	  vtotalleg= VTOTALLEG,
	  vdrkgrleg= VDRKGRLEG,
	  f_total= F_TOTAL,
	  fwholefrt=FWHOLEFRT,
	  g_whole= G_WHOLE,
	  d_total= D_TOTAL,
          pfallprotleg= PFALLPROTLEG,
	  pfseaplantleg= PFSEAPLANTLEG,
	  monopoly=MONOPOLY,
	  satfat=SFAT,
	  sodium=SODI,
	  g_refined=G_REFINED,
	  add_sugars=ADD_SUGARS,
	  outdat=hei2015);
 
run;

/*Step 3.
 Displays and saves the results.
*/ 

Data hei2015r (keep=UserName UserID RecallNo kcal HEI2015C1_TOTALVEG HEI2015C2_GREEN_AND_BEAN HEI2015C3_TOTALFRUIT
      HEI2015C4_WHOLEFRUIT HEI2015C5_WHOLEGRAIN HEI2015C6_TOTALDAIRY HEI2015C7_TOTPROT HEI2015C8_SEAPLANT_PROT 
      HEI2015C9_FATTYACID HEI2015C10_SODIUM HEI2015C11_REFINEDGRAIN HEI2015C12_SFAT HEI2015C13_ADDSUG HEI2015_TOTAL_SCORE);
  Set hei2015;
  Run;

proc means n nmiss min max mean data=hei2015r;
run;

proc export data= hei2015r
  file=RES
  dbms=csv
  replace;
run;
```
