## Supplement 2 Validation file for publication for "Dietaryindex: A User-Friendly and Versatile R Package for Standardizing Dietary Pattern Analysis in Epidemiological and Clinical Studies": hei2015.score.macro.sas

```
/*************************************************************************/
/*************************************************************************/
/*                                                                       */
/*                 THE HEI-2015 SCORING MACRO                            */                                               
/*                  (hei2015.score.macro.sas)                            */
/*************************************************************************/
/*                     VERSION 1.0        06/25/2017                     */   
/*                                                                       */
/*                                                                       */
/* This HEI-2015 macro is to be used to calculate densities and          */
/* and HEI-2015 component and total scores.                              */
/*								         */	
/* The macro requires an input dataset with variables for each of        */
/* the HEI-2015 components, noted below.                                 */
/*								         */	
/* The resulting dataset, which is named by the user, contains the       */
/* same variables as the supplied dataset, and creates 27 new            */
/* variables. These include:					         */
/*                                                                       */				    
/*   The densities (per 1000 kcal) or percent (of total calories)        */
/*    for each of the 13 HEI-2015 components.			         */
/*                                                                       */
/*   Scores for the 13 components of the HEI-2015.                       */
/*                                                                       */
/*   The total HEI-2015 score, which is the sum of the                   */
/*    scores for the 13 components.                                      */
/*                                                                       */                                                                  
/* The syntax for calling the macro is:                                  */
/*                                                                       */
/* %HEI 2015   							         */
/* (indat=,kcal=,vtotalleg=,vdrkgrleg=,f_total=,fwholefrt=,g_whole=      */
/*    d_total=,pfallprotleg=,pfseaplantleg=,monopoly=,satfat=,sodium=,   */
/*    g_refined=,add_sugars=,outdat=)                                    */
/*                                                                       */
/*  where                                                                */
/*                                                                       */
/*   "indat"        * Specifies the dataset to be used.                  */
/*                                                                       */
/*   "kcal"         * Specifies calorie amount.                          */
/*                                                                       */
/*   "vtotalleg"    * Specifies the intake of total veg plus             */
/*                      legumes in cup eq.                               */
/*                                                                       */
/*   "vdrkgrleg"    * Specifies the intake of dark green veg             */
/*                      plus legumes in cup eq.                          */
/*                                                                       */
/*   "f_total"      * Specifies the intake of total fruit in cup eq      */
/*                                                                       */
/*   "fwholefrt"    * Specifies the intake of whole fruit in cup eq.     */
/*                                                                       */
/*   "g_whole"      * Specifies the intake of whole grain in oz. eq.     */
/*                                                                       */
/*   "d_total"      * Specifies the intake of total dairy in cup eq.     */
/*                                                                       */
/*   "pfallprotleg" * Specifies the intake of total protein              */
/*                      (includes legumes) in oz. eq.                    */
/*                                                                       */
/*   "pfseaplantleg"  * Specifies the intake of seafood, fish and plant  */
/*                      protein (includes legumes) in oz. eq.            */
/*                                                                       */
/*   "monopoly"       * Specifies the grams of mono fat plus poly fat.   */
/*                                                                       */
/*   "satfat"         * Specifies the grams of saturated fat.            */
/*                                                                       */
/*   "sodium"         * Specifies the mg of sodium.                      */
/*                                                                       */                                                                 
/*   "g_refined"      * Specifies the intake of refined                  */
/*                       grain in oz. eq.                                */
/*                                                                       */
/*   "add_sugars"     * Specifies the intake of added sugars in tsp. eq. */
/*                                                                       */
/*   "outdat"         * Specifies the name of the resulting dataset.     */
/*								         */
/*                                                                       */                                                                 
/* Caution:  variable names "FARMIN", "FARMAX", "SODMIN",                */
/*   "SODMAX", "RGMIN", "RGMAX", "SFATMIN", "SFATMAX", "ADDSUGMIN",      */
/*   "ADDSUGMAX" are reserved for this macro.                            */
/*                                                                       */
/*                                                                       */
/*************************************************************************/
;

                              


%macro HEI2015 (indat=,kcal=,vtotalleg=,vdrkgrleg=,f_total=,fwholefrt=,g_whole=,d_total=,
  pfallprotleg=,pfseaplantleg=,monopoly=,satfat=,sodium=,g_refined=,add_sugars=,outdat=);

data &outdat (drop=FARMIN FARMAX SODMAX SODMIN RGMIN RGMAX SFATMIN SFATMAX ADDSUGMIN ADDSUGMAX);
  set &indat;

  IF &kcal > 0 then VEGDEN=&vtotalleg/(&kcal/1000);
  HEI2015C1_TOTALVEG=5*(VEGDEN/1.1);
  IF HEI2015C1_TOTALVEG > 5 THEN HEI2015C1_TOTALVEG=5;
  IF VEGDEN=0 THEN HEI2015C1_TOTALVEG=0;

  IF &kcal > 0 then GRBNDEN=&vdrkgrleg/(&kcal/1000);
  HEI2015C2_GREEN_AND_BEAN=5*(GRBNDEN/0.2);
  IF HEI2015C2_GREEN_AND_BEAN > 5 THEN HEI2015C2_GREEN_AND_BEAN=5;
  IF GRBNDEN=0 THEN HEI2015C2_GREEN_AND_BEAN=0;

  IF &kcal > 0 then FRTDEN=&f_total/(&kcal/1000);
  HEI2015C3_TOTALFRUIT=5*(FRTDEN/0.8);
  IF HEI2015C3_TOTALFRUIT > 5 THEN HEI2015C3_TOTALFRUIT=5;
  IF FRTDEN=0 THEN HEI2015C3_TOTALFRUIT=0;	

  IF &kcal > 0 then WHFRDEN=&fwholefrt/(&kcal/1000);
  HEI2015C4_WHOLEFRUIT=5*(WHFRDEN/0.4); 
  IF HEI2015C4_WHOLEFRUIT > 5 THEN HEI2015C4_WHOLEFRUIT=5;
  IF WHFRDEN=0 THEN HEI2015C4_WHOLEFRUIT=0;	

  IF &kcal > 0 then WGRNDEN=&g_whole/(&kcal/1000);
  HEI2015C5_WHOLEGRAIN=10*(WGRNDEN/1.5);
  IF HEI2015C5_WHOLEGRAIN > 10 THEN HEI2015C5_WHOLEGRAIN=10;
  IF WGRNDEN=0 THEN HEI2015C5_WHOLEGRAIN=0;

  IF &kcal > 0 then DAIRYDEN=&d_total/(&kcal/1000);
  HEI2015C6_TOTALDAIRY=10*(DAIRYDEN/1.3);
  IF HEI2015C6_TOTALDAIRY > 10 THEN HEI2015C6_TOTALDAIRY=10;
  IF DAIRYDEN=0 THEN HEI2015C6_TOTALDAIRY=0;

  IF &kcal > 0 then PROTDEN=&pfallprotleg/(&kcal/1000);
  HEI2015C7_TOTPROT=5*(PROTDEN/2.5);
  IF HEI2015C7_TOTPROT > 5 THEN HEI2015C7_TOTPROT=5;
  IF PROTDEN=0 THEN HEI2015C7_TOTPROT=0;

  IF &kcal > 0 then SEAPLDEN=&pfseaplantleg/(&kcal/1000);
  HEI2015C8_SEAPLANT_PROT=5*(SEAPLDEN/0.8);
  IF HEI2015C8_SEAPLANT_PROT > 5 THEN HEI2015C8_SEAPLANT_PROT=5;
  IF SEAPLDEN=0 THEN HEI2015C8_SEAPLANT_PROT=0;

  IF &satfat > 0 THEN FARATIO=&monopoly/&satfat;
  FARMIN=1.2;
  FARMAX=2.5;
  if &satfat=0 and &monopoly=0 then HEI2015C9_FATTYACID=0;
    else if &satfat=0 and &monopoly > 0 then HEI2015C9_FATTYACID=10;
    else if FARATIO >= FARMAX THEN HEI2015C9_FATTYACID=10;
    else if FARATIO <= FARMIN THEN HEI2015C9_FATTYACID=0;
    else HEI2015C9_FATTYACID=10* ( (FARATIO-FARMIN) / (FARMAX-FARMIN) );

  IF &kcal > 0 then SODDEN=&sodium/&kcal;
  SODMIN=1.1;
  SODMAX=2.0;
  IF SODDEN <= SODMIN THEN HEI2015C10_SODIUM=10;
    ELSE IF SODDEN >= SODMAX THEN HEI2015C10_SODIUM=0;
    ELSE HEI2015C10_SODIUM=10 - (10 * (SODDEN-SODMIN) / (SODMAX-SODMIN) );

  IF &kcal > 0 then RGDEN=&g_refined/(&kcal/1000);
  RGMIN=1.8;
  RGMAX=4.3;
  IF RGDEN <= RGMIN THEN HEI2015C11_REFINEDGRAIN=10;
    ELSE IF RGDEN >= RGMAX THEN HEI2015C11_REFINEDGRAIN=0;
    ELSE HEI2015C11_REFINEDGRAIN=10 - ( 10* (RGDEN-RGMIN) / (RGMAX-RGMIN) ); 
 
  IF &kcal > 0 then SFAT_PERC=100*(&satfat*9/&kcal); 
  SFATMIN=8;
  SFATMAX=16;
  IF SFAT_PERC >= SFATMAX THEN HEI2015C12_SFAT=0;
    ELSE IF SFAT_PERC <= SFATMIN THEN HEI2015C12_SFAT=10;
    ELSE HEI2015C12_SFAT= 10 - ( 10* (SFAT_PERC-SFATMIN) / (SFATMAX-SFATMIN) );

  IF &kcal > 0 then ADDSUG_PERC=100*(&add_sugars*16/&kcal); 
  ADDSUGMIN=6.5;
  ADDSUGMAX=26;
  IF ADDSUG_PERC >= ADDSUGMAX THEN HEI2015C13_ADDSUG=0;
    ELSE IF ADDSUG_PERC <= ADDSUGMIN THEN HEI2015C13_ADDSUG=10;
    ELSE HEI2015C13_ADDSUG= 10 - ( 10* (ADDSUG_PERC-ADDSUGMIN) / (ADDSUGMAX-ADDSUGMIN) );


IF &kcal=0 THEN DO;
  HEI2015C1_TOTALVEG=0; HEI2015C2_GREEN_AND_BEAN=0; HEI2015C3_TOTALFRUIT=0; HEI2015C4_WHOLEFRUIT=0; HEI2015C5_WHOLEGRAIN=0; HEI2015C6_TOTALDAIRY=0;
  HEI2015C7_TOTPROT=0;  HEI2015C8_SEAPLANT_PROT=0; HEI2015C9_FATTYACID=0; HEI2015C10_SODIUM=0; HEI2015C11_REFINEDGRAIN=0; HEI2015C12_SFAT=0; HEI2015C13_ADDSUG=0;
  END;

/**Calculate HEI-2015 total score**/
/*total HEI-2015 score is the sum of 13 HEI component scores*/

HEI2015_TOTAL_SCORE = HEI2015C1_TOTALVEG + HEI2015C2_GREEN_AND_BEAN + HEI2015C3_TOTALFRUIT + HEI2015C4_WHOLEFRUIT + HEI2015C5_WHOLEGRAIN + HEI2015C6_TOTALDAIRY +
  HEI2015C7_TOTPROT + HEI2015C8_SEAPLANT_PROT + HEI2015C9_FATTYACID + HEI2015C10_SODIUM + HEI2015C11_REFINEDGRAIN + HEI2015C12_SFAT + HEI2015C13_ADDSUG;


LABEL HEI2015_TOTAL_SCORE='TOTAL HEI-2015 SCORE'
      HEI2015C1_TOTALVEG='HEI-2015 COMPONENT 1 TOTAL VEGETABLES'
      HEI2015C2_GREEN_AND_BEAN='HEI-2015 COMPONENT 2 GREENS AND BEANS'
      HEI2015C3_TOTALFRUIT='HEI-2015 COMPONENT 3 TOTAL FRUIT'
      HEI2015C4_WHOLEFRUIT='HEI-2015 COMPONENT 4 WHOLE FRUIT'
      HEI2015C5_WHOLEGRAIN='HEI-2015 COMPONENT 5 WHOLE GRAINS'
      HEI2015C6_TOTALDAIRY='HEI-2015 COMPONENT 6 DAIRY'
      HEI2015C7_TOTPROT='HEI-2015 COMPONENT 7 TOTAL PROTEIN FOODS'

      HEI2015C8_SEAPLANT_PROT='HEI-2015 COMPONENT 8 SEAFOOD AND PLANT PROTEIN'
      HEI2015C9_FATTYACID='HEI-2015 COMPONENT 9 FATTY ACID RATIO'
      HEI2015C10_SODIUM='HEI-2015 COMPONENT 10 SODIUM'
      HEI2015C11_REFINEDGRAIN='HEI-2015 COMPONENT 11 REFINED GRAINS'
      HEI2015C12_SFAT='HEI-2015 COMPONENT 12 SAT FAT'
      HEI2015C13_ADDSUG='HEI-2015 COMPONENT 13 ADDED SUGAR'
      VEGDEN='DENSITY OF TOTAL VEGETABLES PER 1000 KCAL'
      GRBNDEN='DENSITY OF DARK GREEN VEG AND BEANS PER 1000 KCAL'
      FRTDEN='DENSITY OF TOTAL FRUIT PER 1000 KCAL'
      WHFRDEN='DENSITY OF WHOLE FRUIT PER 1000 KCAL'
      WGRNDEN='DENSITY OF WHOLE GRAIN PER 1000 KCAL'
      DAIRYDEN='DENSITY OF DAIRY PER 1000 KCAL'
      PROTDEN='DENSITY OF TOTAL PROTEIN PER 1000 KCAL'
      SEAPLDEN='DENSITY OF SEAFOOD AND PLANT PROTEIN PER 1000 KCAL'
      FARATIO='FATTY ACID RATIO'
      SODDEN='DENSITY OF SODIUM PER 1000 KCAL'
      RGDEN='DENSITY OF REFINED GRAINS PER 1000 KCAL'
      SFAT_PERC='PERCENT OF CALORIES FROM SAT FAT'
      ADDSUG_PERC='PERCENT OF CALORIES FROM ADDED SUGAR'
      ;

run;


%mend HEI2015;


/*  END OF THE HEI2015 MACRO                                       */
/*******************************************************************/
```
