## Supplement 2 Validation file for publication for "Dietaryindex: A User-Friendly and Versatile R Package for Standardizing Dietary Pattern Analysis in Epidemiological and Clinical Studies": NHANES SAS for HEI2015.sas

```
libname NH '/home/u59265813/sasuser.v94/AA cohort/NHANES/NH';  
libname FPED '/home/u59265813/sasuser.v94/AA cohort/NHANES/FPED'; /*In this Example, the FPED data are in a folder called “FPED”, and the NHANES and Demographic data are in a folder called “NH”, all saved within the “home” folder. These are SAS datasets. */

filename RES '/home/u59265813/sasuser.v94/AA cohort/NHANES/RES'; /*In this Example, the folder is called “RES”, within the “home” folder, and the exported results will be a csv file called “hei2015r”. */


%include '/home/u59265813/sasuser.v94/AA cohort/NHANES/hei2015.score.macro.sas';


title 'HEI-2015 scores for NHANES 2011-2012 day 1, AGE >= 2, RELIABLE DIETS, Include Pregnant and Lactating Women';

/*Step 1: locate the required datasets and variables */

*part a: get FPED data per day;
data FPED;
 set FPED.fped_dr1tot_1718;
run;

*part b: get individual total nutrient intake if reliable recall status;
data NUTRIENT (keep=SEQN WTDRD1 DR1TKCAL DR1TSFAT DR1TALCO DR1TSODI DR1DRSTZ DR1TMFAT DR1TPFAT);
  set NH.DR1TOT_J;
  if DR1DRSTZ=1; /*reliable dietary recall status*/
run;

*part c: get demographic data for persons aged two and older;
data DEMO (keep=SEQN RIDAGEYR RIAGENDR SDDSRVYR SDMVPSU SDMVSTRA);
  set NH.DEMO_J;
  if RIDAGEYR >= 2;
run;

/*Step 2: Combine the required datasets*/

proc sort data=FPED;
  by SEQN;
run;

proc sort data=NUTRIENT;
  by SEQN;
run;

proc sort data=DEMO;
  by SEQN;
run;

data COHORT;
  merge NUTRIENT (in=N) DEMO (in=D) FPED;
  by SEQN;
  if N and D;
run;

/*Step 3: Creates additional required variables: FWHOLEFRT, MONOPOLY, VTOTALLEG, VDRKGRLEG, PFALLPROTLEG and PFSEAPLANTLEG */


data COHORT;
  set COHORT;
  by SEQN;

  FWHOLEFRT=DR1T_F_CITMLB+DR1T_F_OTHER;

  MONOPOLY=DR1TMFAT+DR1TPFAT;

  VTOTALLEG=DR1T_V_TOTAL+DR1T_V_LEGUMES;
  VDRKGRLEG=DR1T_V_DRKGR+DR1T_V_LEGUMES;

  PFALLPROTLEG=DR1T_PF_MPS_TOTAL+DR1T_PF_EGGS+DR1T_PF_NUTSDS+DR1T_PF_SOY+DR1T_PF_LEGUMES; 
  PFSEAPLANTLEG=DR1T_PF_SEAFD_HI+DR1T_PF_SEAFD_LOW+DR1T_PF_NUTSDS+DR1T_PF_SOY+DR1T_PF_LEGUMES;
run;

/*Step 4: Apply the HEI-2015 scoring macro. */

%HEI2015 (indat= COHORT, 
          kcal= DR1TKCAL, 
	  vtotalleg= VTOTALLEG, 
	  vdrkgrleg= VDRKGRLEG, 
	  f_total= DR1T_F_TOTAL, 
	  fwholefrt= FWHOLEFRT, 
	  g_whole= DR1T_G_WHOLE, 
	  d_total= DR1T_D_TOTAL, 
          pfallprotleg= PFALLPROTLEG, 
	  pfseaplantleg= PFSEAPLANTLEG, 
	  monopoly= MONOPOLY, 
	  satfat= DR1TSFAT, 
	  sodium= DR1TSODI, 
	  g_refined= DR1T_G_REFINED, 
	  add_sugars= DR1T_ADD_SUGARS, 
	  outdat= HEI2015); 

/*Step 5: Displays and saves the results. */ 

*part a: this program saves one HEI-2015 score for each individual, based on one 24HR;


data HEI2015R (keep=SEQN DR1TKCAL HEI2015C1_TOTALVEG HEI2015C2_GREEN_AND_BEAN HEI2015C3_TOTALFRUIT HEI2015C4_WHOLEFRUIT 
      HEI2015C5_WHOLEGRAIN HEI2015C6_TOTALDAIRY HEI2015C7_TOTPROT HEI2015C8_SEAPLANT_PROT HEI2015C9_FATTYACID HEI2015C10_SODIUM
      HEI2015C11_REFINEDGRAIN HEI2015C12_SFAT HEI2015C13_ADDSUG HEI2015_TOTAL_SCORE); 
  set HEI2015; 
  run; 
 
*part b: calculates an unweighted mean across all individuals in group; 
 
proc means n nmiss min max mean data=HEI2015R; 
run; 
 
 
*part c: saves results as CSV file one line per day; 
 
proc export data=HEI2015R 
  outfile='/home/u59265813/sasuser.v94/AA cohort/NHANES/RES/HEI2015R.csv'
  dbms=csv 
  replace; 
run; 

proc export data=NH.DR1TOT_J
  outfile='/home/u59265813/sasuser.v94/AA cohort/NHANES/RES/NUTRIENT.csv'
  dbms=csv 
  replace; 
run; 

proc export data=FPED.fped_dr1tot_1718
  outfile='/home/u59265813/sasuser.v94/AA cohort/NHANES/RES/FPED.csv'
  dbms=csv 
  replace; 
run; 

proc export data=NH.DEMO_J
  outfile='/home/u59265813/sasuser.v94/AA cohort/NHANES/RES/DEMO.csv'
  dbms=csv 
  replace; 
run; 
```
