## Supplement 2 Validation file for publication for "Dietaryindex: A User-Friendly and Versatile R Package for Standardizing Dietary Pattern Analysis in Epidemiological and Clinical Studies": Validation.r

```
install.packages("devtools") #If you don't have "devtools" installed already
devtools::install_github("jamesjiadazhan/dietaryindex") # Install the package from GitHub

library(dietaryindex) # Load the package
library(readr)
library(dplyr)

# Load the data
## set up the working directory
setwd("/Users/james/Desktop/Emory University - Ph.D./dietaryindex_package/dietaryindex/Validation file for publication/Raw validation files")
data("NHANES_20172018")
ACS2020_V1_VALIDATION = read_csv("ACS2020_V1_validation.csv")
ACS2020_V2_VALIDATION = read_csv("ACS2020_V2_validation.csv")
AHEI_VALIDATION = read_csv("AHEI_validation.csv")
AHEIP_VALIDATION = read_csv("AHEIP_validation.csv")
DASH_VALIDATION = read_csv("DASH_validation.csv")
DASHI_VALIDATION = read_csv("DASHI_validation.csv")
DII_VALIDATION = read_csv("DII_validation.csv")
DII_VALIDATION = read_csv("/Users/james/Desktop/Emory University - Ph.D./dietaryindex_package/dietaryindex/Validation file for publication/Raw validation files/DII_validation.csv")
HEI2015_VALIDATION = read_csv("HEI2015_validation.csv")
HEI2020_VALIDATION = read_csv("HEI2020_validation.csv")
MED_VALIDATION = read_csv("MED_validation.csv")
MEDI_V2_VALIDATION = read_csv("MEDI_V2_validation.csv")
MEDI_VALIDATION = read_csv("MEDI_validation.csv")
PHDI_VALIDATION = read_csv("PHDI_validation.csv")

# Generate the validation results and save them

setwd("/Users/james/Desktop/Emory University - Ph.D./dietaryindex_package/dietaryindex/Validation file for publication/HEI2015_NHANES_SAS_1718")
## HEI2015_NHANES_FPED validation using the SAS codes from National Cancer Institute (https://epi.grants.cancer.gov/hei/sas-code.html)
# day 1 only
HEI2015_NHANES_FPED_1718 = HEI2015_NHANES_FPED(
    FPED_PATH = NHANES_20172018$FPED,
    NUTRIENT_PATH = NHANES_20172018$NUTRIENT,
    DEMO_PATH = NHANES_20172018$DEMO
)

# save the result
write_csv(HEI2015_NHANES_FPED_1718, "dietaryindex_HEI2015_1718.csv")

# read in the SAS result using the NCI SAS codes
HEI2015_NHANES_SAS_1718 = read_csv("SAS_HEI2015_1718.csv")

# check if the HEI2015_TOTAL_SCORE in the SAS result is the same as the HEI2015_ALL in the R result in two decimals
HEI2015_NHANES_FPED_1718$HEI2015_ALL = round(HEI2015_NHANES_FPED_1718$HEI2015_ALL, 2)
HEI2015_NHANES_SAS_1718$HEI2015_TOTAL_SCORE = round(HEI2015_NHANES_SAS_1718$HEI2015_TOTAL_SCORE, 2)
table(HEI2015_NHANES_FPED_1718$HEI2015_ALL == HEI2015_NHANES_SAS_1718$HEI2015_TOTAL_SCORE)
## all results are TRUE, which means the R result is the same as the SAS result

setwd("/Users/james/Desktop/Emory University - Ph.D./dietaryindex_package/dietaryindex/Validation file for publication/Final validation files")

## ACS2020 validation (Version 1)
ACS2020_V1_VALIDATION_RESULT = ACS2020_V1(
    SERV_DATA = ACS2020_V1_VALIDATION, 
    RESPONDENTID = ACS2020_V1_VALIDATION$id, 
    GENDER = ACS2020_V1_VALIDATION$gender, 
    VEG_SERV_ACS2020 = ACS2020_V1_VALIDATION$vegetable, 
    VEG_ITEMS_SERV_ACS2020 = ACS2020_V1_VALIDATION$vegetable_unique, 
    FRT_SERV_ACS2020 = ACS2020_V1_VALIDATION$fruit, 
    FRT_ITEMS_SERV_ACS2020 = ACS2020_V1_VALIDATION$fruit_unique, 
    WGRAIN_SERV_ACS2020 = ACS2020_V1_VALIDATION$whole_grain, 
    REDPROC_MEAT_SERV_ACS2020 = ACS2020_V1_VALIDATION$red_meat, 
    HPFRG_RATIO_SERV_ACS2020 = ACS2020_V1_VALIDATION$process_food, 
    SSB_FRTJ_SERV_ACS2020 = ACS2020_V1_VALIDATION$ssb)

# Merge the validation data with the result
ACS2020_V1_VALIDATION_RESULT_FINAL = ACS2020_V1_VALIDATION %>%
    left_join(ACS2020_V1_VALIDATION_RESULT, by = c("id" = "RESPONDENTID"))

# Save the result
write_csv(ACS2020_V1_VALIDATION_RESULT_FINAL, "ACS2020_V1_validation_result.csv")

## ACS2020 validation (Version 2)
ACS2020_V2_VALIDATION_RESULT = ACS2020_V2(
    SERV_DATA = ACS2020_V2_VALIDATION, 
    RESPONDENTID = ACS2020_V2_VALIDATION$id, 
    GENDER = ACS2020_V2_VALIDATION$gender, 
    VEG_SERV_ACS2020 = ACS2020_V2_VALIDATION$vegetable, 
    VEG_ITEMS_SERV_ACS2020 = ACS2020_V2_VALIDATION$vegetable_unique, 
    FRT_SERV_ACS2020 = ACS2020_V2_VALIDATION$fruit, 
    FRT_ITEMS_SERV_ACS2020 = ACS2020_V2_VALIDATION$fruit_unique, 
    WGRAIN_SERV_ACS2020 = ACS2020_V2_VALIDATION$whole_grain, 
    REDPROC_MEAT_SERV_ACS2020 = ACS2020_V2_VALIDATION$red_meat, 
    HPFRG_SERV_ACS2020 = ACS2020_V2_VALIDATION$process_food, 
    SSB_FRTJ_SERV_ACS2020 = ACS2020_V2_VALIDATION$ssb,
    TOTALKCAL_ACS2020 = ACS2020_V2_VALIDATION$kcal)

# Merge the validation data with the result
ACS2020_V2_VALIDATION_RESULT_FINAL = ACS2020_V2_VALIDATION %>%
    left_join(ACS2020_V2_VALIDATION_RESULT, by = c("id" = "RESPONDENTID"))

# Save the result
write_csv(ACS2020_V2_VALIDATION_RESULT_FINAL, "ACS2020_V2_validation_result.csv")

## AHEI validation
AHEI_VALIDATION_RESULT = AHEI(
    SERV_DATA = AHEI_VALIDATION,
    RESPONDENTID = AHEI_VALIDATION$id,
    GENDER = AHEI_VALIDATION$gender,
    TOTALKCAL_AHEI = AHEI_VALIDATION$kcal,
    VEG_SERV_AHEI = AHEI_VALIDATION$vegetable,
    FRT_SERV_AHEI = AHEI_VALIDATION$fruit,
    WGRAIN_SERV_AHEI = AHEI_VALIDATION$whole_grain,
    NUTSLEG_SERV_AHEI = AHEI_VALIDATION$nut_legume,
    N3FAT_SERV_AHEI = AHEI_VALIDATION$n3_fat,
    PUFA_SERV_AHEI = AHEI_VALIDATION$pufa,
    SSB_FRTJ_SERV_AHEI = AHEI_VALIDATION$ssb_fruit_juice,
    REDPROC_MEAT_SERV_AHEI = AHEI_VALIDATION$red_processed_meat,
    TRANS_SERV_AHEI = AHEI_VALIDATION$trans_fat,
    SODIUM_SERV_AHEI = AHEI_VALIDATION$sodium,
    ALCOHOL_SERV_AHEI = AHEI_VALIDATION$alcohol
    )

# Merge the validation data with the result
AHEI_VALIDATION_RESULT_FINAL = AHEI_VALIDATION %>%
    left_join(AHEI_VALIDATION_RESULT, by = c("id" = "RESPONDENTID"))

# Save the result
write_csv(AHEI_VALIDATION_RESULT_FINAL, "AHEI_validation_result.csv")

## HEI2015 validation
HEI2015_VALIDATION_RESULT = HEI2015(
    SERV_DATA = HEI2015_VALIDATION,
    RESPONDENTID = HEI2015_VALIDATION$id,
    TOTALKCAL_HEI2015 = HEI2015_VALIDATION$kcal,
    TOTALFRT_SERV_HEI2015 = HEI2015_VALIDATION$total_fruit,
    FRT_SERV_HEI2015 = HEI2015_VALIDATION$whole_fruit,
    VEG_SERV_HEI2015 = HEI2015_VALIDATION$total_vegetable,
    GREENNBEAN_SERV_HEI2015 = HEI2015_VALIDATION$green_and_bean,
    TOTALPRO_SERV_HEI2015 = HEI2015_VALIDATION$total_protein,
    SEAPLANTPRO_SERV_HEI2015 = HEI2015_VALIDATION$seafood_plant_protein,
    WHOLEGRAIN_SERV_HEI2015 = HEI2015_VALIDATION$whole_grain,
    DAIRY_SERV_HEI2015 = HEI2015_VALIDATION$dairy,
    FATTYACID_SERV_HEI2015 = HEI2015_VALIDATION$fatty_acid,
    REFINEDGRAIN_SERV_HEI2015 = HEI2015_VALIDATION$refined_grain,
    SODIUM_SERV_HEI2015 = HEI2015_VALIDATION$sodium,
    ADDEDSUGAR_SERV_HEI2015 = HEI2015_VALIDATION$added_sugar,
    SATFAT_SERV_HEI2015 = HEI2015_VALIDATION$saturated_fat
)

# Merge the validation data with the result
HEI2015_VALIDATION_RESULT_FINAL = HEI2015_VALIDATION %>%
    left_join(HEI2015_VALIDATION_RESULT, by = c("id" = "RESPONDENTID"))

# Save the merged result
write_csv(HEI2015_VALIDATION_RESULT_FINAL, "HEI2015_validation_result.csv")

## HEI2020 validation
HEI2020_VALIDATION_RESULT = HEI2020(
    SERV_DATA = HEI2020_VALIDATION,
    RESPONDENTID = HEI2020_VALIDATION$id,
    AGE = HEI2020_VALIDATION$age,
    TOTALKCAL_HEI2020 = HEI2020_VALIDATION$kcal,
    TOTALFRT_SERV_HEI2020 = HEI2020_VALIDATION$total_fruit,
    FRT_SERV_HEI2020 = HEI2020_VALIDATION$whole_fruit,
    VEG_SERV_HEI2020 = HEI2020_VALIDATION$total_vegetable,
    GREENNBEAN_SERV_HEI2020 = HEI2020_VALIDATION$green_and_bean,
    TOTALPRO_SERV_HEI2020 = HEI2020_VALIDATION$total_protein,
    SEAPLANTPRO_SERV_HEI2020 = HEI2020_VALIDATION$seafood_plant_protein,
    WHOLEGRAIN_SERV_HEI2020 = HEI2020_VALIDATION$whole_grain,
    DAIRY_SERV_HEI2020 = HEI2020_VALIDATION$dairy,
    FATTYACID_SERV_HEI2020 = HEI2020_VALIDATION$fatty_acid,
    REFINEDGRAIN_SERV_HEI2020 = HEI2020_VALIDATION$refined_grain,
    SODIUM_SERV_HEI2020 = HEI2020_VALIDATION$sodium,
    ADDEDSUGAR_SERV_HEI2020 = HEI2020_VALIDATION$added_sugar,
    SATFAT_SERV_HEI2020 = HEI2020_VALIDATION$saturated_fat
)

HEI2020_VALIDATION_RESULT_FINAL = HEI2020_VALIDATION %>%
    left_join(HEI2020_VALIDATION_RESULT, by = c("id" = "RESPONDENTID"))

# Save the merged result
write_csv(HEI2020_VALIDATION_RESULT_FINAL, "HEI2020_validation_result.csv")

## AHEIP validation
AHEIP_VALIDATION_RESULT = AHEIP(
    SERV_DATA = AHEIP_VALIDATION,
    RESPONDENTID = AHEIP_VALIDATION$id,
    VEG_SERV_AHEIP = AHEIP_VALIDATION$vegetable,
    FRT_SERV_AHEIP = AHEIP_VALIDATION$whole_fruit,
    WHITERED_RT_SERV_AHEIP = AHEIP_VALIDATION$white_meat_red_meat,
    FIBER_SERV_AHEIP = AHEIP_VALIDATION$fiber,
    TRANS_SERV_AHEIP = AHEIP_VALIDATION$trans_fat,
    POLYSAT_RT_SERV_AHEIP = AHEIP_VALIDATION$poly_fat_sat_fat,
    CALCIUM_SERV_AHEIP = AHEIP_VALIDATION$calcium,
    FOLATE_SERV_AHEIP = AHEIP_VALIDATION$folate,
    IRON_SERV_AHEIP = AHEIP_VALIDATION$iron
)

# Merge the validation data with the result
AHEIP_VALIDATION_RESULT_FINAL = AHEIP_VALIDATION %>%
    left_join(AHEIP_VALIDATION_RESULT, by = c("id" = "RESPONDENTID"))

# Save the merged result
write_csv(AHEIP_VALIDATION_RESULT_FINAL, "AHEIP_validation_result.csv")

## DASH validation
DASH_VALIDATION_RESULT = DASH(
    SERV_DATA = DASH_VALIDATION,
    RESPONDENTID = DASH_VALIDATION$id,
    TOTALKCAL_DASH = DASH_VALIDATION$kcal,
    FRT_FRTJ_SERV_DASH = DASH_VALIDATION$whole_fruit,
    VEG_SERV_DASH = DASH_VALIDATION$vegetable,
    NUTSLEG_SERV_DASH = DASH_VALIDATION$nut_legume,
    WGRAIN_SERV_DASH = DASH_VALIDATION$whole_grain,
    LOWF_DAIRY_SERV_DASH = DASH_VALIDATION$low_fat_dairy,
    SODIUM_SERV_DASH = DASH_VALIDATION$sodium,
    REDPROC_MEAT_SERV_DASH = DASH_VALIDATION$red_processed_meat,
    SSB_FRTJ_SERV_DASH = DASH_VALIDATION$ssb
)

# Merge the validation data with the result
DASH_VALIDATION_RESULT_FINAL = DASH_VALIDATION %>%
    left_join(DASH_VALIDATION_RESULT, by = c("id" = "RESPONDENTID"))

# Save the merged result
write_csv(DASH_VALIDATION_RESULT_FINAL, "DASH_validation_result.csv")

## DASHI validation
DASHI_VALIDATION_RESULT = DASHI(
    SERV_DATA = DASHI_VALIDATION, 
    RESPONDENTID = DASHI_VALIDATION$id, 
    TOTALKCAL_DASHI = DASHI_VALIDATION$kcal, 
    TOTAL_FAT_DASHI = DASHI_VALIDATION$total_fat, 
    SAT_FAT_DASHI = DASHI_VALIDATION$sat_fat, 
    PROTEIN_DASHI = DASHI_VALIDATION$protein, 
    CHOLESTEROL_DASHI = DASHI_VALIDATION$cholesterol, 
    FIBER_DASHI = DASHI_VALIDATION$fiber, 
    POTASSIUM_DASHI = DASHI_VALIDATION$potassium, 
    MAGNESIUM_DASHI = DASHI_VALIDATION$magnesium,
    CALCIUM_DASHI = DASHI_VALIDATION$calcium, 
    SODIUM_DASHI = DASHI_VALIDATION$sodium
)

# Merge the validation data with the result
DASHI_VALIDATION_RESULT_FINAL = DASHI_VALIDATION %>%
    left_join(DASHI_VALIDATION_RESULT, by = c("id" = "RESPONDENTID"))

# Save the merged result
write_csv(DASHI_VALIDATION_RESULT_FINAL, "DASHI_validation_result.csv")

## MED validation
MED_VALIDATION_RESULT = MED(
    SERV_DATA = MED_VALIDATION, 
    RESPONDENTID = MED_VALIDATION$id, 
    FRT_FRTJ_SERV_MED = MED_VALIDATION$fruit, 
    VEG_SERV_MED = MED_VALIDATION$vegetable, 
    WGRAIN_SERV_MED = MED_VALIDATION$whole_grain, 
    LEGUMES_SERV_MED = MED_VALIDATION$legume, 
    NUTS_SERV_MED = MED_VALIDATION$nut, 
    FISH_SERV_MED = MED_VALIDATION$fish, 
    REDPROC_MEAT_SERV_MED = MED_VALIDATION$red_processed_meat, 
    MONSATFAT_SERV_MED = MED_VALIDATION$monofat_satfat, 
    ALCOHOL_SERV_MED = MED_VALIDATION$alcohol)

# Merge the validation data with the result
MED_VALIDATION_RESULT_FINAL = MED_VALIDATION %>%
    left_join(MED_VALIDATION_RESULT, by = c("id" = "RESPONDENTID"))

# Save the result
write_csv(MED_VALIDATION_RESULT_FINAL, "MED_VALIDATION_RESULT_FINAL.csv")

## MEDI validation
MEDI_VALIDATION_RESULT = MEDI(
    SERV_DATA = MEDI_VALIDATION, 
    RESPONDENTID = MEDI_VALIDATION$id, 
    OLIVE_OIL_SERV_MEDI = MEDI_VALIDATION$olive_oil, 
    VEG_SERV_MEDI = MEDI_VALIDATION$vegetable, 
    FRT_SERV_MEDI = MEDI_VALIDATION$fruit, 
    LEGUMES_SERV_MEDI = MEDI_VALIDATION$legume, 
    NUTS_SERV_MEDI = MEDI_VALIDATION$nut, 
    FISH_SEAFOOD_SERV_MEDI = MEDI_VALIDATION$fish, 
    ALCOHOL_SERV_MEDI = MEDI_VALIDATION$alcohol, 
    SSB_SERV_MEDI = MEDI_VALIDATION$ssb, 
    SWEETS_SERV_MEDI = MEDI_VALIDATION$sweets, 
    DISCRET_FAT_SERV_MEDI = MEDI_VALIDATION$discret_fat, 
    REDPROC_MEAT_SERV_MEDI = MEDI_VALIDATION$red_meat)

# Merge the validation data with the result
MEDI_VALIDATION_RESULT_FINAL = MEDI_VALIDATION %>%
    left_join(MEDI_VALIDATION_RESULT, by = c("id" = "RESPONDENTID"))

# Save the result
write_csv(MEDI_VALIDATION_RESULT_FINAL, "MEDI_validation_result.csv")

## MEDI_V2 validation
MEDI_V2_VALIDATION_RESULT = MEDI_V2(
    SERV_DATA = MEDI_V2_VALIDATION, 
    RESPONDENTID = MEDI_V2_VALIDATION$id, 
    OLIVE_OIL_SERV_MEDI = MEDI_V2_VALIDATION$olive_oil, 
    VEG_SERV_MEDI = MEDI_V2_VALIDATION$vegetable, 
    FRT_SERV_MEDI = MEDI_V2_VALIDATION$fruit, 
    LEGUMES_SERV_MEDI = MEDI_V2_VALIDATION$legume, 
    NUTS_SERV_MEDI = MEDI_V2_VALIDATION$nut, 
    FISH_SEAFOOD_SERV_MEDI = MEDI_V2_VALIDATION$fish, 
    ALCOHOL_SERV_MEDI = MEDI_V2_VALIDATION$alcohol, 
    SSB_SERV_MEDI = MEDI_V2_VALIDATION$ssb, 
    SWEETS_SERV_MEDI = MEDI_V2_VALIDATION$sweets, 
    DISCRET_FAT_SERV_MEDI = MEDI_V2_VALIDATION$discret_fat, 
    REDPROC_MEAT_SERV_MEDI = MEDI_V2_VALIDATION$red_meat)

# Merge the validation data with the result
MEDI_V2_VALIDATION_RESULT_FINAL = MEDI_V2_VALIDATION %>%
    left_join(MEDI_V2_VALIDATION_RESULT, by = c("id" = "RESPONDENTID"))

# Save the result
write_csv(MEDI_V2_VALIDATION_RESULT_FINAL, "MEDI_V2_validation_result.csv")

## PHDI validation
PHDI_VALIDATION_RESULT = PHDI(
    SERV_DATA=PHDI_VALIDATION, 
    PHDI_VALIDATION$id, 
    PHDI_VALIDATION$gender, 
    PHDI_VALIDATION$TOTALKCAL_PHDI, 
    PHDI_VALIDATION$WGRAIN_SERV_PHDI, 
    PHDI_VALIDATION$STARCHY_VEG_SERV_PHDI, 
    PHDI_VALIDATION$VEG_SERV_PHDI, 
    PHDI_VALIDATION$FRT_SERV_PHDI, 
    PHDI_VALIDATION$DAIRY_SERV_PHDI, 
    PHDI_VALIDATION$REDPROC_MEAT_SERV_PHDI, 
    PHDI_VALIDATION$POULTRY_SERV_PHDI, 
    PHDI_VALIDATION$EGG_SERV_PHDI, 
    PHDI_VALIDATION$FISH_SERV_PHDI, 
    PHDI_VALIDATION$NUTS_SERV_PHDI, 
    PHDI_VALIDATION$LEGUMES_SERV_PHDI, 
    PHDI_VALIDATION$SOY_SERV_PHDI, 
    PHDI_VALIDATION$ADDED_FAT_UNSAT_SERV_PHDI, 
    PHDI_VALIDATION$ADDED_FAT_SAT_TRANS_SERV_PHDI, 
    PHDI_VALIDATION$ADDED_SUGAR_SERV_PHDI
    ) 

# Merge the validation data with the result
PHDI_VALIDATION_RESULT_FINAL = PHDI_VALIDATION %>%
    left_join(PHDI_VALIDATION_RESULT, by = c("id" = "RESPONDENTID"))

# Save the result
write_csv(PHDI_VALIDATION_RESULT_FINAL, "PHDI_validation_result.csv")

## DII validation
DII_VALIDATION_RESULT = DII(SERV_DATA = DII_VALIDATION, RESPONDENTID = DII_VALIDATION$id, REPEATNUM = 1, ALCOHOL_DII = DII_VALIDATION$Alcohol, VITB12_DII = DII_VALIDATION$`vitamin B12`, VITB6_DII = DII_VALIDATION$`vitamin B6`, BCAROTENE_DII = DII_VALIDATION$`Beta-carotene`, CAFFEINE_DII = DII_VALIDATION$Caffeine, CARB_DII = DII_VALIDATION$Carbohydrate, CHOLES_DII = DII_VALIDATION$Cholesterol, KCAL_DII = DII_VALIDATION$Energy, EUGENOL_DII = DII_VALIDATION$Eugenol, TOTALFAT_DII = DII_VALIDATION$`Total fat`, FIBER_DII = DII_VALIDATION$Fiber, FOLICACID_DII = DII_VALIDATION$`Folic acid`, GARLIC_DII = DII_VALIDATION$Garlic, GINGER_DII = DII_VALIDATION$Ginger, IRON_DII = DII_VALIDATION$Iron, MG_DII = DII_VALIDATION$Magnesium, MUFA_DII = DII_VALIDATION$MUFA, NIACIN_DII = DII_VALIDATION$Niacin, N3FAT_DII = DII_VALIDATION$`n-3 fatty acid`, N6FAT_DII = DII_VALIDATION$`n-6 fatty acid`, ONION_DII = DII_VALIDATION$Onion, PROTEIN_DII = DII_VALIDATION$Protein, PUFA_DII = DII_VALIDATION$PUFA, RIBOFLAVIN_DII = DII_VALIDATION$Riboflavin, SAFFRON_DII = DII_VALIDATION$Saffron, SATFAT_DII = DII_VALIDATION$`Saturated fat`, SE_DII = DII_VALIDATION$Selenium, THIAMIN_DII = DII_VALIDATION$Thiamin, TRANSFAT_DII = DII_VALIDATION$`Trans fat`, TURMERIC_DII = DII_VALIDATION$Turmeric, VITA_DII = DII_VALIDATION$`Vitamin A`, VITC_DII = DII_VALIDATION$`Vitamin C`, VITD_DII = DII_VALIDATION$`Vitamin D`, VITE_DII = DII_VALIDATION$`Vitamin E`, ZN_DII = DII_VALIDATION$Zinc, TEA_DII = DII_VALIDATION$`Green/black tea`, FLA3OL_DII = DII_VALIDATION$`Flavan-3-ol`, FLAVONES_DII = DII_VALIDATION$Flavones, FLAVONOLS_DII = DII_VALIDATION$Flavonols, FLAVONONES_DII = DII_VALIDATION$Flavonones, ANTHOC_DII = DII_VALIDATION$Anthocyanidins, ISOFLAVONES_DII = DII_VALIDATION$`Isoflavones`, PEPPER_DII = DII_VALIDATION$Pepper, THYME_DII = DII_VALIDATION$Thyme_oregano, ROSEMARY_DII = DII_VALIDATION$Rosemary)

# Merge the validation data with the result
DII_VALIDATION_RESULT_FINAL = DII_VALIDATION %>%
    left_join(DII_VALIDATION_RESULT, by = c("id" = "RESPONDENTID"))

# Save the result
write_csv(DII_VALIDATION_RESULT_FINAL, "DII_validation_result.csv")
```
