## Supplement 3 Validation Figures for "Dietaryindex: A User-Friendly and Versatile R Package for Standardizing Dietary Pattern Analysis in Epidemiological and Clinical Studies": Summary Validation Figures.docx

Supplementary Material 2: Validation Figures

Supplementary Figure 1. ACS2020_V1 Accuracy

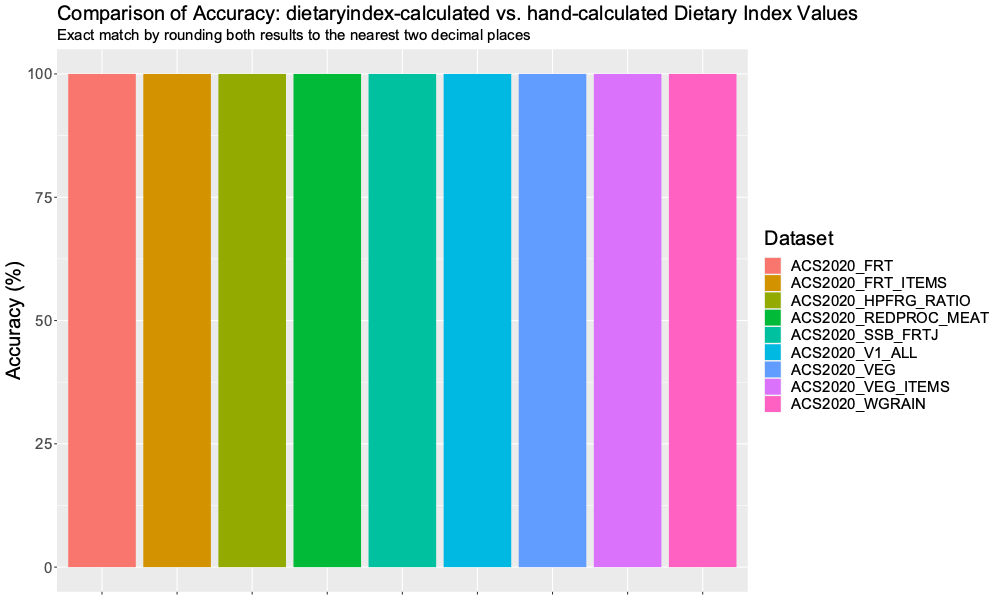

Supplementary Figure 2. ACS2020_V2 Accuracy

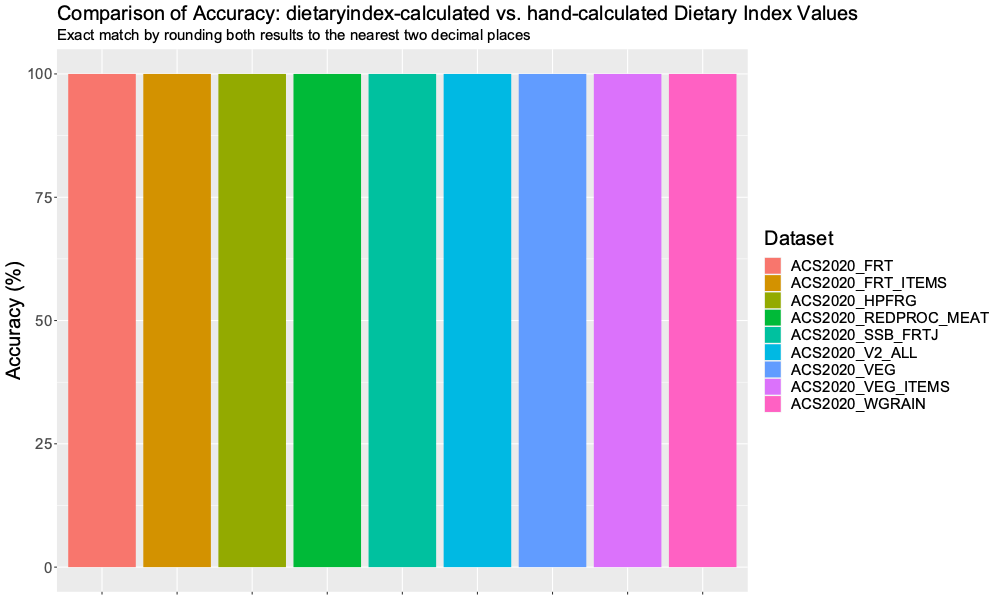

Supplementary Figure 3. AHEI Accuracy

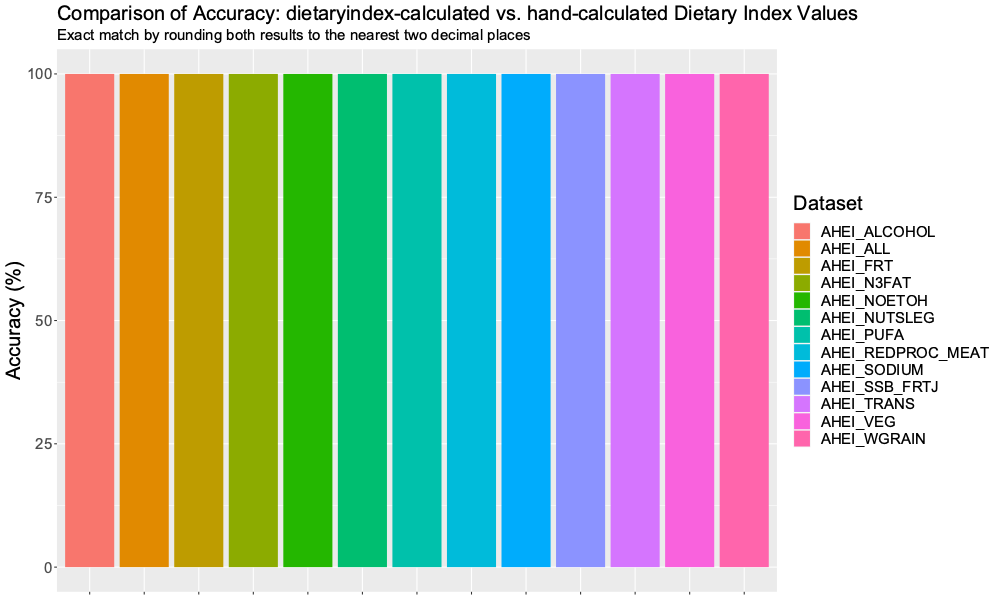

Supplementary Figure 4. AHEIP Accuracy

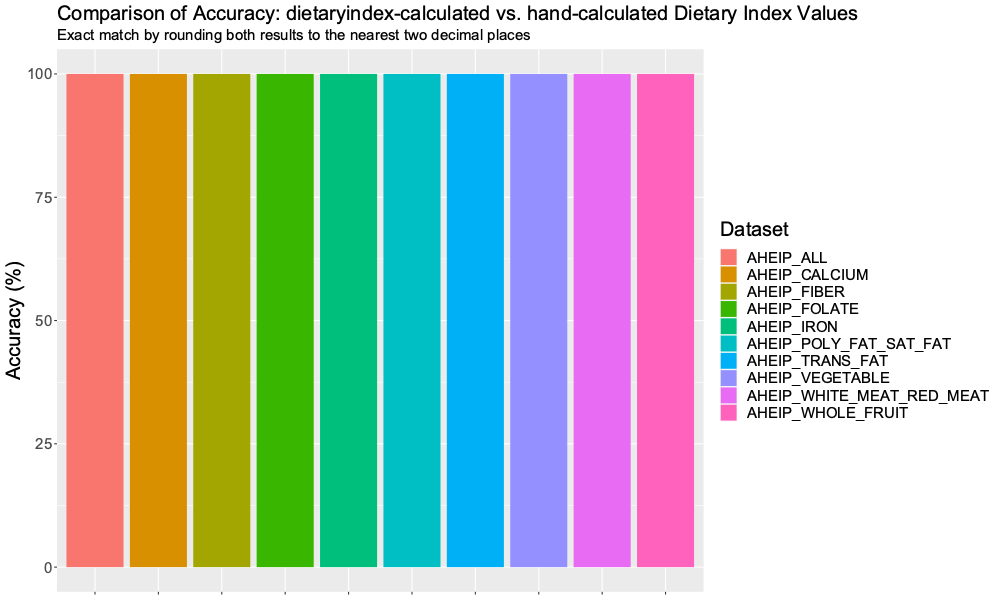

Supplementary Figure 5. DASH Accuracy

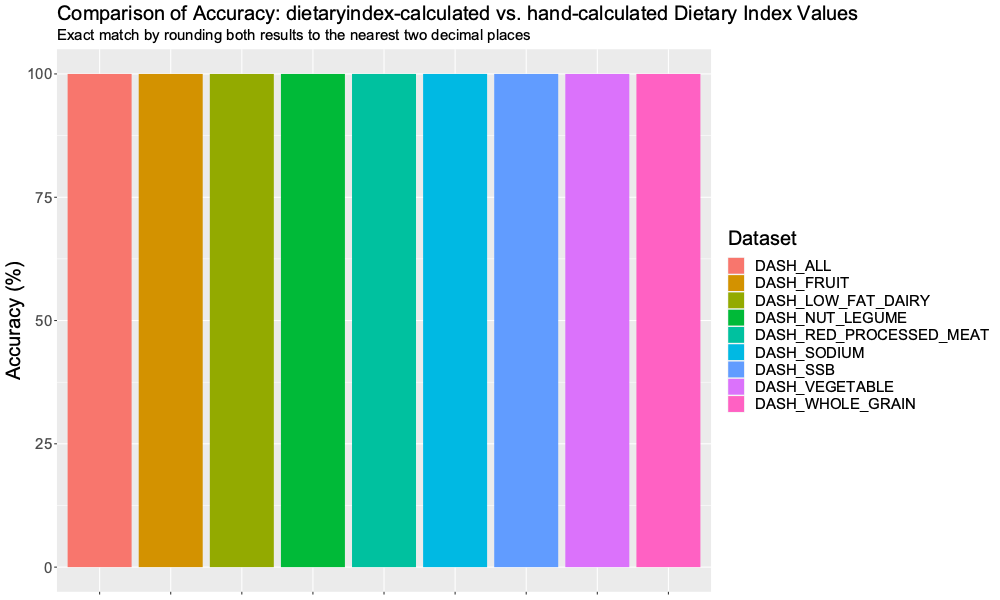

Supplementary Figure 6. DASHI Accuracy

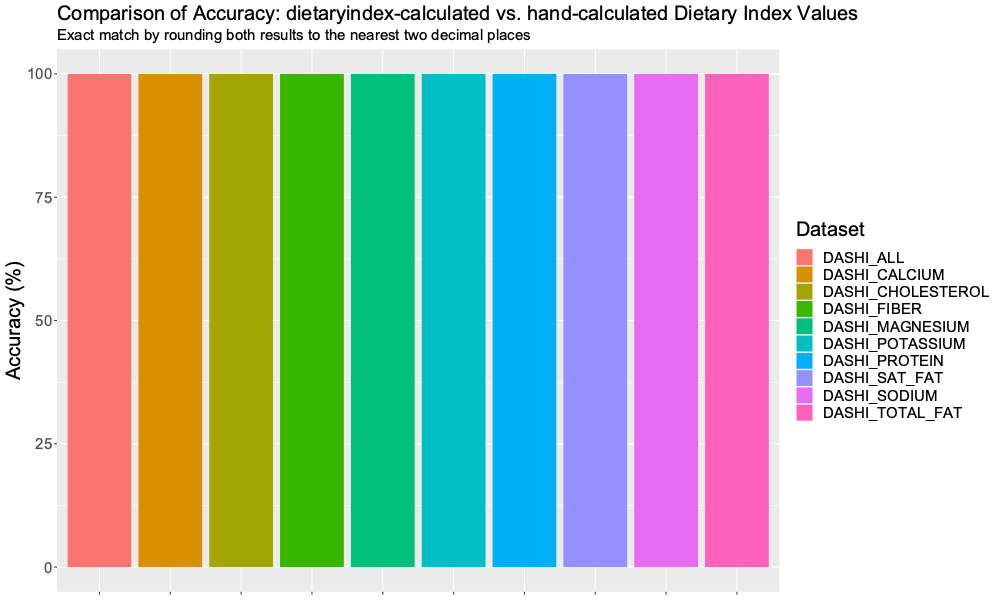

Supplementary Figure 7. DII Accuracy

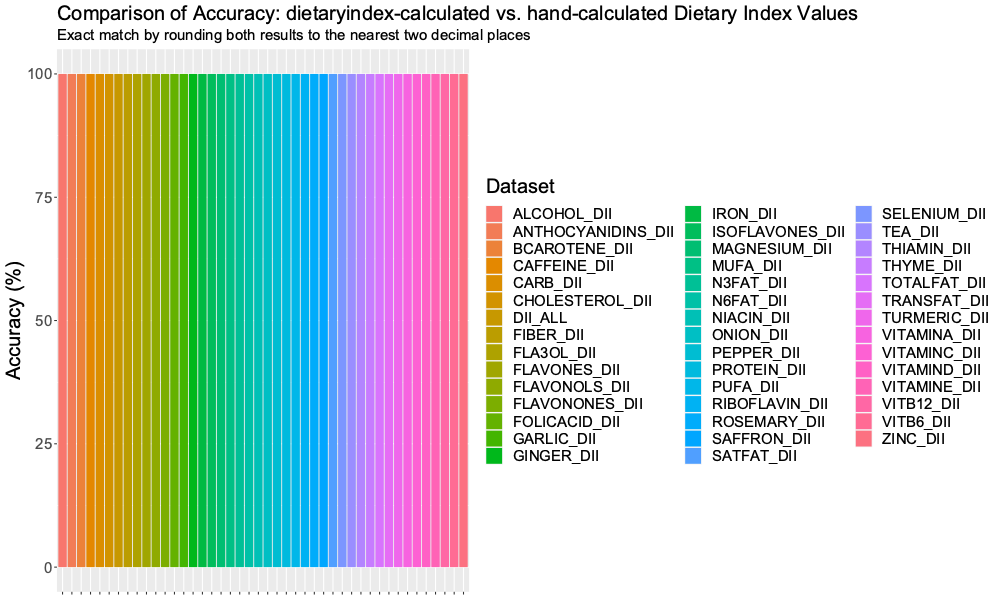

Supplementary Figure 8. HEI2015 Accuracy

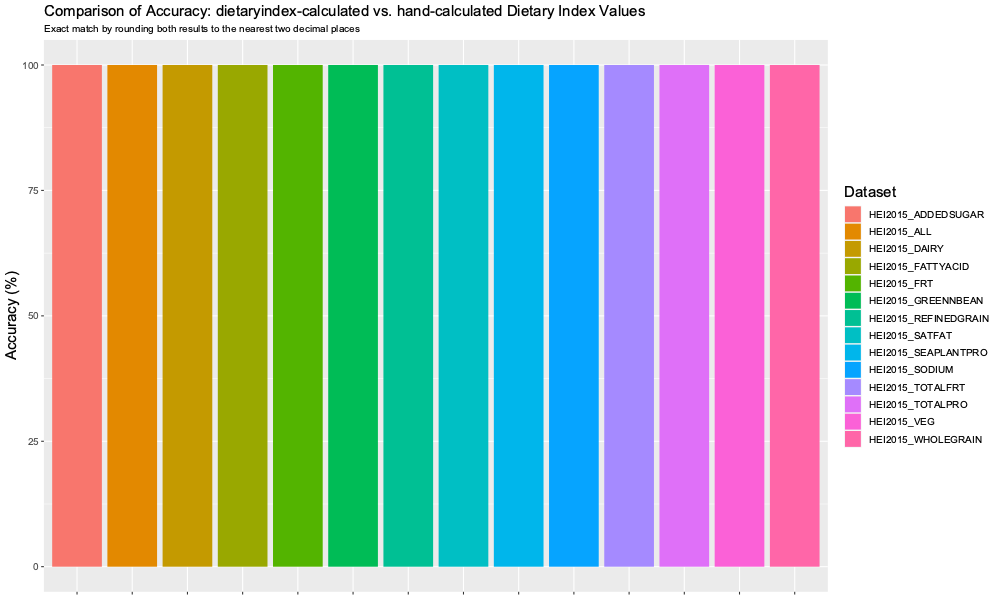

Supplementary Figure 9. HEI2020 Accuracy

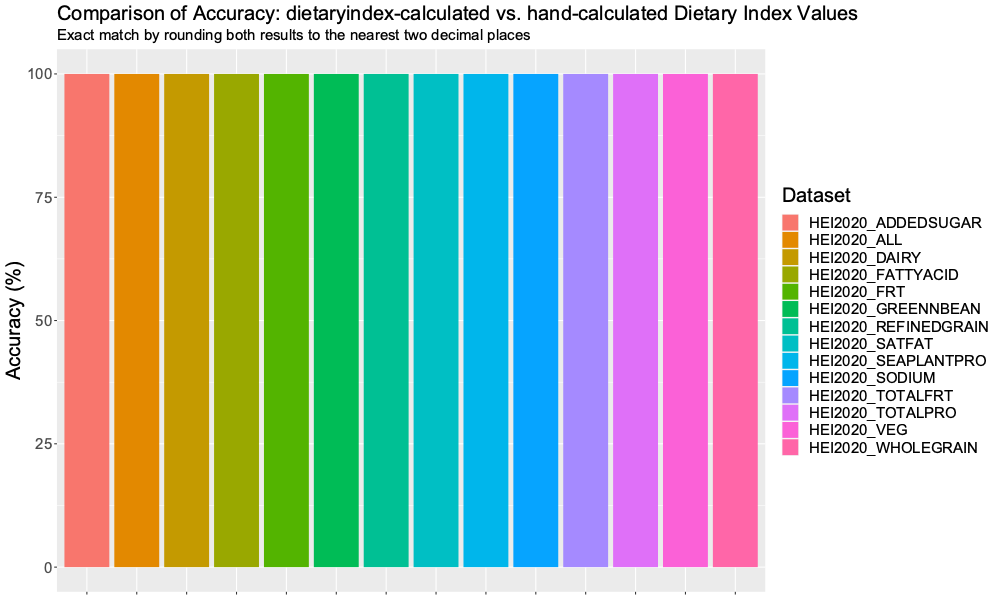

Supplementary Figure 10. MED Accuracy

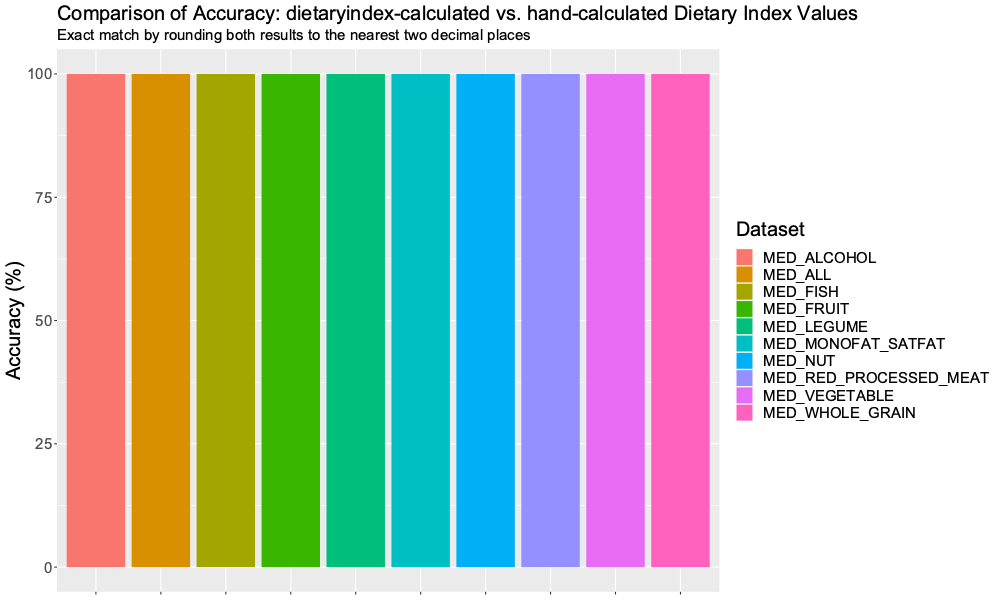

Supplementary Figure 11. MEDI Accuracy

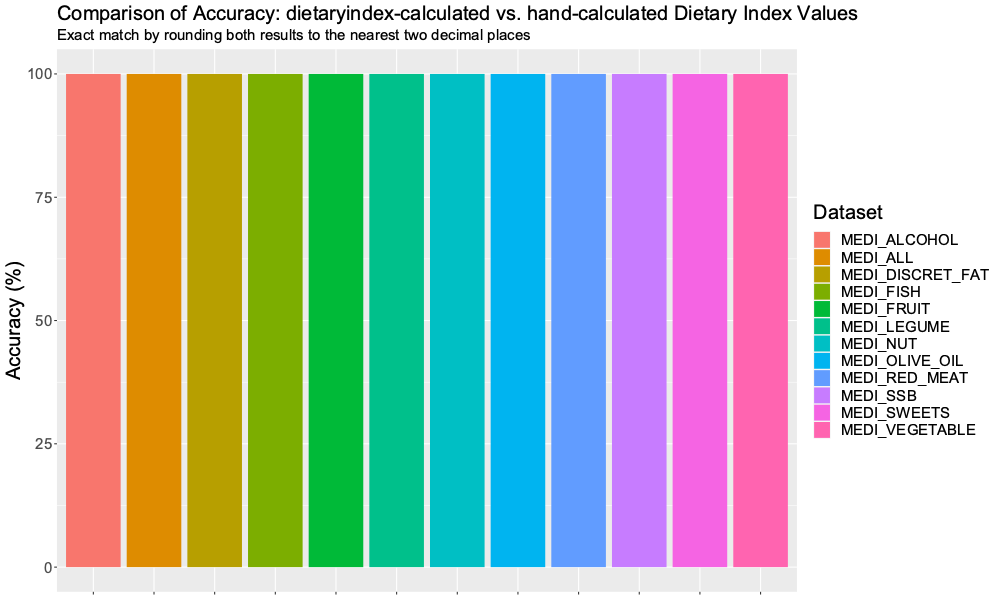

Supplementary Figure 12. MEDI_V2 Accuracy

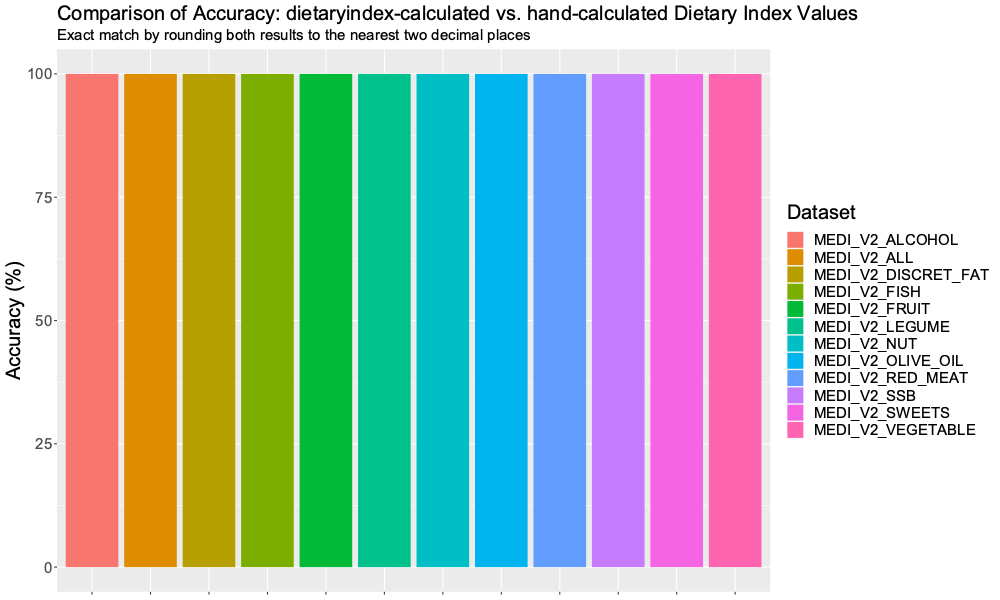

Supplementary Figure 13. PHDI Accuracy

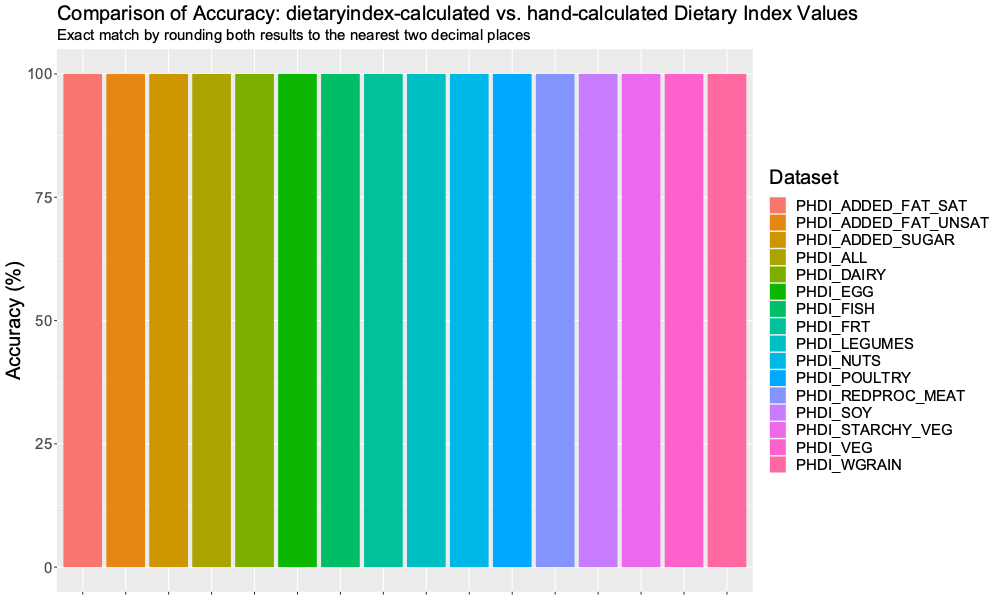
