## Supplement 3 Validation Figures for "Dietaryindex: A User-Friendly and Versatile R Package for Standardizing Dietary Pattern Analysis in Epidemiological and Clinical Studies": Validation figures.r

```
# produce validation graph

# set up working directory
setwd("/Users/james/Desktop/Emory University - Ph.D./dietaryindex_package/dietaryindex/Validation file for publication/Final validation files")
library(ggplot2)
library(dplyr)
library(readr)

# read in dietaryindex-calculated and hand-calculated validation results
ACS2020_V1_validation_result = read_csv("ACS2020_V1_validation_result.csv")
ACS2020_V2_validation_result = read_csv("ACS2020_V2_validation_result.csv")
AHEI_validation_result = read_csv("AHEI_validation_result.csv")
AHEIP_validation_result = read_csv("AHEIP_validation_result.csv")
DASH_validation_result = read_csv("DASH_validation_result.csv")
DASHI_validation_result = read_csv("DASHI_validation_result.csv")
DII_validation_result = read_csv("DII_validation_result.csv")
HEI2015_validation_result = read_csv("HEI2015_validation_result.csv")
HEI2020_validation_result = read_csv("HEI2020_validation_result.csv")
MED_validation_result = read_csv("MED_validation_result.csv")
MEDI_validation_result = read_csv("MEDI_validation_result.csv")
MEDI_V2_validation_result = read_csv("MEDI_V2_validation_result.csv")
PHDI_validation_result = read_csv("PHDI_validation_result.csv")

# read in NHANES NCI SAS and dietaryindex-calculated validation results
setwd("/Users/james/Desktop/Emory University - Ph.D./dietaryindex_package/dietaryindex/Validation file for publication/HEI2015_NHANES_1718")
SAS_HEI2015_1718 = read_csv("SAS_HEI2015_1718.csv")
dietaryindex_HEI2015_1718 = read_csv("dietaryindex_HEI2015_1718.csv")

setwd("/Users/james/Desktop/Emory University - Ph.D./dietaryindex_package/dietaryindex/Validation file for publication/HEI2015_ASA24_example_data")
HEI2015_ASA24_NCI_SAS = read_csv("HEI2015_ASA24_NCI_SAS.csv")
HEI2015_ASA24_dietaryindex = read_csv("HEI2015_ASA24_dietaryindex.csv")

setwd("/Users/james/Desktop/Emory University - Ph.D./dietaryindex_package/dietaryindex/Validation file for publication/HEI2015_DHQ3_example_data")
HEI2015_DHQ3_NCI_SAS = read_csv("Sample total daily results.csv", skip = 1)
HEI2015_DHQ3_dietaryindex = read_csv("HEI2015_DHQ3_dietaryindex.csv")

# Define a function to compute accuracy
get_accuracy <- function(x, y) {
  # round x to 2 decimal places
  x <- round(x, 2)
  # round y to 2 decimal places
  y <- round(y, 2)
  return ((sum(x == y, na.rm = TRUE) / length(x)) * 100)
}

# Define a function to compute accuracy
get_accuracy_diff <- function(x, y) {
  # the maximum tolerance for the difference between x and y
  tolerance <- 0.5
  # Subtract data1 from data2 (or vice versa) and take the absolute value of the differences
  diff <- abs(x - y)
  return ((sum(diff <= tolerance, na.rm = TRUE) / length(x)) * 100)
}

############### Total score of all dietary indexes validation ###############

# Initialize a data frame to store results for the total dietary index score for all dietary indexes
results <- data.frame(
  Dataset = character(),
  Accuracy = numeric(),
  stringsAsFactors = FALSE
)

# List of all your datasets and their corresponding column names
datasets <- list(
  ACS2020_V1 = list(data = ACS2020_V1_validation_result, cols = c("EXP_ACS_ALL", "ACS2020_V1_ALL")),
  ACS2020_V2 = list(data = ACS2020_V2_validation_result, cols = c("EXP_ACS_ALL", "ACS2020_V2_ALL")),
  AHEI = list(data = AHEI_validation_result, cols = c("EXP_AHEI_ALL", "AHEI_ALL")),
  AHEIP = list(data = AHEIP_validation_result, cols = c("EXP_AHEIP_ALL", "AHEIP_ALL")),
  DASH = list(data = DASH_validation_result, cols = c("EXP_DASH_ALL", "DASH_ALL")),
  DASHI = list(data = DASHI_validation_result, cols = c("EXP_DASHI_ALL", "DASHI_ALL")),
  DII = list(data = DII_validation_result, cols = c("EXP_DII_ALL", "DII_ALL")),
  HEI2015 = list(data = HEI2015_validation_result, cols = c("EXP_HEI2015_ALL", "HEI2015_ALL")),
  HEI2020 = list(data = HEI2020_validation_result, cols = c("EXP_HEI2020_ALL", "HEI2020_ALL")),
  MED = list(data = MED_validation_result, cols = c("EXP_MED_ALL", "MED_ALL")),
  MEDI = list(data = MEDI_validation_result, cols = c("EXP_MEDI_ALL", "MEDI_ALL")),
  MEDI_V2 = list(data = MEDI_V2_validation_result, cols = c("EXP_MEDI_ALL", "MEDI_V2_ALL")),
  PHDI = list(data = PHDI_validation_result, cols = c("EXP_PHDI_ALL", "PHDI_ALL"))
)

# Compute accuracy for each dataset
for (name in names(datasets)) {
  dataset <- datasets[[name]]
  accuracy <- get_accuracy(dataset$data[[dataset$cols[1]]], dataset$data[[dataset$cols[2]]])
  results <- rbind(results, data.frame(Dataset = name, Accuracy = accuracy))
}

# Print results
print(results)


# Plot results
ggplot(results, aes(x = Dataset, y = Accuracy, fill=Dataset)) +
    geom_bar(stat = "identity") +
    ylab("Accuracy (%)") +
    xlab(NULL) +
    ggtitle("Comparison of Accuracy: dietaryindex-calculated vs. hand-calculated Dietary Index Values") +
    # add a subtitie
    labs(subtitle = "Exact match by rounding both results to the nearest two decimal places") +
    # increase the title size
    theme(
        plot.title = element_text(size = 20),
        plot.subtitle = element_text(size = 15),
        axis.title = element_text(size = 20),
        axis.text = element_text(size = 15),
        legend.text = element_text(size = 15),
        legend.title = element_text(size = 20),
        axis.text.x = element_blank()
        ) 

######## ACS2020_V1 validation ##########

# Initialize a data frame to store results for all dietary index component scores in ACS2020_V1
results_ACS2020_V1 <- data.frame(
  Dataset = character(),
  Accuracy = numeric(),
  stringsAsFactors = FALSE
)

# List of all your datasets and their corresponding column names
datasets_ACS2020_V1 <- list(
    ACS2020_V1_ALL = list(data = ACS2020_V1_validation_result, cols = c("EXP_ACS_ALL", "ACS2020_V1_ALL")),
    ACS2020_VEG = list(data = ACS2020_V1_validation_result, cols = c("EXP_ACS_VEGETABLE", "ACS2020_VEG")),
    ACS2020_VEG_ITEMS = list(data = ACS2020_V1_validation_result, cols = c("EXP_ACS_VEGETABLE_UNIQUE", "ACS2020_VEG_ITEMS")),
    ACS2020_FRT = list(data = ACS2020_V1_validation_result, cols = c("EXP_ACS_FRUIT", "ACS2020_FRT")),
    ACS2020_FRT_ITEMS = list(data = ACS2020_V1_validation_result, cols = c("EXP_ACS_FRUIT_UNIQUE", "ACS2020_FRT_ITEMS")),
    ACS2020_WGRAIN = list(data = ACS2020_V1_validation_result, cols = c("EXP_ACS_WHOLE_GRAIN", "ACS2020_WGRAIN")),
    ACS2020_REDPROC_MEAT = list(data = ACS2020_V1_validation_result, cols = c("EXP_ACS_RED_MEAT", "ACS2020_REDPROC_MEAT")),
    ACS2020_HPFRG_RATIO = list(data = ACS2020_V1_validation_result, cols = c("EXP_ACS_PROCESSED_FOOD", "ACS2020_HPFRG_RATIO")),
    ACS2020_SSB_FRTJ = list(data = ACS2020_V1_validation_result, cols = c("EXP_ACS_SSB", "ACS2020_SSB_FRTJ"))
)

# Compute accuracy for each dataset
for (name in names(datasets_ACS2020_V1)) {
    dataset <- datasets_ACS2020_V1[[name]]
    accuracy <- get_accuracy(dataset$data[[dataset$cols[1]]], dataset$data[[dataset$cols[2]]])
    results_ACS2020_V1 <- rbind(results_ACS2020_V1, data.frame(Dataset = name, Accuracy = accuracy))
}

# Print results
print(results_ACS2020_V1)

# Plot results
ggplot(results_ACS2020_V1, aes(x = Dataset, y = Accuracy, fill=Dataset)) +
    geom_bar(stat = "identity") +
    ylab("Accuracy (%)") +
    xlab(NULL) +
    ggtitle("Comparison of Accuracy: dietaryindex-calculated vs. hand-calculated Dietary Index Values") +
    # add a subtitie
    labs(subtitle = "Exact match by rounding both results to the nearest two decimal places") +
    # increase the title size
    theme(
        plot.title = element_text(size = 20),
        plot.subtitle = element_text(size = 15),
        axis.title = element_text(size = 20),
        axis.text = element_text(size = 15),
        legend.text = element_text(size = 15),
        legend.title = element_text(size = 20),
        axis.text.x = element_blank()
        )

######### ACS2020_V2 validation ##########

# Initialize a data frame to store results for all dietary index component scores in ACS2020_V2
results_ACS2020_V2 <- data.frame(
  Dataset = character(),
  Accuracy = numeric(),
  stringsAsFactors = FALSE
)

# List of all your datasets and their corresponding column names for ACS2020_V2
datasets_ACS2020_V2 <- list(
    ACS2020_V2_ALL = list(data = ACS2020_V2_validation_result, cols = c("EXP_ACS_ALL", "ACS2020_V2_ALL")),
    ACS2020_VEG = list(data = ACS2020_V2_validation_result, cols = c("EXP_ACS_VEGETABLE", "ACS2020_VEG")),
    ACS2020_VEG_ITEMS = list(data = ACS2020_V2_validation_result, cols = c("EXP_ACS_VEGETABLE_UNIQUE", "ACS2020_VEG_ITEMS")),
    ACS2020_FRT = list(data = ACS2020_V2_validation_result, cols = c("EXP_ACS_FRUIT", "ACS2020_FRT")),
    ACS2020_FRT_ITEMS = list(data = ACS2020_V2_validation_result, cols = c("EXP_ACS_FRUIT_UNIQUE", "ACS2020_FRT_ITEMS")),
    ACS2020_WGRAIN = list(data = ACS2020_V2_validation_result, cols = c("EXP_ACS_WHOLE_GRAIN", "ACS2020_WGRAIN")),
    ACS2020_REDPROC_MEAT = list(data = ACS2020_V2_validation_result, cols = c("EXP_ACS_RED_MEAT", "ACS2020_REDPROC_MEAT")),
    ACS2020_HPFRG = list(data = ACS2020_V2_validation_result, cols = c("EXP_ACS_PROCESSED_FOOD", "ACS2020_HPFRG")),
    ACS2020_SSB_FRTJ = list(data = ACS2020_V2_validation_result, cols = c("EXP_ACS_SSB", "ACS2020_SSB_FRTJ"))
)

# Compute accuracy for each dataset in ACS2020_V2
for (name in names(datasets_ACS2020_V2)) {
    dataset <- datasets_ACS2020_V2[[name]]
    accuracy <- get_accuracy(dataset$data[[dataset$cols[1]]], dataset$data[[dataset$cols[2]]])
    results_ACS2020_V2 <- rbind(results_ACS2020_V2, data.frame(Dataset = name, Accuracy = accuracy))
}

# Print results
print(results_ACS2020_V2)

# Plot results
ggplot(results_ACS2020_V2, aes(x = Dataset, y = Accuracy, fill=Dataset)) +
    geom_bar(stat = "identity") +
    ylab("Accuracy (%)") +
    xlab(NULL) +
    ggtitle("Comparison of Accuracy: dietaryindex-calculated vs. hand-calculated Dietary Index Values") +
    # add a subtitie
    labs(subtitle = "Exact match by rounding both results to the nearest two decimal places") +
    # increase the title size
    theme(
        plot.title = element_text(size = 20),
        plot.subtitle = element_text(size = 15),
        axis.title = element_text(size = 20),
        axis.text = element_text(size = 15),
        legend.text = element_text(size = 15),
        legend.title = element_text(size = 20),
        axis.text.x = element_blank()
        )

################# AHEI validation ######################

# Initialize a data frame to store results for all dietary index component scores in AHEI
results_AHEI <- data.frame(
  Dataset = character(),
  Accuracy = numeric(),
  stringsAsFactors = FALSE
)

# List of all your datasets and their corresponding column names for AHEI
datasets_AHEI <- list(
    AHEI_ALL = list(data = AHEI_validation_result, cols = c("EXP_AHEI_ALL", "AHEI_ALL")),
    AHEI_NOETOH = list(data = AHEI_validation_result, cols = c("EXP_AHEI_NOETOH", "AHEI_NOETOH")),
    AHEI_VEG = list(data = AHEI_validation_result, cols = c("EXP_AHEI_VEG", "AHEI_VEG")),
    AHEI_FRT = list(data = AHEI_validation_result, cols = c("EXP_AHEI_FRT", "AHEI_FRT")),
    AHEI_WGRAIN = list(data = AHEI_validation_result, cols = c("EXP_AHEI_WGRAIN", "AHEI_WGRAIN")),
    AHEI_NUTSLEG = list(data = AHEI_validation_result, cols = c("EXP_AHEI_NUTSLEG", "AHEI_NUTSLEG")),
    AHEI_N3FAT = list(data = AHEI_validation_result, cols = c("EXP_AHEI_N3FAT", "AHEI_N3FAT")),
    AHEI_PUFA = list(data = AHEI_validation_result, cols = c("EXP_AHEI_PUFA", "AHEI_PUFA")),
    AHEI_SSB_FRTJ = list(data = AHEI_validation_result, cols = c("EXP_AHEI_SSB_FRTJ", "AHEI_SSB_FRTJ")),
    AHEI_REDPROC_MEAT = list(data = AHEI_validation_result, cols = c("EXP_AHEI_REDPROC_MEAT", "AHEI_REDPROC_MEAT")),
    AHEI_TRANS = list(data = AHEI_validation_result, cols = c("EXP_AHEI_TRANS", "AHEI_TRANS")),
    AHEI_SODIUM = list(data = AHEI_validation_result, cols = c("EXP_AHEI_SODIUM", "AHEI_SODIUM")),
    AHEI_ALCOHOL = list(data = AHEI_validation_result, cols = c("EXP_AHEI_ALCOHOL", "AHEI_ALCOHOL"))
)

# Compute accuracy for each dataset in AHEI
for (name in names(datasets_AHEI)) {
    dataset <- datasets_AHEI[[name]]
    accuracy <- get_accuracy(dataset$data[[dataset$cols[1]]], dataset$data[[dataset$cols[2]]])
    results_AHEI <- rbind(results_AHEI, data.frame(Dataset = name, Accuracy = accuracy))
}

# Print results
print(results_AHEI)

# Plot results
ggplot(results_AHEI, aes(x = Dataset, y = Accuracy, fill=Dataset)) +
    geom_bar(stat = "identity") +
    ylab("Accuracy (%)") +
    xlab(NULL) +
    ggtitle("Comparison of Accuracy: dietaryindex-calculated vs. hand-calculated Dietary Index Values") +
    # add a subtitie
    labs(subtitle = "Exact match by rounding both results to the nearest two decimal places") +
    # increase the title size
    theme(
        plot.title = element_text(size = 20),
        plot.subtitle = element_text(size = 15),
        axis.title = element_text(size = 20),
        axis.text = element_text(size = 15),
        legend.text = element_text(size = 15),
        legend.title = element_text(size = 20),
        axis.text.x = element_blank()
        )

################# AHEIP validation ######################

# Initialize a data frame to store results for all dietary index component scores in AHEIP
results_AHEIP <- data.frame(
  Dataset = character(),
  Accuracy = numeric(),
  stringsAsFactors = FALSE
)

# List of all your datasets and their corresponding column names for AHEIP
datasets_AHEIP <- list(
    AHEIP_ALL = list(data = AHEIP_validation_result, cols = c("EXP_AHEIP_ALL", "AHEIP_ALL")),
    AHEIP_VEGETABLE = list(data = AHEIP_validation_result, cols = c("EXP_AHEIP_VEGETABLE", "AHEIP_VEG")),
    AHEIP_WHOLE_FRUIT = list(data = AHEIP_validation_result, cols = c("EXP_AHEIP_WHOLE_FRUIT", "AHEIP_FRT")),
    AHEIP_WHITE_MEAT_RED_MEAT = list(data = AHEIP_validation_result, cols = c("EXP_AHEIP_WHITE_MEAT_RED_MEAT", "AHEIP_WHITEREAD")),
    AHEIP_FIBER = list(data = AHEIP_validation_result, cols = c("EXP_AHEIP_FIBER", "AHEIP_FIBER")),
    AHEIP_TRANS_FAT = list(data = AHEIP_validation_result, cols = c("EXP_AHEIP_TRANS_FAT", "AHEIP_TRANS")),
    AHEIP_POLY_FAT_SAT_FAT = list(data = AHEIP_validation_result, cols = c("EXP_AHEIP_POLY_FAT_SAT_FAT", "AHEIP_POLYSAT")),
    AHEIP_CALCIUM = list(data = AHEIP_validation_result, cols = c("EXP_AHEIP_CALCIUM", "AHEIP_CALCIUM")),
    AHEIP_FOLATE = list(data = AHEIP_validation_result, cols = c("EXP_AHEIP_FOLATE", "AHEIP_FOLATE")),
    AHEIP_IRON = list(data = AHEIP_validation_result, cols = c("EXP_AHEIP_IRON", "AHEIP_IRON"))
)

# Compute accuracy for each dataset in AHEIP
for (name in names(datasets_AHEIP)) {
    dataset <- datasets_AHEIP[[name]]
    accuracy <- get_accuracy(dataset$data[[dataset$cols[1]]], dataset$data[[dataset$cols[2]]])
    results_AHEIP <- rbind(results_AHEIP, data.frame(Dataset = name, Accuracy = accuracy))
}

# Print results
print(results_AHEIP)

# Plot results
ggplot(results_AHEIP, aes(x = Dataset, y = Accuracy, fill=Dataset)) +
    geom_bar(stat = "identity") +
    ylab("Accuracy (%)") +
    xlab(NULL) +
    ggtitle("Comparison of Accuracy: dietaryindex-calculated vs. hand-calculated Dietary Index Values") +
    # add a subtitie
    labs(subtitle = "Exact match by rounding both results to the nearest two decimal places") +
    # increase the title size
    theme(
        plot.title = element_text(size = 20),
        plot.subtitle = element_text(size = 15),
        axis.title = element_text(size = 20),
        axis.text = element_text(size = 15),
        legend.text = element_text(size = 15),
        legend.title = element_text(size = 20),
        axis.text.x = element_blank()
        )

################# DASH validation ######################

# Initialize a data frame to store results for all dietary index component scores in DASH
results_DASH <- data.frame(
  Dataset = character(),
  Accuracy = numeric(),
  stringsAsFactors = FALSE
)

# List of all your datasets and their corresponding column names for DASH
datasets_DASH <- list(
    DASH_ALL = list(data = DASH_validation_result, cols = c("EXP_DASH_ALL", "DASH_ALL")),
    DASH_FRUIT = list(data = DASH_validation_result, cols = c("EXP_DASH_FRUIT", "DASH_FRT")),
    DASH_VEGETABLE = list(data = DASH_validation_result, cols = c("EXP_DASH_VEGETABLE", "DASH_VEG")),
    DASH_NUT_LEGUME = list(data = DASH_validation_result, cols = c("EXP_DASH_NUT_LEGUME", "DASH_NUTSLEG")),
    DASH_WHOLE_GRAIN = list(data = DASH_validation_result, cols = c("EXP_DASH_WHOLE_GRAIN", "DASH_WGRAIN")),
    DASH_LOW_FAT_DAIRY = list(data = DASH_validation_result, cols = c("EXP_DASH_LOW_FAT_DAIRY", "DASH_LOWF_DAIRY")),
    DASH_SODIUM = list(data = DASH_validation_result, cols = c("EXP_DASH_SODIUM", "DASH_SODIUM")),
    DASH_RED_PROCESSED_MEAT = list(data = DASH_validation_result, cols = c("EXP_DASH_RED_PROCESSED_MEAT", "DASH_REDPROC_MEAT")),
    DASH_SSB = list(data = DASH_validation_result, cols = c("EXP_DASH_SSB", "DASH_SSB_FRTJ"))
)

# Compute accuracy for each dataset in DASH
for (name in names(datasets_DASH)) {
    dataset <- datasets_DASH[[name]]
    accuracy <- get_accuracy(dataset$data[[dataset$cols[1]]], dataset$data[[dataset$cols[2]]])
    results_DASH <- rbind(results_DASH, data.frame(Dataset = name, Accuracy = accuracy))
}

# Print results
print(results_DASH)

# Plot results
ggplot(results_DASH, aes(x = Dataset, y = Accuracy, fill=Dataset)) +
    geom_bar(stat = "identity") +
    ylab("Accuracy (%)") +
    xlab(NULL) +
    ggtitle("Comparison of Accuracy: dietaryindex-calculated vs. hand-calculated Dietary Index Values") +
    # add a subtitie
    labs(subtitle = "Exact match by rounding both results to the nearest two decimal places") +
    # increase the title size
    theme(
        plot.title = element_text(size = 20),
        plot.subtitle = element_text(size = 15),
        axis.title = element_text(size = 20),
        axis.text = element_text(size = 15),
        legend.text = element_text(size = 15),
        legend.title = element_text(size = 20),
        axis.text.x = element_blank()
        )

################# DASHI validation ######################

# Initialize a data frame to store results for all dietary index component scores in DASHI
results_DASHI <- data.frame(
  Dataset = character(),
  Accuracy = numeric(),
  stringsAsFactors = FALSE
)

colnames(DASHI_validation_result)

# List of all your datasets and their corresponding column names for DASHI
datasets_DASHI <- list(
    DASHI_ALL = list(data = DASHI_validation_result, cols = c("EXP_DASHI_ALL", "DASHI_ALL")),
    DASHI_TOTAL_FAT = list(data = DASHI_validation_result, cols = c("EXP_DASHI_TOTAL_FAT", "DASHI_TOTAL_FAT")),
    DASHI_SAT_FAT = list(data = DASHI_validation_result, cols = c("EXP_DASHI_SAT_FAT", "DASHI_SAT_FAT")),
    DASHI_PROTEIN = list(data = DASHI_validation_result, cols = c("EXP_DASHI_PROTEIN", "DASHI_PROTEIN")),
    DASHI_CHOLESTEROL = list(data = DASHI_validation_result, cols = c("EXP_DASHI_CHOLESTEROL", "DASHI_CHOLESTEROL")),
    DASHI_FIBER = list(data = DASHI_validation_result, cols = c("EXP_DASHI_FIBER", "DASHI_FIBER")),
    DASHI_POTASSIUM = list(data = DASHI_validation_result, cols = c("EXP_DASHI_POTASSIUM", "DASHI_POTASSIUM")),
    DASHI_MAGNESIUM = list(data = DASHI_validation_result, cols = c("EXP_DASHI_MAGNESIUM", "DASHI_MAGNESIUM")),
    DASHI_CALCIUM = list(data = DASHI_validation_result, cols = c("EXP_DASHI_CALCIUM", "DASHI_CALCIUM")),
    DASHI_SODIUM = list(data = DASHI_validation_result, cols = c("EXP_DASHI_SODIUM", "DASHI_SODIUM"))
)


# Compute accuracy for each dataset in DASHI
for (name in names(datasets_DASHI)) {
    dataset <- datasets_DASHI[[name]]
    accuracy <- get_accuracy(dataset$data[[dataset$cols[1]]], dataset$data[[dataset$cols[2]]])
    results_DASHI <- rbind(results_DASHI, data.frame(Dataset = name, Accuracy = accuracy))
}

# Print results
print(results_DASHI)

# Plot results
ggplot(results_DASHI, aes(x = Dataset, y = Accuracy, fill=Dataset)) +
    geom_bar(stat = "identity") +
    ylab("Accuracy (%)") +
    xlab(NULL) +
    ggtitle("Comparison of Accuracy: dietaryindex-calculated vs. hand-calculated Dietary Index Values") +
    # add a subtitie
    labs(subtitle = "Exact match by rounding both results to the nearest two decimal places") +
    # increase the title size
    theme(
        plot.title = element_text(size = 20),
        plot.subtitle = element_text(size = 15),
        axis.title = element_text(size = 20),
        axis.text = element_text(size = 15),
        legend.text = element_text(size = 15),
        legend.title = element_text(size = 20),
        axis.text.x = element_blank()
        )

################# DII validation ######################

# Initialize a data frame to store results for all dietary index component scores in DII
results_DII <- data.frame(
  Dataset = character(),
  Accuracy = numeric(),
  stringsAsFactors = FALSE
)

# List of all your datasets and their corresponding column names for DII
datasets_DII <- list(
    DII_ALL = list(data = DII_validation_result, cols = c("EXP_DII_ALL", "DII_ALL")),
    ALCOHOL_DII = list(data = DII_validation_result, cols = c("EXP_ALCOHOL_DII", "ALCOHOL_DII")),
    VITB12_DII = list(data = DII_validation_result, cols = c("EXP_VITB12_DII", "VITB12_DII")),
    VITB6_DII = list(data = DII_validation_result, cols = c("EXP_VITB6_DII", "VITB6_DII")),
    BCAROTENE_DII = list(data = DII_validation_result, cols = c("EXP_BCAROTENE_DII", "BCAROTENE_DII")),
    CAFFEINE_DII = list(data = DII_validation_result, cols = c("EXP_CAFFEINE_DII", "CAFFEINE_DII")),
    CARB_DII = list(data = DII_validation_result, cols = c("EXP_CARB_DII", "CARB_DII")),
    CHOLESTEROL_DII = list(data = DII_validation_result, cols = c("EXP_CHOLES_DII", "CHOLES_DII")),
    TOTALFAT_DII = list(data = DII_validation_result, cols = c("EXP_TOTALFAT_DII", "TOTALFAT_DII")),
    FIBER_DII = list(data = DII_validation_result, cols = c("EXP_FIBER_DII", "FIBER_DII")),
    FOLICACID_DII = list(data = DII_validation_result, cols = c("EXP_FOLICACID_DII", "FOLICACID_DII")),
    GARLIC_DII = list(data = DII_validation_result, cols = c("EXP_GARLIC_DII", "GARLIC_DII")),
    GINGER_DII = list(data = DII_validation_result, cols = c("EXP_GINGER_DII", "GINGER_DII")),
    IRON_DII = list(data = DII_validation_result, cols = c("EXP_IRON_DII", "IRON_DII")),
    MAGNESIUM_DII = list(data = DII_validation_result, cols = c("EXP_MG_DII", "MG_DII")),
    MUFA_DII = list(data = DII_validation_result, cols = c("EXP_MUFA_DII", "MUFA_DII")),
    NIACIN_DII = list(data = DII_validation_result, cols = c("EXP_NIACIN_DII", "NIACIN_DII")),
    N3FAT_DII = list(data = DII_validation_result, cols = c("EXP_N3FAT_DII", "N3FAT_DII")),
    N6FAT_DII = list(data = DII_validation_result, cols = c("EXP_N6FAT_DII", "N6FAT_DII")),
    ONION_DII = list(data = DII_validation_result, cols = c("EXP_ONION_DII", "ONION_DII")),
    PROTEIN_DII = list(data = DII_validation_result, cols = c("EXP_PROTEIN_DII", "PROTEIN_DII")),
    PUFA_DII = list(data = DII_validation_result, cols = c("EXP_PUFA_DII", "PUFA_DII")),
    RIBOFLAVIN_DII = list(data = DII_validation_result, cols = c("EXP_RIBOFLAVIN_DII", "RIBOFLAVIN_DII")),
    SAFFRON_DII = list(data = DII_validation_result, cols = c("EXP_SAFFRON_DII", "SAFFRON_DII")),
    SATFAT_DII = list(data = DII_validation_result, cols = c("EXP_SATFAT_DII", "SATFAT_DII")),
    SELENIUM_DII = list(data = DII_validation_result, cols = c("EXP_SE_DII", "SE_DII")),
    THIAMIN_DII = list(data = DII_validation_result, cols = c("EXP_THIAMIN_DII", "THIAMIN_DII")),
    TRANSFAT_DII = list(data = DII_validation_result, cols = c("EXP_TRANSFAT_DII", "TRANSFAT_DII")),
    TURMERIC_DII = list(data = DII_validation_result, cols = c("EXP_TURMERIC_DII", "TURMERIC_DII")),
    VITAMINA_DII = list(data = DII_validation_result, cols = c("EXP_VITA_DII", "VITA_DII")),
    VITAMINC_DII = list(data = DII_validation_result, cols = c("EXP_VITC_DII", "VITC_DII")),
    VITAMIND_DII = list(data = DII_validation_result, cols = c("EXP_VITD_DII", "VITD_DII")),
    VITAMINE_DII = list(data = DII_validation_result, cols = c("EXP_VITE_DII", "VITE_DII")),
    ZINC_DII = list(data = DII_validation_result, cols = c("EXP_ZN_DII", "ZN_DII")),
    TEA_DII = list(data = DII_validation_result, cols = c("EXP_TEA_DII", "TEA_DII")),
    FLA3OL_DII = list(data = DII_validation_result, cols = c("EXP_FLA3OL_DII", "FLA3OL_DII")),
    FLAVONES_DII = list(data = DII_validation_result, cols = c("EXP_FLAVONES_DII", "FLAVONES_DII")),
    FLAVONOLS_DII = list(data = DII_validation_result, cols = c("EXP_FLAVONOLS_DII", "FLAVONOLS_DII")),
    FLAVONONES_DII = list(data = DII_validation_result, cols = c("EXP_FLAVONONES_DII", "FLAVONONES_DII")),
    ANTHOCYANIDINS_DII = list(data = DII_validation_result, cols = c("EXP_ANTHOC_DII", "ANTHOC_DII")),
    ISOFLAVONES_DII = list(data = DII_validation_result, cols = c("EXP_ISOFLAVONES_DII", "ISOFLAVONES_DII")),
    PEPPER_DII = list(data = DII_validation_result, cols = c("EXP_PEPPER_DII", "PEPPER_DII")),
    THYME_DII = list(data = DII_validation_result, cols = c("EXP_THYME_DII", "THYME_DII")),
    ROSEMARY_DII = list(data = DII_validation_result, cols = c("EXP_ROSEMARY_DII", "ROSEMARY_DII"))
)

# Compute accuracy for each dataset in DII
for (name in names(datasets_DII)) {
    dataset <- datasets_DII[[name]]
    accuracy <- get_accuracy(dataset$data[[dataset$cols[1]]], dataset$data[[dataset$cols[2]]])
    results_DII <- rbind(results_DII, data.frame(Dataset = name, Accuracy = accuracy))
}

# Print results
print(results_DII)

# Plot results
ggplot(results_DII, aes(x = Dataset, y = Accuracy, fill=Dataset)) +
    geom_bar(stat = "identity") +
    ylab("Accuracy (%)") +
    xlab(NULL) +
    ggtitle("Comparison of Accuracy: dietaryindex-calculated vs. hand-calculated Dietary Index Values") +
    # add a subtitie
    labs(subtitle = "Exact match by rounding both results to the nearest two decimal places") +
    # increase the title size
    theme(
        plot.title = element_text(size = 20),
        plot.subtitle = element_text(size = 15),
        axis.title = element_text(size = 20),
        axis.text = element_text(size = 15),
        legend.text = element_text(size = 15),
        legend.title = element_text(size = 20),
        axis.text.x = element_blank()
        )

################### HEI2015 validation ######################

# Initialize a data frame to store results for all dietary index component scores in HEI2015
results_HEI2015 <- data.frame(
  Dataset = character(),
  Accuracy = numeric(),
  stringsAsFactors = FALSE
)

# List of all your datasets and their corresponding column names for HEI2015
datasets_HEI2015 <- list(
    HEI2015_ALL = list(data = HEI2015_validation_result, cols = c("EXP_HEI2015_ALL", "HEI2015_ALL")),
    HEI2015_TOTALFRT = list(data = HEI2015_validation_result, cols = c("EXP_HEI2015_TOTALFRT", "HEI2015_TOTALFRT")),
    HEI2015_FRT = list(data = HEI2015_validation_result, cols = c("EXP_HEI2015_FRT", "HEI2015_FRT")),
    HEI2015_VEG = list(data = HEI2015_validation_result, cols = c("EXP_HEI2015_VEG", "HEI2015_VEG")),
    HEI2015_GREENNBEAN = list(data = HEI2015_validation_result, cols = c("EXP_HEI2015_GREENNBEAN", "HEI2015_GREENNBEAN")),
    HEI2015_TOTALPRO = list(data = HEI2015_validation_result, cols = c("EXP_HEI2015_TOTALPRO", "HEI2015_TOTALPRO")),
    HEI2015_SEAPLANTPRO = list(data = HEI2015_validation_result, cols = c("EXP_HEI2015_SEAPLANTPRO", "HEI2015_SEAPLANTPRO")),
    HEI2015_WHOLEGRAIN = list(data = HEI2015_validation_result, cols = c("EXP_HEI2015_WHOLEGRAIN", "HEI2015_WHOLEGRAIN")),
    HEI2015_DAIRY = list(data = HEI2015_validation_result, cols = c("EXP_HEI2015_DAIRY", "HEI2015_DAIRY")),
    HEI2015_FATTYACID = list(data = HEI2015_validation_result, cols = c("EXP_HEI2015_FATTYACID", "HEI2015_FATTYACID")),
    HEI2015_REFINEDGRAIN = list(data = HEI2015_validation_result, cols = c("EXP_HEI2015_REFINEDGRAIN", "HEI2015_REFINEDGRAIN")),
    HEI2015_SODIUM = list(data = HEI2015_validation_result, cols = c("EXP_HEI2015_SODIUM", "HEI2015_SODIUM")),
    HEI2015_ADDEDSUGAR = list(data = HEI2015_validation_result, cols = c("EXP_HEI2015_ADDEDSUGAR", "HEI2015_ADDEDSUGAR")),
    HEI2015_SATFAT = list(data = HEI2015_validation_result, cols = c("EXP_HEI2015_SATFAT", "HEI2015_SATFAT"))
)

# Compute accuracy for each dataset in HEI2015
for (name in names(datasets_HEI2015)) {
    dataset <- datasets_HEI2015[[name]]
    accuracy <- get_accuracy(dataset$data[[dataset$cols[1]]], dataset$data[[dataset$cols[2]]])
    results_HEI2015 <- rbind(results_HEI2015, data.frame(Dataset = name, Accuracy = accuracy))
}

# Print results
print(results_HEI2015)

# Plot results
ggplot(results_HEI2015, aes(x = Dataset, y = Accuracy, fill=Dataset)) +
    geom_bar(stat = "identity") +
    ylab("Accuracy (%)") +
    xlab(NULL) +
    ggtitle("Comparison of Accuracy: dietaryindex-calculated vs. hand-calculated Dietary Index Values") +
    # add a subtitie
    labs(subtitle = "Exact match by rounding both results to the nearest two decimal places") +
    # increase the title size
    theme(
        plot.title = element_text(size = 15),
        plot.subtitle = element_text(size = 10),
        axis.title = element_text(size = 15),
        axis.text = element_text(size = 10),
        legend.text = element_text(size = 10),
        legend.title = element_text(size = 15),
        axis.text.x = element_blank()
        )

################### HEI2020 validation ######################

# Initialize a data frame to store results for all HEI2020 component scores
results_HEI2020 <- data.frame(
  Dataset = character(),
  Accuracy = numeric(),
  stringsAsFactors = FALSE
)

# List of all the datasets and their corresponding column names for HEI2020
datasets_HEI2020 <- list(
  HEI2020_ALL = list(data = HEI2020_validation_result, cols = c("EXP_HEI2020_ALL", "HEI2020_ALL")),
  HEI2020_TOTALFRT = list(data = HEI2020_validation_result, cols = c("EXP_HEI2020_TOTALFRT", "HEI2020_TOTALFRT")),
  HEI2020_FRT = list(data = HEI2020_validation_result, cols = c("EXP_HEI2020_FRT", "HEI2020_FRT")),
  HEI2020_VEG = list(data = HEI2020_validation_result, cols = c("EXP_HEI2020_VEG", "HEI2020_VEG")),
  HEI2020_GREENNBEAN = list(data = HEI2020_validation_result, cols = c("EXP_HEI2020_GREENNBEAN", "HEI2020_GREENNBEAN")),
  HEI2020_TOTALPRO = list(data = HEI2020_validation_result, cols = c("EXP_HEI2020_TOTALPRO", "HEI2020_TOTALPRO")),
  HEI2020_SEAPLANTPRO = list(data = HEI2020_validation_result, cols = c("EXP_HEI2020_SEAPLANTPRO", "HEI2020_SEAPLANTPRO")),
  HEI2020_WHOLEGRAIN = list(data = HEI2020_validation_result, cols = c("EXP_HEI2020_WHOLEGRAIN", "HEI2020_WHOLEGRAIN")),
  HEI2020_DAIRY = list(data = HEI2020_validation_result, cols = c("EXP_HEI2020_DAIRY", "HEI2020_DAIRY")),
  HEI2020_FATTYACID = list(data = HEI2020_validation_result, cols = c("EXP_HEI2020_FATTYACID", "HEI2020_FATTYACID")),
  HEI2020_REFINEDGRAIN = list(data = HEI2020_validation_result, cols = c("EXP_HEI2020_REFINEDGRAIN", "HEI2020_REFINEDGRAIN")),
  HEI2020_SODIUM = list(data = HEI2020_validation_result, cols = c("EXP_HEI2020_SODIUM", "HEI2020_SODIUM")),
  HEI2020_ADDEDSUGAR = list(data = HEI2020_validation_result, cols = c("EXP_HEI2020_ADDEDSUGAR", "HEI2020_ADDEDSUGAR")),
  HEI2020_SATFAT = list(data = HEI2020_validation_result, cols = c("EXP_HEI2020_SATFAT", "HEI2020_SATFAT"))
)

# Compute accuracy for each dataset in HEI2020
for (name in names(datasets_HEI2020)) {
  dataset <- datasets_HEI2020[[name]]
  accuracy <- get_accuracy(dataset$data[[dataset$cols[1]]], dataset$data[[dataset$cols[2]]])
  results_HEI2020 <- rbind(results_HEI2020, data.frame(Dataset = name, Accuracy = accuracy))
}

# Print results
print(results_HEI2020)

# Plot results
ggplot(results_HEI2020, aes(x = Dataset, y = Accuracy, fill = Dataset)) +
  geom_bar(stat = "identity") +
  ylab("Accuracy (%)") +
  xlab(NULL) +
  ggtitle("Comparison of Accuracy: dietaryindex-calculated vs. hand-calculated Dietary Index Values") +
  # add a subtitie
  labs(subtitle = "Exact match by rounding both results to the nearest two decimal places") +
  # increase the title size
  theme(
    plot.title = element_text(size = 20),
    plot.subtitle = element_text(size = 15),
    axis.title = element_text(size = 20),
    axis.text = element_text(size = 15),
    legend.text = element_text(size = 15),
    legend.title = element_text(size = 20),
    axis.text.x = element_blank()
  )

################### MED validation ######################

# Initialize a data frame to store results for all MED component scores
results_MED <- data.frame(
  Dataset = character(),
  Accuracy = numeric(),
  stringsAsFactors = FALSE
)

# List of all the datasets and their corresponding column names for MED
datasets_MED <- list(
  MED_ALL = list(data = MED_validation_result, cols = c("EXP_MED_ALL", "MED_ALL")),
  MED_FRUIT = list(data = MED_validation_result, cols = c("EXP_MED_FRUIT", "MED_FRT")),
  MED_VEGETABLE = list(data = MED_validation_result, cols = c("EXP_MED_VEGETABLE", "MED_VEG")),
  MED_WHOLE_GRAIN = list(data = MED_validation_result, cols = c("EXP_MED_WHOLE_GRAIN", "MED_WGRAIN")),
  MED_LEGUME = list(data = MED_validation_result, cols = c("EXP_MED_LEGUME", "MED_LEGUMES")),
  MED_NUT = list(data = MED_validation_result, cols = c("EXP_MED_NUT", "MED_NUTS")),
  MED_FISH = list(data = MED_validation_result, cols = c("EXP_MED_FISH", "MED_FISH")),
  MED_RED_PROCESSED_MEAT = list(data = MED_validation_result, cols = c("EXP_MED_RED_PROCESSED_MEAT", "MED_REDPROC_MEAT")),
  MED_MONOFAT_SATFAT = list(data = MED_validation_result, cols = c("EXP_MED_MONOFAT_SATFAT", "MED_MONSATFAT")),
  MED_ALCOHOL = list(data = MED_validation_result, cols = c("EXP_MED_ALCOHOL", "MED_ALCOHOL"))
)

# Compute accuracy for each dataset in MED
for (name in names(datasets_MED)) {
  dataset <- datasets_MED[[name]]
  accuracy <- get_accuracy(dataset$data[[dataset$cols[1]]], dataset$data[[dataset$cols[2]]])
  results_MED <- rbind(results_MED, data.frame(Dataset = name, Accuracy = accuracy))
}

# Print results
print(results_MED)

# Plot results
ggplot(results_MED, aes(x = Dataset, y = Accuracy, fill = Dataset)) +
  geom_bar(stat = "identity") +
  ylab("Accuracy (%)") +
  xlab(NULL) +
  ggtitle("Comparison of Accuracy: dietaryindex-calculated vs. hand-calculated Dietary Index Values") +
  # add a subtitie
  labs(subtitle = "Exact match by rounding both results to the nearest two decimal places") +
  # increase the title size
  theme(
    plot.title = element_text(size = 20),
    plot.subtitle = element_text(size = 15),
    axis.title = element_text(size = 20),
    axis.text = element_text(size = 15),
    legend.text = element_text(size = 15),
    legend.title = element_text(size = 20),
    axis.text.x = element_blank()
  )

################### MEDI validation ######################

# Initialize a data frame to store results for all MEDI component scores
results_MEDI <- data.frame(
  Dataset = character(),
  Accuracy = numeric(),
  stringsAsFactors = FALSE
)

# List of all the datasets and their corresponding column names for MEDI
datasets_MEDI <- list(
  MEDI_ALL = list(data = MEDI_validation_result, cols = c("EXP_MEDI_ALL", "MEDI_ALL")),
  MEDI_OLIVE_OIL = list(data = MEDI_validation_result, cols = c("EXP_MEDI_OLIVE_OIL", "MEDI_OLIVE_OIL")),
  MEDI_VEGETABLE = list(data = MEDI_validation_result, cols = c("EXP_MEDI_VEGETABLE", "MEDI_VEG")),
  MEDI_FRUIT = list(data = MEDI_validation_result, cols = c("EXP_MEDI_FRUIT", "MEDI_FRT")),
  MEDI_LEGUME = list(data = MEDI_validation_result, cols = c("EXP_MEDI_LEGUME", "MEDI_LEGUMES")),
  MEDI_NUT = list(data = MEDI_validation_result, cols = c("EXP_MEDI_NUT", "MEDI_NUTS")),
  MEDI_FISH = list(data = MEDI_validation_result, cols = c("EXP_MEDI_FISH", "MEDI_FISH")),
  MEDI_ALCOHOL = list(data = MEDI_validation_result, cols = c("EXP_MEDI_ALCOHOL", "MEDI_ALCOHOL")),
  MEDI_SSB = list(data = MEDI_validation_result, cols = c("EXP_MEDI_SSB", "MEDI_SSB")),
  MEDI_SWEETS = list(data = MEDI_validation_result, cols = c("EXP_MEDI_SWEETS", "MEDI_SWEETS")),
  MEDI_DISCRET_FAT = list(data = MEDI_validation_result, cols = c("EXP_MEDI_DISCRET_FAT", "MEDI_DISCRET_FAT")),
  MEDI_RED_MEAT = list(data = MEDI_validation_result, cols = c("EXP_MEDI_RED_MEAT", "MEDI_REDPROC_MEAT"))
)

# Compute accuracy for each dataset in MEDI
for (name in names(datasets_MEDI)) {
  dataset <- datasets_MEDI[[name]]
  accuracy <- get_accuracy(dataset$data[[dataset$cols[1]]], dataset$data[[dataset$cols[2]]])
  results_MEDI <- rbind(results_MEDI, data.frame(Dataset = name, Accuracy = accuracy))
}

# Print results
print(results_MEDI)

# Plot results
ggplot(results_MEDI, aes(x = Dataset, y = Accuracy, fill = Dataset)) +
  geom_bar(stat = "identity") +
  ylab("Accuracy (%)") +
  xlab(NULL) +
  ggtitle("Comparison of Accuracy: dietaryindex-calculated vs. hand-calculated Dietary Index Values") +
  # add a subtitie
  labs(subtitle = "Exact match by rounding both results to the nearest two decimal places") +
  # increase the title size
  theme(
    plot.title = element_text(size = 20),
    plot.subtitle = element_text(size = 15),
    axis.title = element_text(size = 20),
    axis.text = element_text(size = 15),
    legend.text = element_text(size = 15),
    legend.title = element_text(size = 20),
    axis.text.x = element_blank()
  )

################### MEDI_V2 validation ######################

# Initialize a data frame to store results for all MEDI_V2 component scores
results_MEDI_V2 <- data.frame(
  Dataset = character(),
  Accuracy = numeric(),
  stringsAsFactors = FALSE
)

# List of all the datasets and their corresponding column names for MEDI_V2
datasets_MEDI_V2 <- list(
  MEDI_V2_ALL = list(data = MEDI_V2_validation_result, cols = c("EXP_MEDI_ALL", "MEDI_V2_ALL")),
  MEDI_V2_OLIVE_OIL = list(data = MEDI_V2_validation_result, cols = c("EXP_MEDI_OLIVE_OIL", "MEDI_OLIVE_OIL")),
  MEDI_V2_VEGETABLE = list(data = MEDI_V2_validation_result, cols = c("EXP_MEDI_VEGETABLE", "MEDI_VEG")),
  MEDI_V2_FRUIT = list(data = MEDI_V2_validation_result, cols = c("EXP_MEDI_FRUIT", "MEDI_FRT")),
  MEDI_V2_LEGUME = list(data = MEDI_V2_validation_result, cols = c("EXP_MEDI_LEGUME", "MEDI_LEGUMES")),
  MEDI_V2_NUT = list(data = MEDI_V2_validation_result, cols = c("EXP_MEDI_NUT", "MEDI_NUTS")),
  MEDI_V2_FISH = list(data = MEDI_V2_validation_result, cols = c("EXP_MEDI_FISH", "MEDI_FISH")),
  MEDI_V2_ALCOHOL = list(data = MEDI_V2_validation_result, cols = c("EXP_MEDI_ALCOHOL", "MEDI_ALCOHOL")),
  MEDI_V2_SSB = list(data = MEDI_V2_validation_result, cols = c("EXP_MEDI_SSB", "MEDI_SSB")),
  MEDI_V2_SWEETS = list(data = MEDI_V2_validation_result, cols = c("EXP_MEDI_SWEETS", "MEDI_SWEETS")),
  MEDI_V2_DISCRET_FAT = list(data = MEDI_V2_validation_result, cols = c("EXP_MEDI_DISCRET_FAT", "MEDI_DISCRET_FAT")),
  MEDI_V2_RED_MEAT = list(data = MEDI_V2_validation_result, cols = c("EXP_MEDI_RED_MEAT", "MEDI_REDPROC_MEAT"))
)

# Compute accuracy for each dataset in MEDI_V2
for (name in names(datasets_MEDI_V2)) {
  dataset <- datasets_MEDI_V2[[name]]
  accuracy <- get_accuracy(dataset$data[[dataset$cols[1]]], dataset$data[[dataset$cols[2]]])
  results_MEDI_V2 <- rbind(results_MEDI_V2, data.frame(Dataset = name, Accuracy = accuracy))
}

# Print results
print(results_MEDI_V2)

# Plot results
ggplot(results_MEDI_V2, aes(x = Dataset, y = Accuracy, fill = Dataset)) +
  geom_bar(stat = "identity") +
  ylab("Accuracy (%)") +
  xlab(NULL) +
  ggtitle("Comparison of Accuracy: dietaryindex-calculated vs. hand-calculated Dietary Index Values") +
  # add a subtitie
  labs(subtitle = "Exact match by rounding both results to the nearest two decimal places") +
  # increase the title size
  theme(
    plot.title = element_text(size = 20),
    plot.subtitle = element_text(size = 15),
    axis.title = element_text(size = 20),
    axis.text = element_text(size = 15),
    legend.text = element_text(size = 15),
    legend.title = element_text(size = 20),
    axis.text.x = element_blank()
  )


################### PHDI validation ######################

# Initialize a data frame to store results for all PHDI component scores
results_PHDI <- data.frame(
  Dataset = character(),
  Accuracy = numeric(),
  stringsAsFactors = FALSE
)

# List of all the datasets and their corresponding column names for PHDI
datasets_PHDI <- list(
  PHDI_ALL = list(data = PHDI_validation_result, cols = c("EXP_PHDI_ALL", "PHDI_ALL")),
  PHDI_WGRAIN = list(data = PHDI_validation_result, cols = c("EXP_PHDI_WGRAIN", "PHDI_WGRAIN")),
  PHDI_STARCHY_VEG = list(data = PHDI_validation_result, cols = c("EXP_PHDI_STARCHY_VEG", "PHDI_STARCHY_VEG")),
  PHDI_VEG = list(data = PHDI_validation_result, cols = c("EXP_PHDI_VEG", "PHDI_VEG")),
  PHDI_FRT = list(data = PHDI_validation_result, cols = c("EXP_PHDI_FRT", "PHDI_FRT")),
  PHDI_DAIRY = list(data = PHDI_validation_result, cols = c("EXP_PHDI_DAIRY", "PHDI_DAIRY")),
  PHDI_REDPROC_MEAT = list(data = PHDI_validation_result, cols = c("EXP_PHDI_REDPROC_MEAT", "PHDI_REDPROC_MEAT")),
  PHDI_POULTRY = list(data = PHDI_validation_result, cols = c("EXP_PHDI_POULTRY", "PHDI_POULTRY")),
  PHDI_EGG = list(data = PHDI_validation_result, cols = c("EXP_PHDI_EGG", "PHDI_EGG")),
  PHDI_FISH = list(data = PHDI_validation_result, cols = c("EXP_PHDI_FISH", "PHDI_FISH")),
  PHDI_NUTS = list(data = PHDI_validation_result, cols = c("EXP_PHDI_NUTS", "PHDI_NUTS")),
  PHDI_LEGUMES = list(data = PHDI_validation_result, cols = c("EXP_PHDI_LEGUMES", "PHDI_LEGUMES")),
  PHDI_SOY = list(data = PHDI_validation_result, cols = c("EXP_PHDI_SOY", "PHDI_SOY")),
  PHDI_ADDED_FAT_UNSAT = list(data = PHDI_validation_result, cols = c("EXP_PHDI_ADDED_FAT_UNSAT", "PHDI_ADDED_FAT_UNSAT")),
  PHDI_ADDED_FAT_SAT = list(data = PHDI_validation_result, cols = c("EXP_PHDI_ADDED_FAT_SAT", "PHDI_ADDED_FAT_SAT")),
  PHDI_ADDED_SUGAR = list(data = PHDI_validation_result, cols = c("EXP_PHDI_ADDED_SUGAR", "PHDI_ADDED_SUGAR"))
)

# Compute accuracy for each dataset in PHDI
for (name in names(datasets_PHDI)) {
  dataset <- datasets_PHDI[[name]]
  accuracy <- get_accuracy(dataset$data[[dataset$cols[1]]], dataset$data[[dataset$cols[2]]])
  results_PHDI <- rbind(results_PHDI, data.frame(Dataset = name, Accuracy = accuracy))
}

# Print results
print(results_PHDI)

# Plot results
ggplot(results_PHDI, aes(x = Dataset, y = Accuracy, fill = Dataset)) +
  geom_bar(stat = "identity") +
  ylab("Accuracy (%)") +
  xlab(NULL) +
  ggtitle("Comparison of Accuracy: dietaryindex-calculated vs. hand-calculated Dietary Index Values") +
  # add a subtitie
  labs(subtitle = "Exact match by rounding both results to the nearest two decimal places") +
  # increase the title size
  theme(
    plot.title = element_text(size = 20),
    plot.subtitle = element_text(size = 15),
    axis.title = element_text(size = 20),
    axis.text = element_text(size = 15),
    legend.text = element_text(size = 15),
    legend.title = element_text(size = 20),
    axis.text.x = element_blank()
  )


###################### HEI2015 validation in NHANES using dietaryindex-calculated results vs. National Cancer Institute (NCI) SAS results ######################
# Initialize a data frame to store results for all HEI2015_1718 results
results_HEI2015_1718 <- data.frame(
  Dataset = character(),
  Accuracy = numeric(),
  stringsAsFactors = FALSE
)

# Create a list with SAS_HEI2015_1718 and dietaryindex_HEI2015_1718
datasets_HEI2015_1718 <- list(
    HEI2015_ALL = list(data1 = SAS_HEI2015_1718, data2 = dietaryindex_HEI2015_1718, cols = c("HEI2015_TOTAL_SCORE", "HEI2015_ALL")),
    HEI2015_TOTALFRT = list(data1 = SAS_HEI2015_1718, data2 = dietaryindex_HEI2015_1718, cols = c("HEI2015C3_TOTALFRUIT", "HEI2015_TOTALFRT")),
    HEI2015_FRT = list(data1 = SAS_HEI2015_1718, data2 = dietaryindex_HEI2015_1718, cols = c("HEI2015C4_WHOLEFRUIT", "HEI2015_FRT")),
    HEI2015_VEG = list(data1 = SAS_HEI2015_1718, data2 = dietaryindex_HEI2015_1718, cols = c("HEI2015C1_TOTALVEG", "HEI2015_VEG")),
    HEI2015_GREENNBEAN = list(data1 = SAS_HEI2015_1718, data2 = dietaryindex_HEI2015_1718, cols = c("HEI2015C2_GREEN_AND_BEAN", "HEI2015_GREENNBEAN")),
    HEI2015_TOTALPRO = list(data1 = SAS_HEI2015_1718, data2 = dietaryindex_HEI2015_1718, cols = c("HEI2015C7_TOTPROT", "HEI2015_TOTALPRO")),
    HEI2015_SEAPLANTPRO = list(data1 = SAS_HEI2015_1718, data2 = dietaryindex_HEI2015_1718, cols = c("HEI2015C8_SEAPLANT_PROT", "HEI2015_SEAPLANTPRO")),
    HEI2015_WHOLEGRAIN = list(data1 = SAS_HEI2015_1718, data2 = dietaryindex_HEI2015_1718, cols = c("HEI2015C5_WHOLEGRAIN", "HEI2015_WHOLEGRAIN")),
    HEI2015_DAIRY = list(data1 = SAS_HEI2015_1718, data2 = dietaryindex_HEI2015_1718, cols = c("HEI2015C6_TOTALDAIRY", "HEI2015_DAIRY")),
    HEI2015_FATTYACID = list(data1 = SAS_HEI2015_1718, data2 = dietaryindex_HEI2015_1718, cols = c("HEI2015C9_FATTYACID", "HEI2015_FATTYACID")),
    HEI2015_REFINEDGRAIN = list(data1 = SAS_HEI2015_1718, data2 = dietaryindex_HEI2015_1718, cols = c("HEI2015C11_REFINEDGRAIN", "HEI2015_REFINEDGRAIN")),
    HEI2015_SODIUM = list(data1 = SAS_HEI2015_1718, data2 = dietaryindex_HEI2015_1718, cols = c("HEI2015C10_SODIUM", "HEI2015_SODIUM")),
    HEI2015_ADDEDSUGAR = list(data1 = SAS_HEI2015_1718, data2 = dietaryindex_HEI2015_1718, cols = c("HEI2015C13_ADDSUG", "HEI2015_ADDEDSUGAR")),
    HEI2015_SATFAT = list(data1 = SAS_HEI2015_1718, data2 = dietaryindex_HEI2015_1718, cols = c("HEI2015C12_SFAT", "HEI2015_SATFAT"))
    )

# Compute accuracy for each dataset in PHDI
for (name in names(datasets_HEI2015_1718)) {
  dataset <- datasets_HEI2015_1718[[name]]
  data1_values <- as.numeric(dataset$data1[[dataset$cols[1]]])  # Convert to numeric
  data2_values <- as.numeric(dataset$data2[[dataset$cols[2]]])  # Convert to numeric

  accuracy <- get_accuracy(data1_values, data2_values)
  results_HEI2015_1718 <- rbind(results_HEI2015_1718, data.frame(Component = name, Accuracy = accuracy))
}

# Print results
print(results_HEI2015_1718)

# Plot results
ggplot(results_HEI2015_1718, aes(x = Component, y = Accuracy, fill = Component)) +
  geom_bar(stat = "identity") +
  ylab("Accuracy (%)") +
  xlab(NULL) +
  ggtitle("Accuracy of HEI2015 in NHANES: dietaryindex-calculated vs. SAS-calculated Dietary Index Values from NCI") +
  # add a subtitie
  labs(subtitle = "Exact match by rounding both results to the nearest two decimal places") +
  # increase the title size
  theme(
    plot.title = element_text(size = 20),
    plot.subtitle = element_text(size = 15),
    axis.title = element_text(size = 20),
    axis.text = element_text(size = 15),
    legend.text = element_text(size = 15),
    legend.title = element_text(size = 20),
    axis.text.x = element_blank()
  )


###################### HEI2015 validation in ASA24 using dietaryindex-calculated results vs. National Cancer Institute (NCI) SAS results ######################
# Initialize a data frame to store results for all HEI2015_ASA24 results
results_HEI2015_ASA24 <- data.frame(
  Dataset = character(),
  Accuracy = numeric(),
  stringsAsFactors = FALSE
)

colnames(HEI2015_ASA24_NCI_SAS)
colnames(HEI2015_ASA24_dietaryindex)

# r$> colnames(HEI2015_ASA24_NCI_SAS)
#  [1] "UserName"                 "UserID"                   "RecallNo"                 "KCAL"                     "HEI2015C1_TOTALVEG"       "HEI2015C2_GREEN_AND_BEAN"
#  [7] "HEI2015C3_TOTALFRUIT"     "HEI2015C4_WHOLEFRUIT"     "HEI2015C5_WHOLEGRAIN"     "HEI2015C6_TOTALDAIRY"     "HEI2015C7_TOTPROT"        "HEI2015C8_SEAPLANT_PROT" 
# [13] "HEI2015C9_FATTYACID"      "HEI2015C10_SODIUM"        "HEI2015C11_REFINEDGRAIN"  "HEI2015C12_SFAT"          "HEI2015C13_ADDSUG"        "HEI2015_TOTAL_SCORE"     

# r$> colnames(HEI2015_ASA24_dietaryindex)
#  [1] "UserName"             "UserID"               "TOTALKCAL"            "HEI2015_ALL"          "HEI2015_TOTALFRT"     "HEI2015_FRT"          "HEI2015_VEG"          "HEI2015_GREENNBEAN"  
#  [9] "HEI2015_TOTALPRO"     "HEI2015_SEAPLANTPRO"  "HEI2015_WHOLEGRAIN"   "HEI2015_DAIRY"        "HEI2015_FATTYACID"    "HEI2015_REFINEDGRAIN" "HEI2015_SODIUM"       "HEI2015_ADDEDSUGAR"  
# [17] "HEI2015_SATFAT"

# Create a list with HEI2015_ASA24_NCI_SAS and HEI2015_ASA24_dietaryindex
datasets_HEI2015_ASA24<- list(
    HEI2015_ALL = list(data1 = HEI2015_ASA24_NCI_SAS, data2 = HEI2015_ASA24_dietaryindex, cols = c("HEI2015_TOTAL_SCORE", "HEI2015_ALL")),
    HEI2015_TOTALFRT = list(data1 = HEI2015_ASA24_NCI_SAS, data2 = HEI2015_ASA24_dietaryindex, cols = c("HEI2015C3_TOTALFRUIT", "HEI2015_TOTALFRT")),
    HEI2015_FRT = list(data1 = HEI2015_ASA24_NCI_SAS, data2 = HEI2015_ASA24_dietaryindex, cols = c("HEI2015C4_WHOLEFRUIT", "HEI2015_FRT")),
    HEI2015_VEG = list(data1 = HEI2015_ASA24_NCI_SAS, data2 = HEI2015_ASA24_dietaryindex, cols = c("HEI2015C1_TOTALVEG", "HEI2015_VEG")),
    HEI2015_GREENNBEAN = list(data1 = HEI2015_ASA24_NCI_SAS, data2 = HEI2015_ASA24_dietaryindex, cols = c("HEI2015C2_GREEN_AND_BEAN", "HEI2015_GREENNBEAN")),
    HEI2015_TOTALPRO = list(data1 = HEI2015_ASA24_NCI_SAS, data2 = HEI2015_ASA24_dietaryindex, cols = c("HEI2015C7_TOTPROT", "HEI2015_TOTALPRO")),
    HEI2015_SEAPLANTPRO = list(data1 = HEI2015_ASA24_NCI_SAS, data2 = HEI2015_ASA24_dietaryindex, cols = c("HEI2015C8_SEAPLANT_PROT", "HEI2015_SEAPLANTPRO")),
    HEI2015_WHOLEGRAIN = list(data1 = HEI2015_ASA24_NCI_SAS, data2 = HEI2015_ASA24_dietaryindex, cols = c("HEI2015C5_WHOLEGRAIN", "HEI2015_WHOLEGRAIN")),
    HEI2015_DAIRY = list(data1 = HEI2015_ASA24_NCI_SAS, data2 = HEI2015_ASA24_dietaryindex, cols = c("HEI2015C6_TOTALDAIRY", "HEI2015_DAIRY")),
    HEI2015_FATTYACID = list(data1 = HEI2015_ASA24_NCI_SAS, data2 = HEI2015_ASA24_dietaryindex, cols = c("HEI2015C9_FATTYACID", "HEI2015_FATTYACID")),
    HEI2015_REFINEDGRAIN = list(data1 = HEI2015_ASA24_NCI_SAS, data2 = HEI2015_ASA24_dietaryindex, cols = c("HEI2015C11_REFINEDGRAIN", "HEI2015_REFINEDGRAIN")),
    HEI2015_SODIUM = list(data1 = HEI2015_ASA24_NCI_SAS, data2 = HEI2015_ASA24_dietaryindex, cols = c("HEI2015C10_SODIUM", "HEI2015_SODIUM")),
    HEI2015_ADDEDSUGAR = list(data1 = HEI2015_ASA24_NCI_SAS, data2 = HEI2015_ASA24_dietaryindex, cols = c("HEI2015C13_ADDSUG", "HEI2015_ADDEDSUGAR")),
    HEI2015_SATFAT = list(data1 = HEI2015_ASA24_NCI_SAS, data2 = HEI2015_ASA24_dietaryindex, cols = c("HEI2015C12_SFAT", "HEI2015_SATFAT"))
    )

# Compute accuracy for each dataset in PHDI
for (name in names(datasets_HEI2015_ASA24)) {
  dataset <- datasets_HEI2015_ASA24[[name]]
  data1_values <- as.numeric(dataset$data1[[dataset$cols[1]]])  # Convert to numeric
  data2_values <- as.numeric(dataset$data2[[dataset$cols[2]]])  # Convert to numeric

  accuracy <- get_accuracy(data1_values, data2_values)
  results_HEI2015_ASA24 <- rbind(results_HEI2015_ASA24, data.frame(Component = name, Accuracy = accuracy))
}

# Print results
print(results_HEI2015_ASA24)

# Plot results
ggplot(results_HEI2015_ASA24, aes(x = Component, y = Accuracy, fill = Component)) +
  geom_bar(stat = "identity") +
  ylab("Accuracy (%)") +
  xlab(NULL) +
  ggtitle("Accuracy of HEI2015 in ASA24: dietaryindex-calculated vs. SAS-calculated Dietary Index Values from NCI") +
  # add a subtitie
  labs(subtitle = "Exact match by rounding both results to the nearest two decimal places") +
  # increase the title size
  theme(
    plot.title = element_text(size = 20),
    plot.subtitle = element_text(size = 15),
    axis.title = element_text(size = 20),
    axis.text = element_text(size = 15),
    legend.text = element_text(size = 15),
    legend.title = element_text(size = 20),
    axis.text.x = element_blank()
  )

############### HEI2015 validation in DHQ3 using dietaryindex-calculated results vs. National Cancer Institute (NCI) SAS results ######################
# Initialize a data frame to store results for all HEI2015_DHQ3 results
results_HEI2015_DHQ3 <- data.frame(
  Dataset = character(),
  Accuracy = numeric(),
  stringsAsFactors = FALSE
)

colnames(HEI2015_DHQ3_NCI_SAS)
colnames(HEI2015_DHQ3_dietaryindex)

# Create a list with HEI2015_DHQ3_NCI_SAS and HEI2015_DHQ3_dietaryindex
datasets_HEI2015_DHQ3<- list(
    HEI2015_ALL = list(data1 = HEI2015_DHQ3_NCI_SAS, data2 = HEI2015_DHQ3_dietaryindex, cols = c("Total HEI-2015 Score", "HEI2015_ALL")),
    HEI2015_TOTALFRT = list(data1 = HEI2015_DHQ3_NCI_SAS, data2 = HEI2015_DHQ3_dietaryindex, cols = c("HEI-2015 - Total Fruits - Component Score", "HEI2015_TOTALFRT")),
    HEI2015_FRT = list(data1 = HEI2015_DHQ3_NCI_SAS, data2 = HEI2015_DHQ3_dietaryindex, cols = c("HEI-2015 - Whole Fruits - Component Score", "HEI2015_FRT")),
    HEI2015_VEG = list(data1 = HEI2015_DHQ3_NCI_SAS, data2 = HEI2015_DHQ3_dietaryindex, cols = c("HEI-2015 - Total Vegetables - Component Score", "HEI2015_VEG")),
    HEI2015_GREENNBEAN = list(data1 = HEI2015_DHQ3_NCI_SAS, data2 = HEI2015_DHQ3_dietaryindex, cols = c("HEI-2015 - Greens and Beans - Component Score", "HEI2015_GREENNBEAN")),
    HEI2015_TOTALPRO = list(data1 = HEI2015_DHQ3_NCI_SAS, data2 = HEI2015_DHQ3_dietaryindex, cols = c("HEI-2015 - Total Protein Foods - Component Score", "HEI2015_TOTALPRO")),
    HEI2015_SEAPLANTPRO = list(data1 = HEI2015_DHQ3_NCI_SAS, data2 = HEI2015_DHQ3_dietaryindex, cols = c("HEI-2015 - Seafood and Plant Proteins - Component Score", "HEI2015_SEAPLANTPRO")),
    HEI2015_WHOLEGRAIN = list(data1 = HEI2015_DHQ3_NCI_SAS, data2 = HEI2015_DHQ3_dietaryindex, cols = c("HEI-2015 - Whole Grains - Component Score", "HEI2015_WHOLEGRAIN")),
    HEI2015_DAIRY = list(data1 = HEI2015_DHQ3_NCI_SAS, data2 = HEI2015_DHQ3_dietaryindex, cols = c("HEI-2015 - Dairy - Component Score", "HEI2015_DAIRY")),
    HEI2015_FATTYACID = list(data1 = HEI2015_DHQ3_NCI_SAS, data2 = HEI2015_DHQ3_dietaryindex, cols = c("HEI-2015 - Fatty Acids - Component Score", "HEI2015_FATTYACID")),
    HEI2015_REFINEDGRAIN = list(data1 = HEI2015_DHQ3_NCI_SAS, data2 = HEI2015_DHQ3_dietaryindex, cols = c("HEI-2015 - Refined Grains - Component Score", "HEI2015_REFINEDGRAIN")),
    HEI2015_SODIUM = list(data1 = HEI2015_DHQ3_NCI_SAS, data2 = HEI2015_DHQ3_dietaryindex, cols = c("HEI-2015 - Sodium - Component Score", "HEI2015_SODIUM")),
    HEI2015_ADDEDSUGAR = list(data1 = HEI2015_DHQ3_NCI_SAS, data2 = HEI2015_DHQ3_dietaryindex, cols = c("HEI-2015 - Added Sugars - Component Score", "HEI2015_ADDEDSUGAR")),
    HEI2015_SATFAT = list(data1 = HEI2015_DHQ3_NCI_SAS, data2 = HEI2015_DHQ3_dietaryindex, cols = c("HEI-2015 - Saturated Fats - Component Score", "HEI2015_SATFAT"))
    )

# Compute accuracy for each dataset in PHDI
for (name in names(datasets_HEI2015_DHQ3)) {
  dataset <- datasets_HEI2015_DHQ3[[name]]
  data1_values <- as.numeric(dataset$data1[[dataset$cols[1]]])  # Convert to numeric
  data2_values <- as.numeric(dataset$data2[[dataset$cols[2]]])  # Convert to numeric

  accuracy <- get_accuracy_diff(data1_values, data2_values)
  results_HEI2015_DHQ3 <- rbind(results_HEI2015_DHQ3, data.frame(Component = name, Accuracy = accuracy))
}

# Print results
print(results_HEI2015_DHQ3)

# Plot results
ggplot(results_HEI2015_DHQ3, aes(x = Component, y = Accuracy, fill = Component)) +
  geom_bar(stat = "identity") +
  ylab("Accuracy (%)") +
  xlab(NULL) +
  ggtitle("Accuracy of HEI2015 in DHQ3: dietaryindex-calculated vs. internal-calculated Dietary Index Values from NCI") +
  # add a subtitie
  labs(subtitle = "Check if the differences between 2 dietary indexes are within 0.5") +
  # increase the title size
  theme(
    plot.title = element_text(size = 20),
    plot.subtitle = element_text(size = 15),
    axis.title = element_text(size = 20),
    axis.text = element_text(size = 15),
    legend.text = element_text(size = 15),
    legend.title = element_text(size = 20),
    axis.text.x = element_blank()
  )
```
