## Supplementary figures and images for "Dietaryindex: A User-Friendly and Versatile R Package for Standardizing Dietary Pattern Analysis in Epidemiological and Clinical Studies"

### Supplementary Figure 1. ACS2020_V1 Accuracy.png

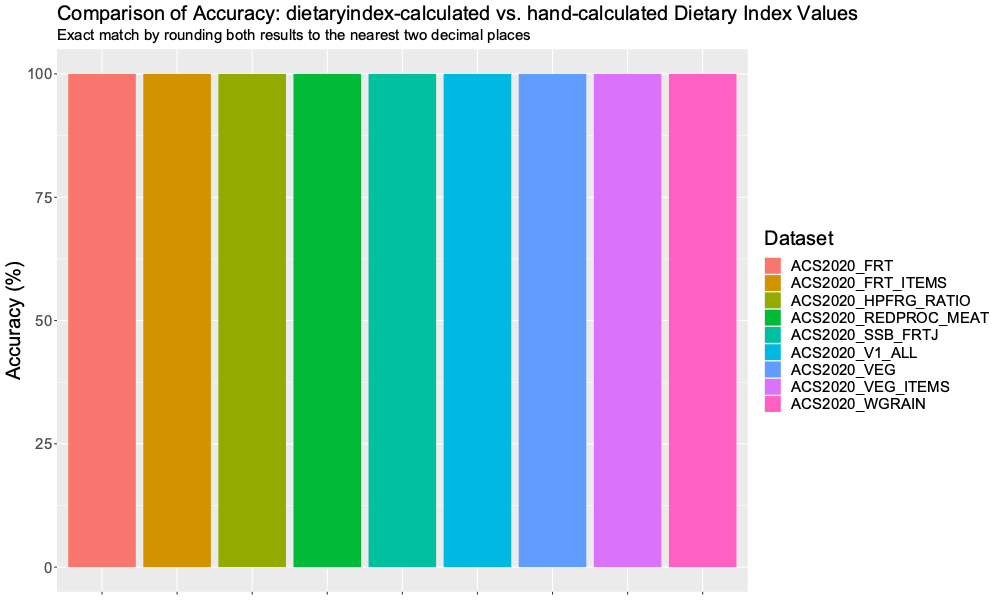

### Supplementary Figure 2. ACS2020_V2 Accuracy.png

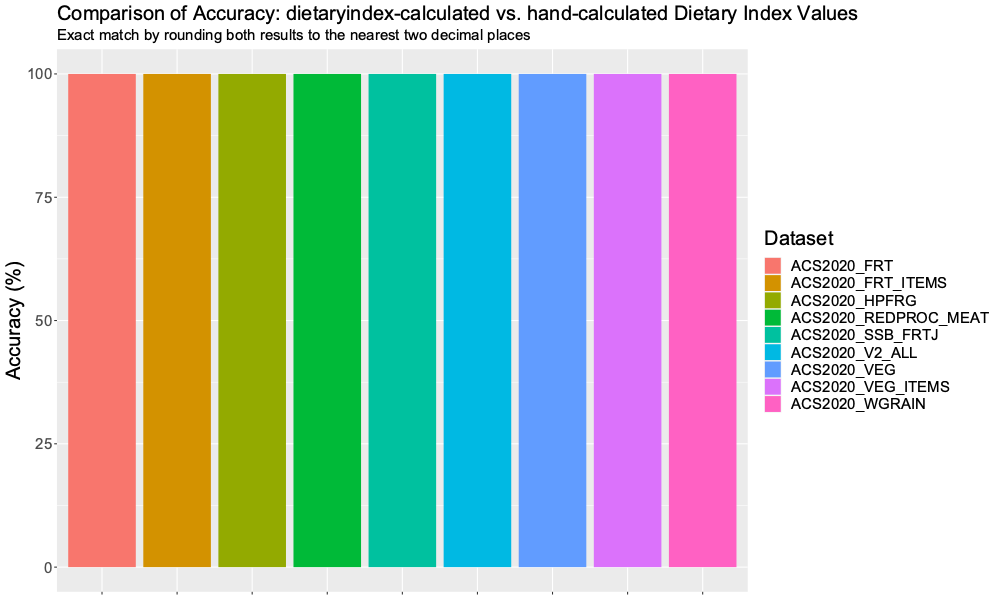

### Supplementary Figure 3. AHEI Accuracy.png

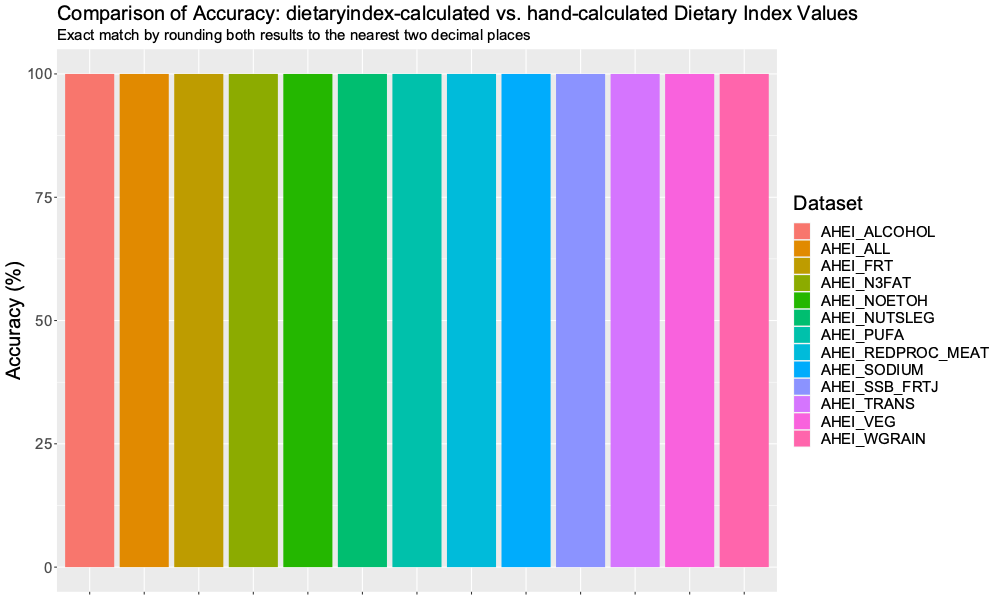

### Supplementary Figure 4. AHEIP Accuracy.png

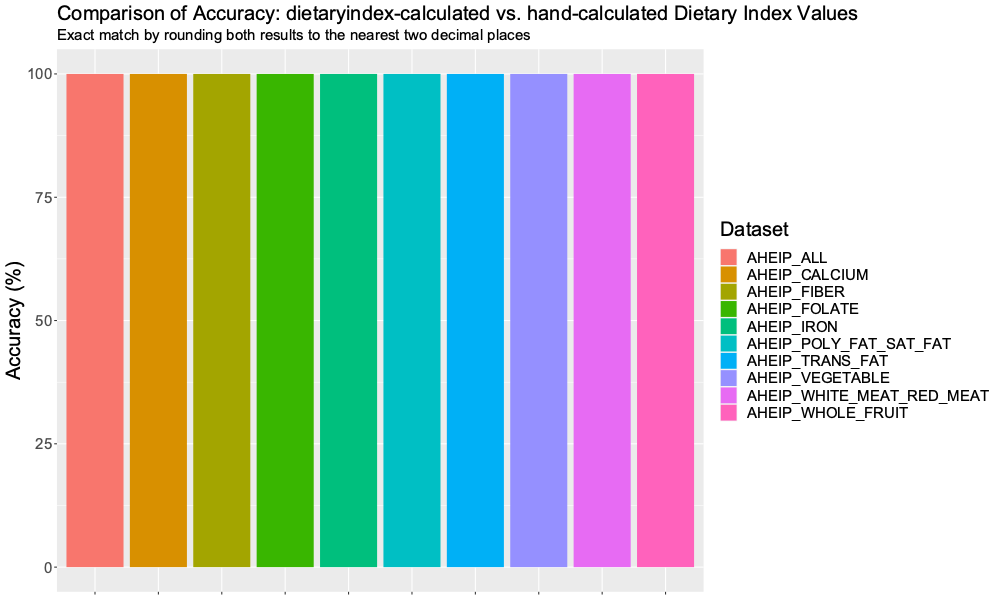

### Supplementary Figure 5. DASH Accuracy.png

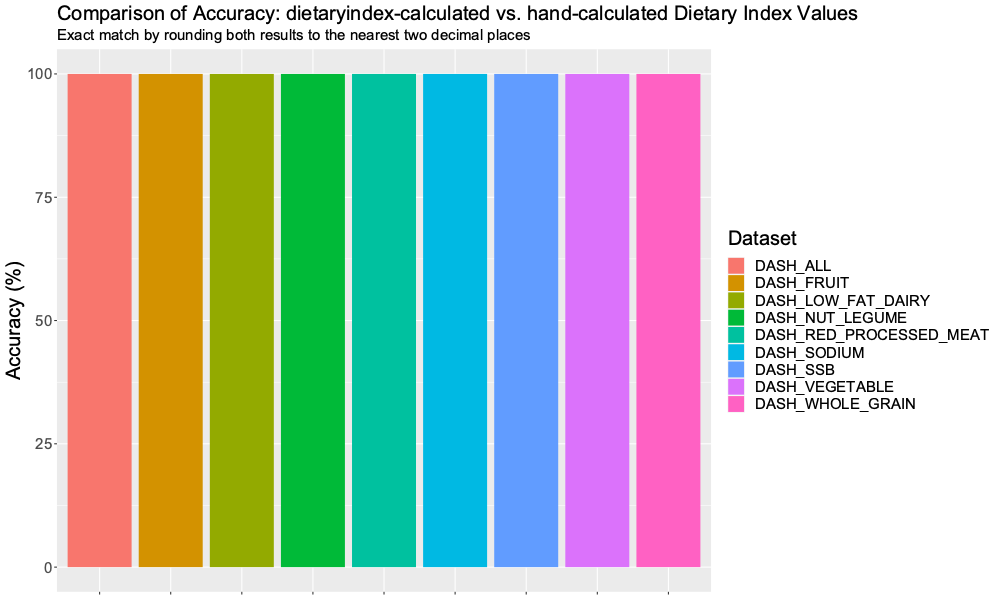

### Supplementary Figure 6. DASHI Accuracy.png

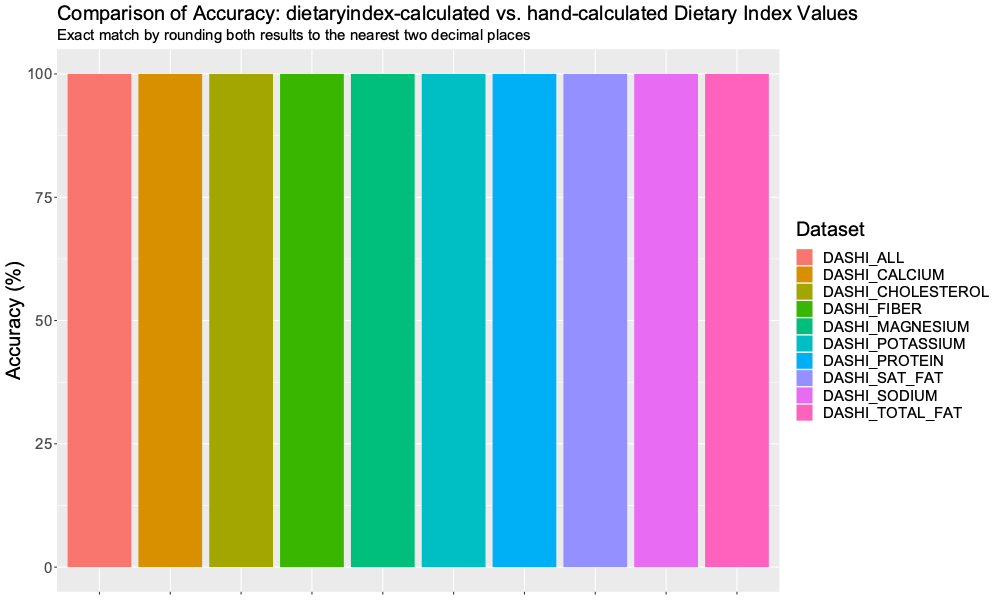

### Supplementary Figure 7. DII Accuracy.png

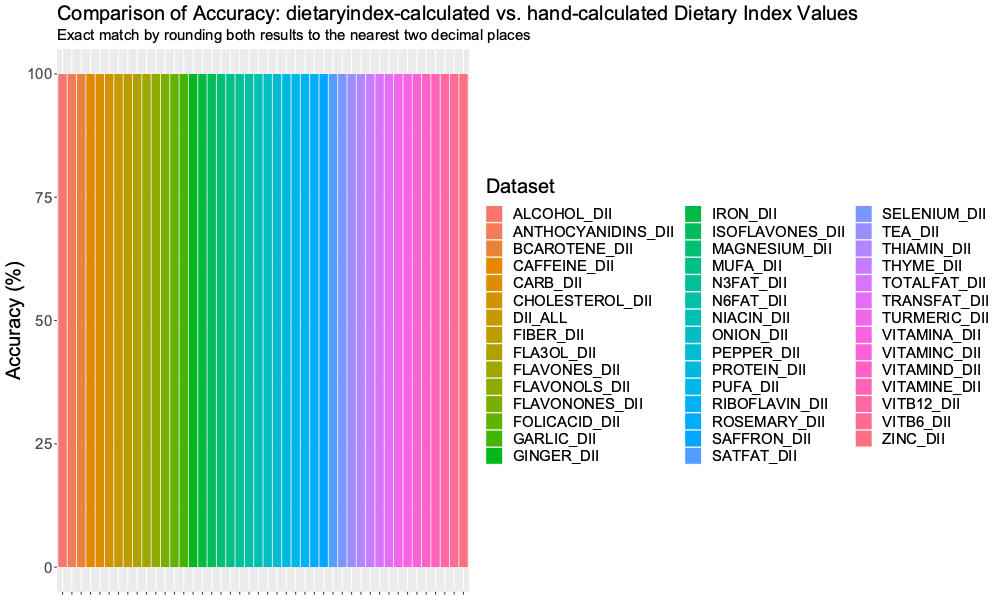

### Supplementary Figure 8. HEI2015 Accuracy.png

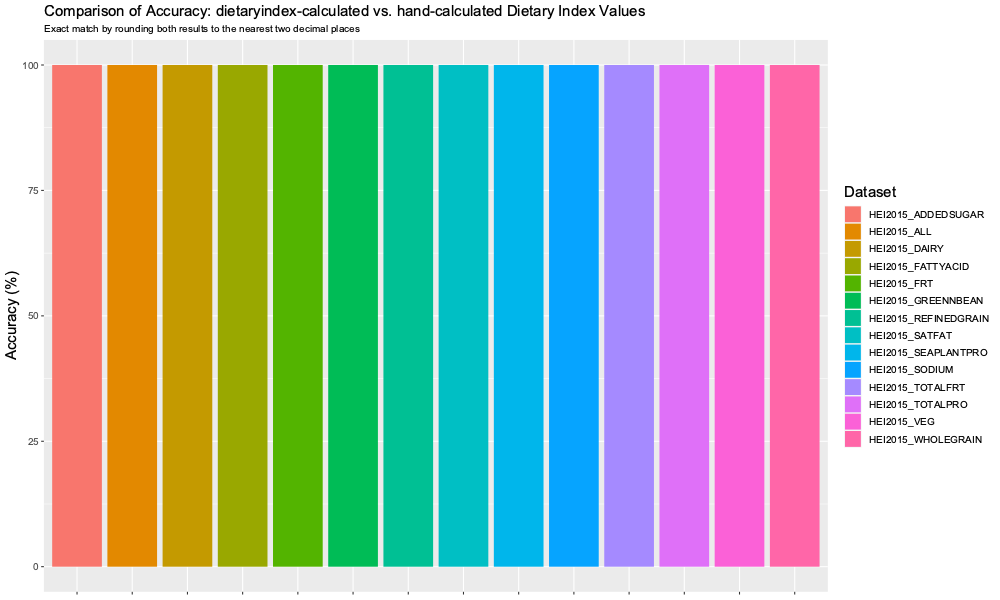

### Supplementary Figure 9. HEI2020 Accuracy.png

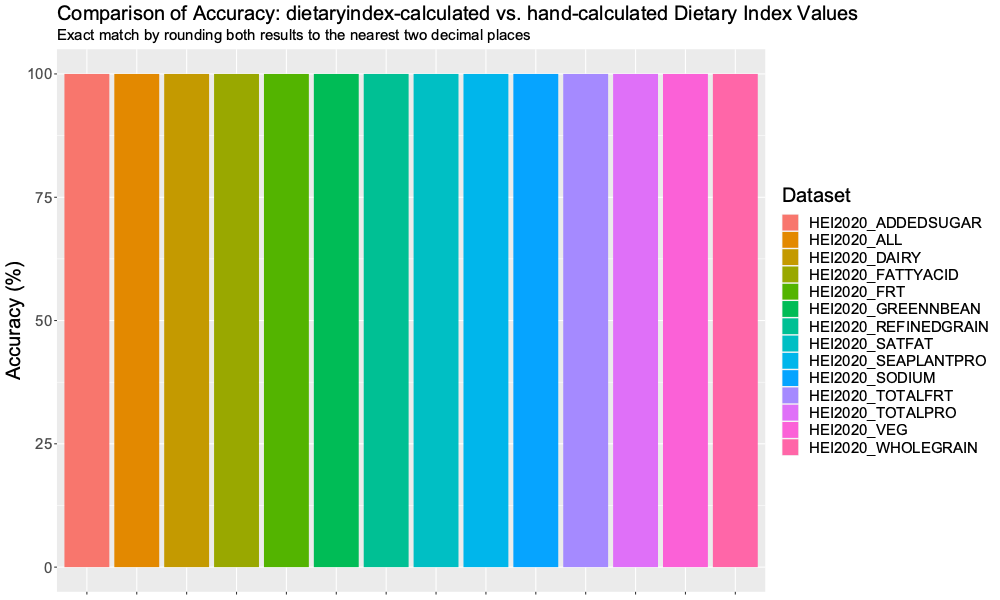

### Supplementary Figure 10. MED Accuracy.png

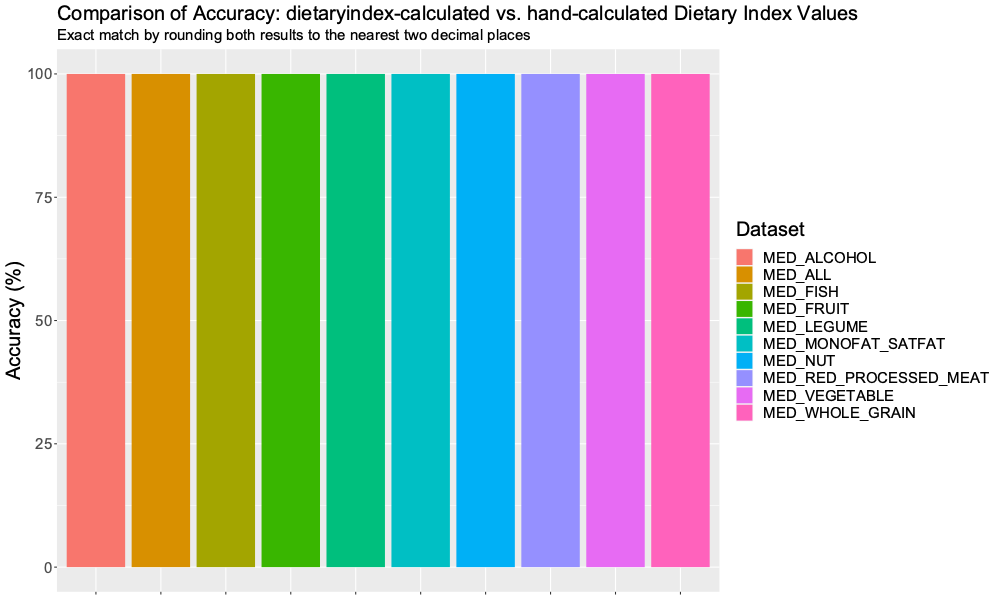

### Supplementary Figure 11. MEDI Accuracy.png

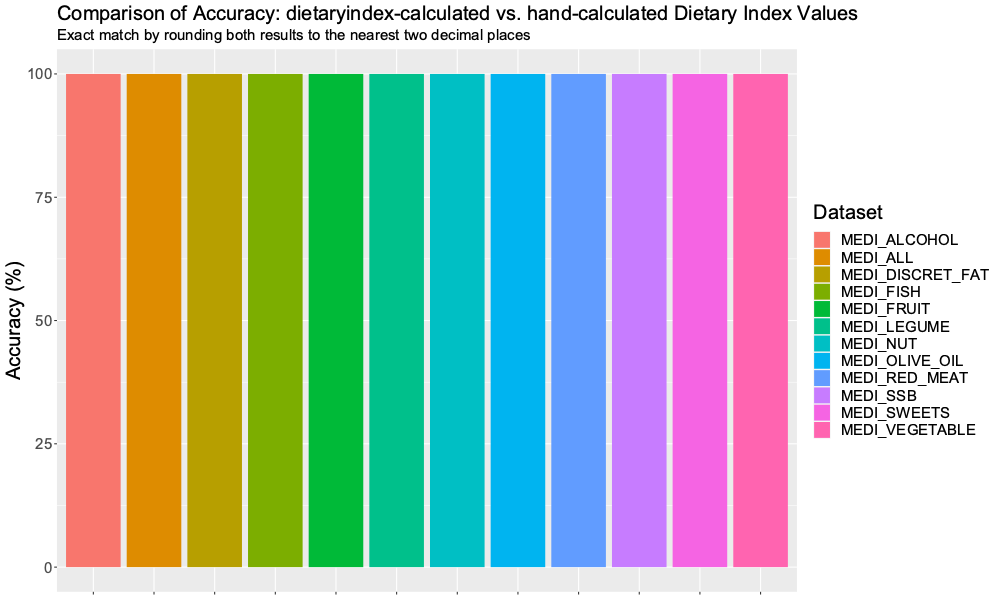

### Supplementary Figure 12. MEDI_V2 Accuracy.png

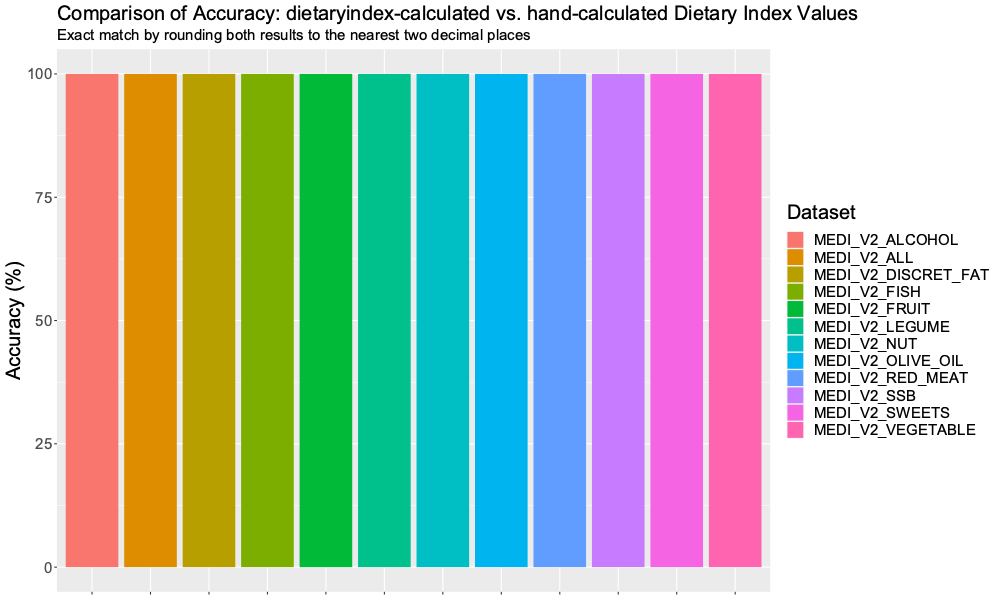

### Supplementary Figure 13. PHDI Accuracy.png

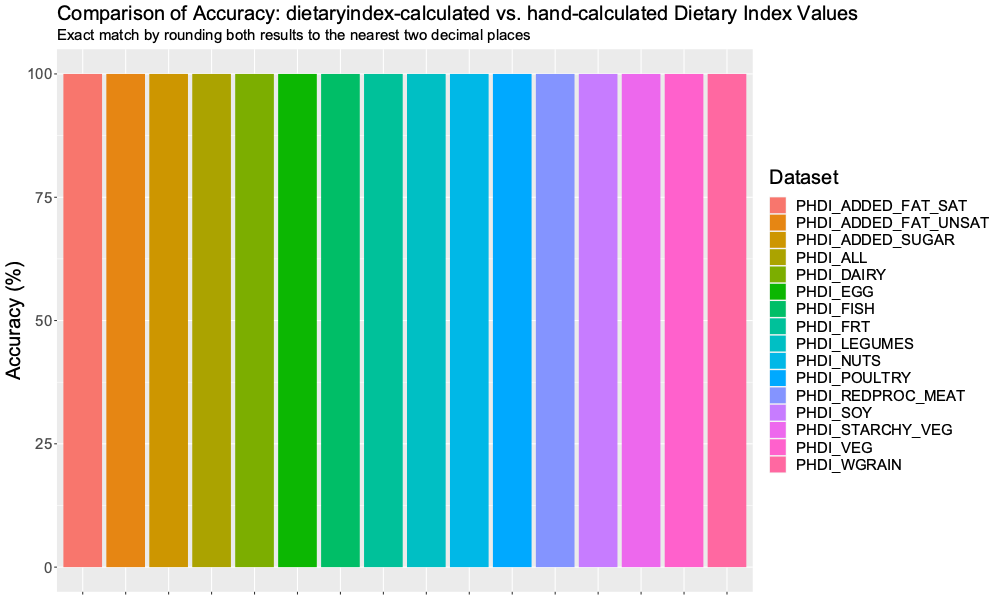
