## Supplement 4 R scripts for case studies for "Dietaryindex: A User-Friendly and Versatile R Package for Standardizing Dietary Pattern Analysis in Epidemiological and Clinical Studies": Case study 1,2,3.r

```
# remove memory
rm(list=ls())

install.packages("devtools") #If you don't have "devtools" installed already
devtools::install_github("jamesjiadazhan/dietaryindex") # Install the package from GitHub

# remove utils from library if you want to reload the package
install.packages("/Users/james/Downloads/dietaryindex-main", repos = NULL, type = "source")

# If the previous steps not working, you can try the following steps by removing the "#" marks
# library(devtools) # Load devtools
# install_github("jamesjiadazhan/dietaryindex")

library(ggplot2)
library(dplyr)
# library(magrittr)
library(tidyr)

library(dietaryindex)
# remove dietaryindex from library if you want to reload the package
# detach("package:dietaryindex", unload=TRUE)

# Load the data
data("DASH_trial")
data("PREDIMED_trial")
data("NHANES_20172018")

# set up working dictionary
setwd("/Users/james/Desktop/NHANES_combined")

# Load the NHANES data from 2005 to 2018
## NHANES 2005-2006
load("NHANES_20052006.rda")

## NHANES 2007-2008
load("NHANES_20072008.rda")

## NHANES 2009-2010
load("NHANES_20092010.rda")

## NHANES 2011-2012
load("NHANES_20112012.rda")

## NHANES 2013-2014
load("NHANES_20132014.rda")

## NHANES 2015-2016
load("NHANES_20152016.rda")

# DASHI from the DASH trial
DASHI_DASH = DASHI(
    SERV_DATA = DASH_trial, 
    RESPONDENTID = DASH_trial$Diet_Type,
    TOTALKCAL_DASHI = DASH_trial$Kcal, 
    TOTAL_FAT_DASHI = DASH_trial$Totalfat_Percent, 
    SAT_FAT_DASHI = DASH_trial$Satfat_Percent, 
    PROTEIN_DASHI = DASH_trial$Protein_Percent, 
    CHOLESTEROL_DASHI = DASH_trial$Cholesterol, 
    FIBER_DASHI = DASH_trial$Fiber, 
    POTASSIUM_DASHI = DASH_trial$Potassium, 
    MAGNESIUM_DASHI = DASH_trial$Magnesium, 
    CALCIUM_DASHI = DASH_trial$Calcium, 
    SODIUM_DASHI = DASH_trial$Sodium)

# MEDI for the PREDIMED trial
MEDI_PREDIMED = MEDI(
  SERV_DATA = PREDIMED_trial,
  RESPONDENTID = PREDIMED_trial$Diet_Type,
  OLIVE_OIL_SERV_MEDI = PREDIMED_trial$Virgin_Oliveoil,
  FRT_SERV_MEDI = PREDIMED_trial$Fruits, 
  VEG_SERV_MEDI = PREDIMED_trial$Vegetables,
  LEGUMES_SERV_MEDI = PREDIMED_trial$Legumes,
  NUTS_SERV_MEDI = PREDIMED_trial$Total_nuts,
  FISH_SEAFOOD_SERV_MEDI = PREDIMED_trial$Fish_Seafood,
  ALCOHOL_SERV_MEDI = PREDIMED_trial$Alcohol,
  SSB_SERV_MEDI = PREDIMED_trial$Soda_Drinks,
  SWEETS_SERV_MEDI = PREDIMED_trial$Sweets,
  DISCRET_FAT_SERV_MEDI = PREDIMED_trial$Refined_Oliveoil,
  REDPROC_MEAT_SERV_MEDI = PREDIMED_trial$Meat)


## NHANES 2017-2018
# DASHI for day 1 and day 2
DASHI_NHANES = DASHI_NHANES_FPED(NHANES_20172018$NUTRIENT, NHANES_20172018$NUTRIENT2)

# MEDI for day 1 and day 2
MEDI_NHANES = MEDI_NHANES_FPED(FPED_IND_PATH=NHANES_20172018$FPED_IND, NUTRIENT_IND_PATH=NHANES_20172018$NUTRIENT_IND, FPED_IND_PATH2=NHANES_20172018$FPED_IND2, NUTRIENT_IND_PATH2=NHANES_20172018$NUTRIENT_IND2)

# DASH for day 1 and day 2
DASH_NHANES = DASH_NHANES_FPED(NHANES_20172018$FPED_IND, NHANES_20172018$NUTRIENT_IND, NHANES_20172018$FPED_IND2, NHANES_20172018$NUTRIENT_IND2)

# MED for day 1 and day 2
MED_NHANES = MED_NHANES_FPED(FPED_PATH=NHANES_20172018$FPED, NUTRIENT_PATH=NHANES_20172018$NUTRIENT, DEMO_PATH=NHANES_20172018$DEMO, FPED_PATH2=NHANES_20172018$FPED2, NUTRIENT_PATH2=NHANES_20172018$NUTRIENT2)

# AHEI for day 1 and day 2
AHEI_NHANES = AHEI_NHANES_FPED(NHANES_20172018$FPED_IND, NHANES_20172018$NUTRIENT_IND, NHANES_20172018$FPED_IND2, NHANES_20172018$NUTRIENT_IND2)

# DII for day 1 and day 2
DII_NHANES = DII_NHANES_FPED(FPED_PATH=NHANES_20172018$FPED, NUTRIENT_PATH=NHANES_20172018$NUTRIENT, DEMO_PATH=NHANES_20172018$DEMO, FPED_PATH2=NHANES_20172018$FPED2, NUTRIENT_PATH2=NHANES_20172018$NUTRIENT2)

# HEI2020 for day 1 and day 2
## 2017-2018
HEI2020_NHANES_1718 = HEI2020_NHANES_FPED(FPED_PATH=NHANES_20172018$FPED, NUTRIENT_PATH=NHANES_20172018$NUTRIENT, DEMO_PATH=NHANES_20172018$DEMO, FPED_PATH2=NHANES_20172018$FPED2, NUTRIENT_PATH2=NHANES_20172018$NUTRIENT2)

# HEI2020 for day 1
## 2017-2018
HEI2020_NHANES_1718_d1 = HEI2020_NHANES_FPED(FPED_PATH=NHANES_20172018$FPED, NUTRIENT_PATH=NHANES_20172018$NUTRIENT, DEMO_PATH=NHANES_20172018$DEMO)
### subset for toddlers
HEI2020_NHANES_1718_d1_toddler = HEI2020_NHANES_1718_d1 %>%
    dplyr::filter(RIDAGEYR < 2)
### subset for non-toddlers (children and adults)
HEI2020_NHANES_1718_d1_nontoddler = HEI2020_NHANES_1718_d1 %>%
    dplyr::filter(RIDAGEYR >= 2)

## 2015-2016
HEI2020_NHANES_1516_d1 = HEI2020_NHANES_FPED(FPED_PATH=NHANES_20152016$FPED, NUTRIENT_PATH=NHANES_20152016$NUTRIENT, DEMO_PATH=NHANES_20152016$DEMO)
### subset for toddlers
HEI2020_NHANES_1516_d1_toddler = HEI2020_NHANES_1516_d1 %>%
    dplyr::filter(RIDAGEYR < 2)
### subset for non-toddlers (children and adults)
HEI2020_NHANES_1516_d1_nontoddler = HEI2020_NHANES_1516_d1 %>%
    dplyr::filter(RIDAGEYR >= 2)
    
## 2013-2014
HEI2020_NHANES_1314_d1 = HEI2020_NHANES_FPED(FPED_PATH=NHANES_20132014$FPED, NUTRIENT_PATH=NHANES_20132014$NUTRIENT, DEMO_PATH=NHANES_20132014$DEMO)
### subset for toddlers
HEI2020_NHANES_1314_d1_toddler = HEI2020_NHANES_1314_d1 %>%
    dplyr::filter(RIDAGEYR < 2)
### subset for non-toddlers (children and adults)
HEI2020_NHANES_1314_d1_nontoddler = HEI2020_NHANES_1314_d1 %>%
    dplyr::filter(RIDAGEYR >= 2)

## 2011-2012
HEI2020_NHANES_1112_d1 = HEI2020_NHANES_FPED(FPED_PATH=NHANES_20112012$FPED, NUTRIENT_PATH=NHANES_20112012$NUTRIENT, DEMO_PATH=NHANES_20112012$DEMO)
### subset for toddlers
HEI2020_NHANES_1112_d1_toddler = HEI2020_NHANES_1112_d1 %>%
    dplyr::filter(RIDAGEYR < 2)
### subset for non-toddlers (children and adults)
HEI2020_NHANES_1112_d1_nontoddler = HEI2020_NHANES_1112_d1 %>%
    dplyr::filter(RIDAGEYR >= 2)

## 2009-2010
HEI2020_NHANES_0910_d1 = HEI2020_NHANES_FPED(FPED_PATH=NHANES_20092010$FPED, NUTRIENT_PATH=NHANES_20092010$NUTRIENT, DEMO_PATH=NHANES_20092010$DEMO)
### subset for toddlers
HEI2020_NHANES_0910_d1_toddler = HEI2020_NHANES_0910_d1 %>%
    dplyr::filter(RIDAGEYR < 2)
### subset for non-toddlers (children and adults)
HEI2020_NHANES_0910_d1_nontoddler = HEI2020_NHANES_0910_d1 %>%
    dplyr::filter(RIDAGEYR >= 2)

## 2007-2008
HEI2020_NHANES_0708_d1 = HEI2020_NHANES_FPED(FPED_PATH=NHANES_20072008$FPED, NUTRIENT_PATH=NHANES_20072008$NUTRIENT, DEMO_PATH=NHANES_20072008$DEMO)
### subset for toddlers
HEI2020_NHANES_0708_d1_toddler = HEI2020_NHANES_0708_d1 %>%
    dplyr::filter(RIDAGEYR < 2)
### subset for non-toddlers (children and adults)
HEI2020_NHANES_0708_d1_nontoddler = HEI2020_NHANES_0708_d1 %>%
    dplyr::filter(RIDAGEYR >= 2)

## 2005-2006
HEI2020_NHANES_0506_d1 = HEI2020_NHANES_FPED(FPED_PATH=NHANES_20052006$FPED, NUTRIENT_PATH=NHANES_20052006$NUTRIENT, DEMO_PATH=NHANES_20052006$DEMO)
### subset for toddlers
HEI2020_NHANES_0506_d1_toddler = HEI2020_NHANES_0506_d1 %>%
    dplyr::filter(RIDAGEYR < 2)
### subset for non-toddlers (children and adults)
HEI2020_NHANES_0506_d1_nontoddler = HEI2020_NHANES_0506_d1 %>%
    dplyr::filter(RIDAGEYR >= 2)

# Case study 1: A comparative analysis of results derived from clinical trials (i.e., DASH and PREDIMED) juxtaposed with findings from an epidemiological study (i.e., NHANES) from 2017-2018, utilizing DASHI and MEDI dietary indexes
## Create a vector of dietary index names for DASHI and MEDI in the DASH and PREDIMED trials and NHANES 2017-2018
x_case1 = c("DASHI_DASH_DASHlowSodium", "DASHI_DASH_DASHMedSodium", "DASHI_DASH_Control", "MEDI_PREDIMED_Med_Oliveoil", "MEDI_PREDIMED_Med_Nuts", "MEDI_PREDIMED_Control", "DASHI_NHANES", "MEDI_NHANES")
## Create a vector of mean dietary index percentile for DASHI and MEDI in the DASH and PREDIMED trials and NHANES 2017-2018
y_case1 = c(DASHI_DASH$DASHI_ALL[2]/9, DASHI_DASH$DASHI_ALL[3]/9, DASHI_DASH$DASHI_ALL[5]/9, MEDI_PREDIMED$MEDI_ALL[1]/11,  MEDI_PREDIMED$MEDI_ALL[2]/11, MEDI_PREDIMED$MEDI_ALL[3]/11, mean(DASHI_NHANES$DASHI_ALL, na.rm=TRUE)/9, mean(MEDI_NHANES$MEDI_ALL, na.rm=TRUE)/11)
## Create a vector of data source
z_case1 = c("DASH_trial", "DASH_trial", "DASH_trial", "PREDIMED_trial", "PREDIMED_trial", "PREDIMED_trial", "NHANES_20172018", "NHANES_20172018")
## Create a vector of the dietary index type
w_case1 = c("DASHI", "DASHI", "DASHI", "MEDI", "MEDI", "MEDI", "DASHI", "MEDI")

# multiply all y values by 100 to get the percentage
y_case1 = y_case1*100

## create a data frame
df_case1 = data.frame(x_case1, y_case1, z_case1, w_case1)

# reorder the z_case1 factor levels so taht the order of NHANES 2017-2018 is always on the left side
df_case1$z_case1 = factor(df_case1$z_case1, levels = c("NHANES_20172018", "DASH_trial", "PREDIMED_trial"))

# reorder the x_case1 factor levels
df_case1$x_case1 = factor(df_case1$x_case1, levels = c("DASHI_NHANES", "DASHI_DASH_Control", "DASHI_DASH_DASHMedSodium", "DASHI_DASH_DASHlowSodium", "MEDI_NHANES", "MEDI_PREDIMED_Control", "MEDI_PREDIMED_Med_Nuts", "MEDI_PREDIMED_Med_Oliveoil"))


## plot the data with histogram with different colors for different x values
ggplot(df_case1, aes(x=z_case1, y=y_case1, fill=x_case1)) + 
  geom_bar(stat = "identity", position = "dodge") +
  theme_bw() +
  # create a facet grid with the dietary index type
  facet_wrap(. ~ w_case1, scales = "free_x") +
  labs(
    x = "Data Source", 
    y = "Mean dietary index percentile", 
    title = "DASHI and MEDI Dietary Indexes in the DASH and PREDIMED trials and NHANES 2017-2018", 
    fill = "Dieary indexes for specific diets"
    ) +
  theme(
    # increase the plot title size
    plot.title = element_text(size=18),
    # remove the x axis title
    axis.title.x = element_blank(),
    # increase the x axis text size and rotate the text by 90 degrees
    axis.text.x = element_text(angle = 45, hjust = 1, size = 14),
    # increase the y axis title and text size
    axis.title.y = element_text(size=18),
    axis.text.y = element_text(size=14),
    # increase the legend title and text size
    legend.title = element_text(size=18),
    legend.text = element_text(size=16),
    # increase the facet label size
    strip.text = element_text(size = 16)
  ) +
  # add numeric labels to the bars and increase their size
  geom_text(aes(label = round(y_case1, 2)), vjust = -0.5, size = 4.5, position = position_dodge(0.9)) +
  # add custom fill labels
  scale_fill_discrete(labels = c(
    "DASHI_DASH_DASHlowSodium" = "DASHI for DASH trial low sodium diet",
    "DASHI_DASH_DASHMedSodium" = "DASHI for DASH trial medium sodium diet",
    "DASHI_DASH_Control" = "DASHI for DASH trial control diet",
    "MEDI_PREDIMED_Med_Oliveoil" = "MEDI for PREDIMED mediterranean olive oil diet",
    "MEDI_PREDIMED_Med_Nuts" = "MEDI for PREDIMED mediterranean nuts diet",
    "MEDI_PREDIMED_Control" = "MEDI for PREDIMED control diet",
    "DASHI_NHANES" = "DASHI for NHANES 2017-18",
    "MEDI_NHANES" = "MEDI for NHANES 2017-18"
  ))


# Case study 2: A time series of cross-sectional computation of the HEI2020 from 2005 to 2018
x_case2 = c("2005-2006", "2007-2008", "2009-2010", "2011-2012", "2013-2014", "2015-2016", "2017-2018")
# mean for toddlers
y1_case2 = c(mean(HEI2020_NHANES_0506_d1_toddler$HEI2020_ALL, na.rm=TRUE), mean(HEI2020_NHANES_0708_d1_toddler$HEI2020_ALL, na.rm=TRUE), mean(HEI2020_NHANES_0910_d1_toddler$HEI2020_ALL, na.rm=TRUE), mean(HEI2020_NHANES_1112_d1_toddler$HEI2020_ALL, na.rm=TRUE), mean(HEI2020_NHANES_1314_d1_toddler$HEI2020_ALL, na.rm=TRUE), mean(HEI2020_NHANES_1516_d1_toddler$HEI2020_ALL, na.rm=TRUE), mean(HEI2020_NHANES_1718_d1_toddler$HEI2020_ALL, na.rm=TRUE))
# mean for non-toddlers
y2_case2 = c(mean(HEI2020_NHANES_0506_d1_nontoddler$HEI2020_ALL, na.rm=TRUE), mean(HEI2020_NHANES_0708_d1_nontoddler$HEI2020_ALL, na.rm=TRUE), mean(HEI2020_NHANES_0910_d1_nontoddler$HEI2020_ALL, na.rm=TRUE), mean(HEI2020_NHANES_1112_d1_nontoddler$HEI2020_ALL, na.rm=TRUE), mean(HEI2020_NHANES_1314_d1_nontoddler$HEI2020_ALL, na.rm=TRUE), mean(HEI2020_NHANES_1516_d1_nontoddler$HEI2020_ALL, na.rm=TRUE), mean(HEI2020_NHANES_1718_d1_nontoddler$HEI2020_ALL, na.rm=TRUE))

# sd for toddlers
sd1_case2 = c(sd(HEI2020_NHANES_0506_d1_toddler$HEI2020_ALL, na.rm=TRUE), sd(HEI2020_NHANES_0708_d1_toddler$HEI2020_ALL, na.rm=TRUE), sd(HEI2020_NHANES_0910_d1_toddler$HEI2020_ALL, na.rm=TRUE), sd(HEI2020_NHANES_1112_d1_toddler$HEI2020_ALL, na.rm=TRUE), sd(HEI2020_NHANES_1314_d1_toddler$HEI2020_ALL, na.rm=TRUE), sd(HEI2020_NHANES_1516_d1_toddler$HEI2020_ALL, na.rm=TRUE), sd(HEI2020_NHANES_1718_d1_toddler$HEI2020_ALL, na.rm=TRUE))
# sd for non-toddlers
sd2_case2 = c(sd(HEI2020_NHANES_0506_d1_nontoddler$HEI2020_ALL, na.rm=TRUE), sd(HEI2020_NHANES_0708_d1_nontoddler$HEI2020_ALL, na.rm=TRUE), sd(HEI2020_NHANES_0910_d1_nontoddler$HEI2020_ALL, na.rm=TRUE), sd(HEI2020_NHANES_1112_d1_nontoddler$HEI2020_ALL, na.rm=TRUE), sd(HEI2020_NHANES_1314_d1_nontoddler$HEI2020_ALL, na.rm=TRUE), sd(HEI2020_NHANES_1516_d1_nontoddler$HEI2020_ALL, na.rm=TRUE), sd(HEI2020_NHANES_1718_d1_nontoddler$HEI2020_ALL, na.rm=TRUE))


## create a data frame
df_case2 = data.frame(x_case2, y1_case2, y2_case2, sd1_case2, sd2_case2)

# Convert data frame to long format
df_case2_long <- df_case2 %>%
  pivot_longer(cols = starts_with(c("y", "sd")),
               names_to = "group",
               values_to = "HEI2020")

# Adjust group labels
df_case2_long$group1 <- ifelse(df_case2_long$group %in% c("y1_case2", "sd1_case2"), "Toddler", "Non-toddler (children and adults)")

df_case2_long_mean_only = df_case2_long %>% filter(group %in% c("y1_case2", "y2_case2"))

# Plot data with mean only
ggplot(df_case2_long_mean_only, aes(x=x_case2, y=HEI2020, group=group1, color=group1)) +
  geom_line() +
  geom_point() +
  theme_bw() +
  labs(x = "NHANES Cycle", y = "Mean HEI2020", title = "Mean HEI2020 from 2005 to 2018", color = "Group") +
  theme(plot.title = element_text(size=22),
        axis.title.x = element_text(size=18),
        axis.title.y = element_text(size=18),
        axis.text.x = element_text(size=14, angle = 90, hjust = 1),
        legend.text = element_text(size=16),
        legend.title = element_text(size=18),
        aspect.ratio = 0.5) +
  geom_text(aes(label = round(HEI2020, 2)), vjust = -0.5, size = 4)

# # Separate data into mean and sd
# df_case2_long_mean <- df_case2_long %>% filter(group %in% c("y1_case2", "y2_case2"))
# df_case2_long_sd <- df_case2_long %>% filter(group %in% c("sd1_case2", "sd2_case2"))

# # Rename HEI2020 to Mean_HEI2020 and SD_HEI2020
# names(df_case2_long_mean)[names(df_case2_long_mean) == "HEI2020"] <- "Mean_HEI2020"
# names(df_case2_long_sd)[names(df_case2_long_sd) == "HEI2020"] <- "SD_HEI2020"

# # Merge the two data frames
# df_case2_long_2 <- merge(df_case2_long_mean, df_case2_long_sd, by = c("x_case2", "group1"))

# # Plot data with error bars 
# ggplot(df_case2_long_2, aes(x=x_case2, y=Mean_HEI2020, color=group1)) +
#   geom_line() +
#   geom_point() +
#   geom_errorbar(aes(ymin=Mean_HEI2020-SD_HEI2020, ymax=Mean_HEI2020+SD_HEI2020), width=0.2) +
#   theme_bw() +
#   labs(x = "NHANES Cycle", y = "Mean HEI2020", title = "Mean HEI2020 from 2005 to 2018", color = "Group") +
#   theme(plot.title = element_text(size=22),
#         axis.title.x = element_text(size=18),
#         axis.title.y = element_text(size=18),
#         axis.text.x = element_text(size=14, angle = 90, hjust = 1),
#         aspect.ratio = 0.5) +
#   geom_text(aes(label = round(Mean_HEI2020, 2)), vjust = -0.5, size = 3.5)
# # This plot is visually challenging to interpret, so we just plot the mean instead.


# Case study 3: Comprehensive Calculation of Multiple Dietary indexes, using the NHANES data in 2017-2018
## create 2 vectors for the x and y axis
x_case3 = c("DASHI", "MEDI", "DASH", "MED", "HEI2020", "AHEI", "DII")
y_case3 = c(mean(DASHI_NHANES$DASHI_ALL, na.rm=TRUE)/9, mean(MEDI_NHANES$MEDI_ALL, na.rm=TRUE)/11, mean(DASH_NHANES$DASH_ALL, na.rm=TRUE)/40, mean(MED_NHANES$MED_ALL, na.rm=TRUE)/9, mean(HEI2020_NHANES_1718$HEI2020_ALL, na.rm=TRUE)/100, mean(AHEI_NHANES$AHEI_ALL, na.rm=TRUE)/110, 0.75)
# multiply all y values by 100 to get the percentage
y_case3 = y_case3*100

## create a data frame
df_case3 = data.frame(x_case3, y_case3)

## plot the data with histogram with different colors for different x values
ggplot(df_case3, aes(x_case3, y_case3, fill=x_case3)) + 
  geom_bar(stat = "identity") + theme_bw() + 
  labs(x = "", y = "Mean dietary index percentile", title = "Dietary Indexes in NHANES 2017-2018", fill = "Dietary Index") + 
  theme(plot.title = element_text(size=22)) +
  theme(axis.title.x = element_blank()) +
  theme(axis.title.y = element_text(size=18)) +
  theme(axis.text.x = element_blank()) +
  theme(axis.text.y = element_text(size=14)) +
  # increase the size of the legend title
  theme(legend.title = element_text(size=18)) +
  # increase the size of the legend text
  theme(legend.text = element_text(size=16)) +
  theme(aspect.ratio = 0.5) +
  # add numeric labels to the bars
  geom_text(aes(label = round(y_case3, 2)), vjust = -0.5, size = 4.5)
```
